# A novel inhibitor against the bromodomain *Pf*BDP1 of the malaria pathogen *Plasmodium falciparum*

**DOI:** 10.1101/2025.01.16.633141

**Authors:** Marius Amann, Robin Warstat, Kay Kristin Rechten, Philip Theuer, Magdalena Schustereder, Sophie Clavey, Bernhard Breit, Oliver Einsle, Martin Huegle, Michaela Petter, Stefan Guenther

## Abstract

The rise of drug resistances in malaria necessitates the exploration of novel therapeutic strategies. Targeting epigenetic pathways could open new, promising treatment avenues. In this study, we focus on *Pf*BDP1, an essential bromodomain protein in *P. falciparum*. Utilizing the pan-selective bromodomain inhibitor MPM6, we identified a potent initial hit and subsequently developed it into a nanomolar binder. Through a combination of virtual docking, isothermal titration calorimetry, and X-ray crystallography, we elucidated the molecular interactions of the new inhibitors with *Pf*BD1. Our findings include the first cocrystallized structures of *Pf*BD1 and *Pv*BD1 with these inhibitors, providing insights into their binding mechanisms. Further validation using conditional knockdown of *Pf*BDP1 in *P. falciparum* demonstrated parasite sensitivity to the inhibitor, underscoring its potential as a targeted therapeutic approach against malaria.

## Results

Malaria, caused by the eukaryotic parasite *Plasmodium spec*., is a major public health issue, with approximately 249 million infected and 608,000 deaths worldwide reported in 2022^[1]^. The majority of casualties are attributed to *Plasmodium falciparum* (*P. falciparum*), the most lethal variant of the *Plasmodium* parasites. Despite intensive efforts to control the disease, little progress has been made in recent years towards reducing malaria burden and death; indeed, drug resistances have developed against the two most common drugs (artemisinin-based combination therapies and atovaquone–proguanil), highlighting the need for new therapeutic approaches^[2–4]^. Recent studies see this potential in the disruption of epigenetic pathways^[5–8]^. Potential targets are the nine predicted bromodomain (BD) proteins (BRDs), among which *Pf*BDP1 has been identified as essential in the blood stage of *P. falciparum*^[9–11]^. By conditional knockdown (KD), Josling *et al*. identified *Pf*BDP1 as an essential BRD for erythrocyte invasion, leading to a significantly reduced multiplication rate during the intraerythrocytic developmental cycle (IDC)^[12]^. It remains elusive whether this effect could also be induced by directly targeting the BD through small molecule inhibition. A recent structural and binding analysis of *Pf*BD1 revealed that a tetra-acetylated histone H4 peptide showed highest binding affinity; however, there is no cocrystallized structure with a natural ligand or inhibitor^[13]^. Within the present study, we address the effect of inhibiting the BD of *Pf*BDP1 (*Pf*BD1) using small molecule inhibitors based on the recently published human pan-selective BD inhibitor (BDi) MPM6^[14]^. We first simplified the inhibitor to identify relevant parts for target affinity. We then introduced new modifications to increase affinity, which was achieved by rational molecule design in combination with molecular modeling, chemical synthesis, isothermal titration calorimetry (ITC), and X-ray crystallography (XRD). With the newly developed inhibitors, we succeeded in generating cocrystal structures of *Pf*BD1 and *Pv*BD1, the BD of *Pv*BDP1 from *P. vivax*, showing detailed insights into the molecular interactions. Subsequently, the best inhibitors were tested for activity against various *P. falciparum* strains during their IDC. Conditional KD of *Pf*BDP1 was used for target validation, demonstrating differential sensitivity to one of the inhibitors when limiting *Pf*BDP1 expression. Additionally, in the presence of this inhibitor, *Pf*BDP1 was displaced from its target genes, as shown by Chromatin Immunoprecipitation (ChIP).

First, we determined the affinity of the pan-selective BDi MPM6 for *Pf*BD1 (5.5 µM, Table 1), which thus represented our first hit. Subsequently, we generated simplified variants of MPM6 to verify the contribution of the chloride and amino groups. Both resulting ligands showed slightly decreased affinity, with the contribution of the chloride (RMM1, 5.8 µM) found to be more important than that of the amino group (MPM2, 6.7 µM). Removal of both groups led to a more pronounced reduction in affinity (MPM3, 10 µM). The cocrystal structure of MPM2 with *Pf*BD1 (Fig. 1 A) is consistent with the apo structure (PDB ID 7M97), except for minor deviations of the side chains for Ile355 and an additional alternative conformation for Cys367 (Fig.1 A). The binding mode (BM) of the 4-acyl pyrrole corresponds to that known for human BDs (PDB ID 7R5B)^[14]^.The aniline moiety in the binding pocket (BP) of *Pf*BD1 adopts two 180° rotated conformations, but cannot displace the conserved water, as is the case for human BDs^[14]^ (SI. Fig. S1A). Initial models based on this structure and the affinity data suggested that small hydrophobic groups in ortho position should be beneficial. The resulting ligands (Table 1) confirmed this and showed that larger groups lead to weaker affinities. All molecules tested showed improved affinity over MPM3, but only methyl- and CF^3^ -groups (RMM2, 3.9 µM, and RMM4, 2.9 µM) led to better affinity than for the first hit (MPM6, 5.5 µM). The cocrystal structure of *Pf*BD1 in complex with RMM2 (Fig. 1B) in comparison with *Pf*BD1-MPM2 reveals a similar binding mode for both inhibitors (SI. Fig. S1B). Unlike MPM2, the phenyl group of RMM2 adopts only one conformation, with the methyl group forming hydrophobic interactions with the side chains of Ile355, the gatekeeper Val419, and the seven-membered ring. The phenyl ring is also tilted about 20° and is placed closer to the outermost conserved water.

**Table 1.**
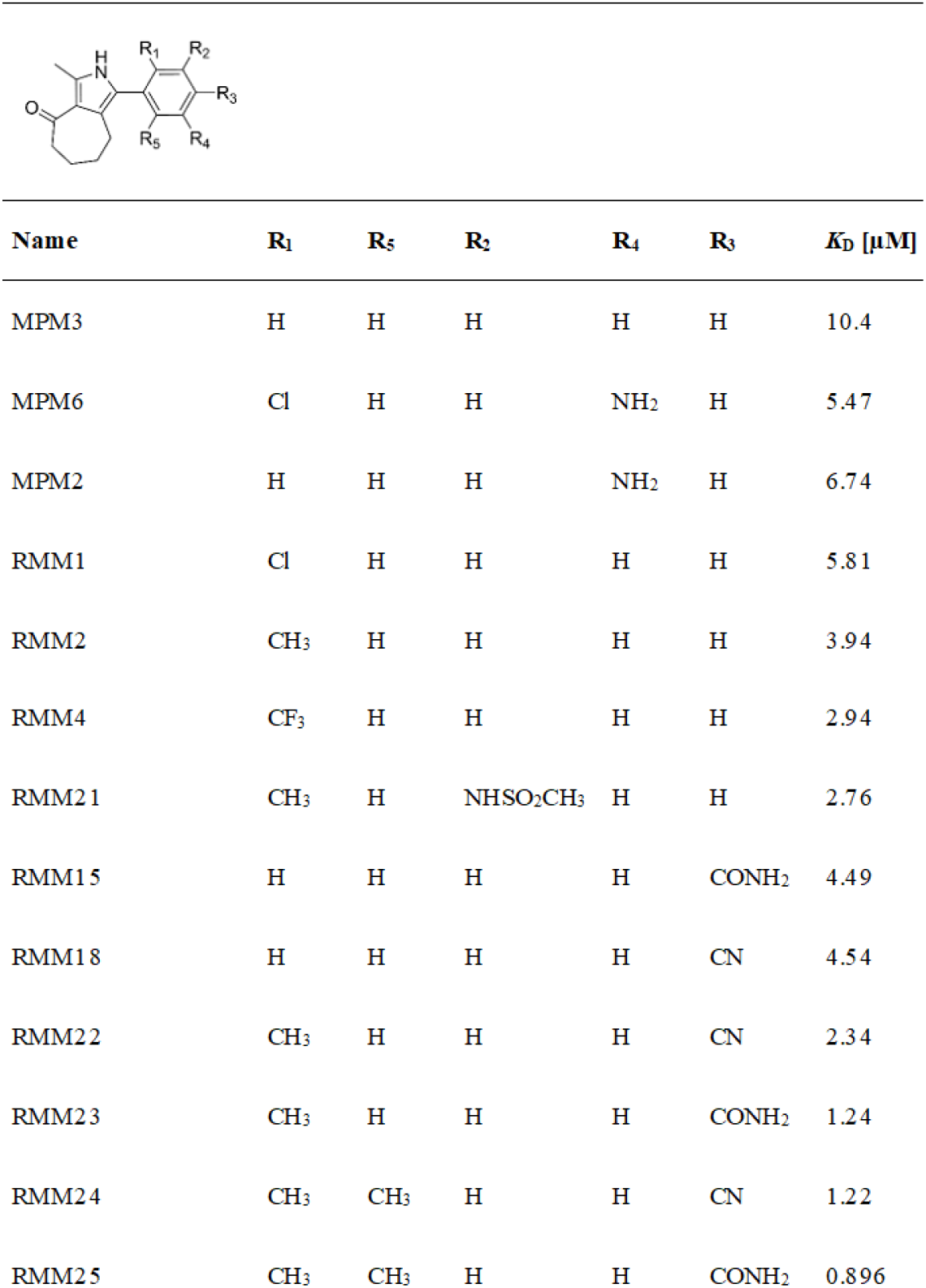
Series of synthesized compounds tested with ITC.

| Name | R<sub>1</sub> | R<sub>5</sub> | R<sub>2</sub> | R<sub>4</sub> | R<sub>3</sub> | K<sub>D</sub> [μM] |
| --- | --- | --- | --- | --- | --- | --- |
| MPM3 | H | H | H | H | H | 10.4 |
| MPM6 | Cl | H | H | NH<sub>2</sub> | H | 5.47 |
| MPM2 | H | H | H | NH<sub>2</sub> | H | 6.74 |
| RMM1 | Cl | H | H | H | H | 5.81 |
| RMM2 | CH<sub>3</sub> | H | H | H | H | 3.94 |
| RMM4 | CF<sub>3</sub> | H | H | H | H | 2.94 |
| RMM21 | CH<sub>3</sub> | H | NHSO<sub>2</sub>CH<sub>3</sub> | H | H | 2.76 |
| RMM15 | H | H | H | H | CONH<sub>2</sub> | 4.49 |
| RMM18 | H | H | H | H | CN | 4.54 |
| RMM22 | CH<sub>3</sub> | H | H | H | CN | 2.34 |
| RMM23 | CH<sub>3</sub> | H | H | H | CONH<sub>2</sub> | 1.24 |
| RMM24 | CH<sub>3</sub> | CH<sub>3</sub> | H | H | CN | 1.22 |
| RMM25 | CH<sub>3</sub> | CH<sub>3</sub> | H | H | CONH<sub>2</sub> | 0.896 |

**Figure 1:**
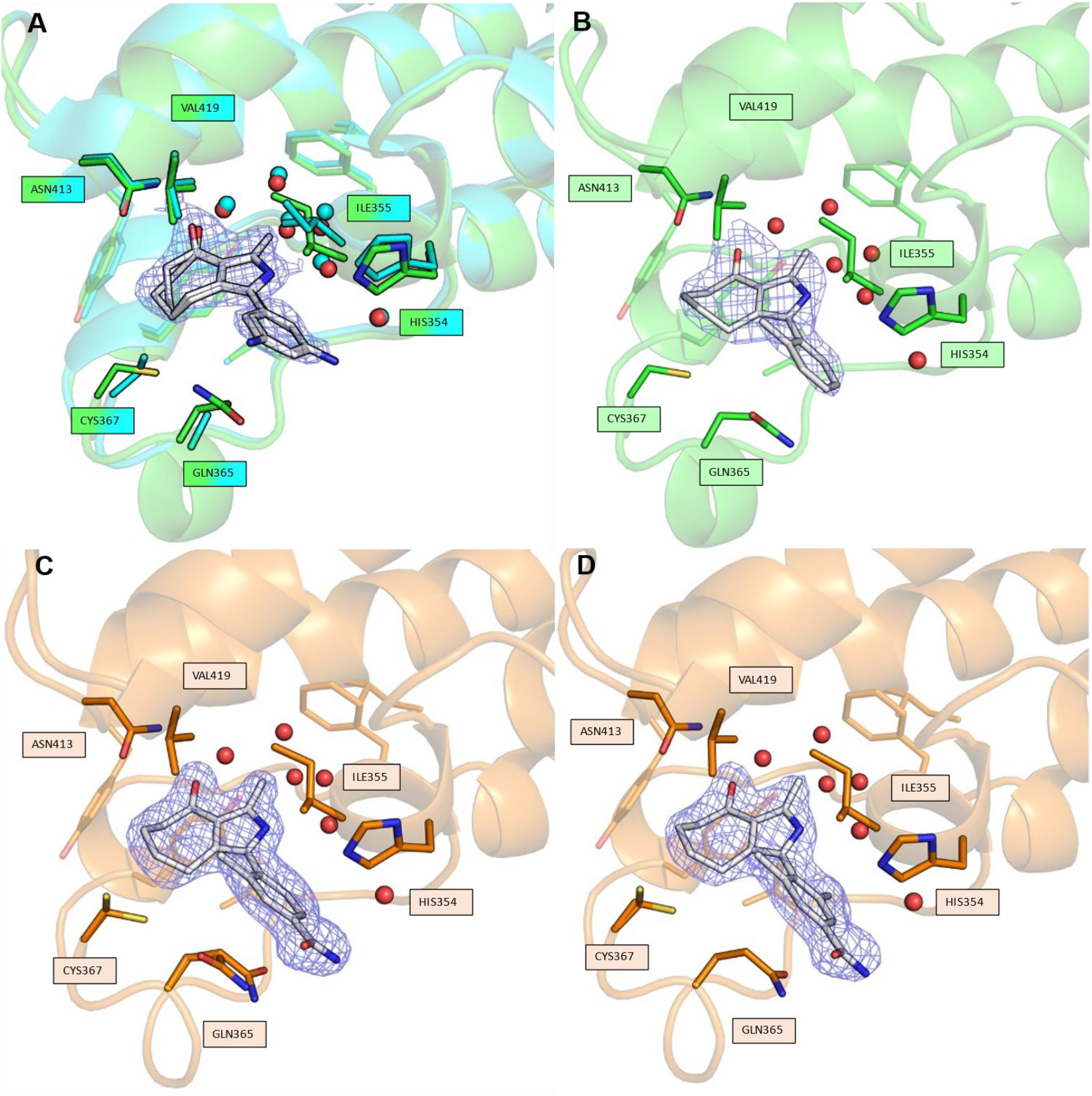
X-ray structures of PfBD1 and PvBD1 in complex with our designed ligands alongside their electron density maps contoured at 1.0 σ. **A:** *Pf*BD1 (green) complexed with MPM2 (white) overlayed with apo-*Pf*BD1 (teal; PDB ID:7M97). The conformation of the residues inside the BPs are almost identical except for showing an additional conformation for Cys367 and a side chain flip for Ile355. **B**: *Pf*BD1 (green) complexed with RMM2 (white). The *ortho*-methyl group forms hydrophobic interactions with the side chains of Ile355, the gatekeeper Val419, and the seven-membered ring. **C**: *Pv*BD1 (orange) complexed with RMM23 (white). Two conformations of GLN365 are visible. **D**: *Pv*BD1 (orange) complexed with RMM25 (white). The additional ortho-methyl group points towards the inner pocket.

In addition to the crystallization with *Pf*BD1, we successfully obtained cocrystal structures with the more stable homolog *Pv*BD1, which we first achieved with RMM21 (SI. Fig. S1C). The BP is indistinguishable from that of *Pf*BD1 and, like *Pf*BD1-MPM2, shows two conformations for Cys367 (Fig 1A). Apart from this, the BM of RMM4 is similar to *Pf*BD1-RMM2, although the alignment of a fluorine of the CF^3^ group between the side chains of Ile355 and the gatekeeper Val419 shifts the phenyl ring even further towards the outermost conserved water (SI. Fig. S1D).

Next, we investigated the influence of the *meta* position. Based on the affinity increase from MPM3 to MPM2, we assumed that hydrophilic groups improve affinity. Of the compounds tested (Table 1), the methyl variant (RMM6, 6.2 µM) showed a moderate affinity increase compared to MPM2 (6.7 µM), but only the sulfonamide-substituted forms (RMM11, 4.7 µM and RMM13, 4.9 µM) showed affinity improvement over the first hit (MPM6, 5.5 µM). Based on our models we assumed hydrogen bonds to the side chains of Gln365 and His354 for the sulfonamide variants. In the next step, we tried to enable hydrogen bonds via the para position, which should allow for similar hydrogen bonds, although unlike MPM2 not the whole phenyl ring would have to be rotated to align the functional group, and this should lead to better affinity. All hydrogen bond donors, as well as the cyanide group, resulted in better affinity than MPM2 and MPM3. This can be demonstrated particularly by the example of the differently methylated *p*-amides, in which the affinity collapses for the dimethylation, as hydrogen bonds can no longer be formed. For instance, both a single methyl group at the para position and a dimethylated para-amide result in a significant loss in affinity. (SI. Table S2). With the amide (RMM15, 4.49 µM) and the cyanide (RMM18, 4.54 µM) variants, we identified two molecules with slightly better affinity than the best meta-variant.

Next, we attempted to combine some of the best ortho modifications with some of the best meta or para substituents (Table 1). Since our models suggested that meta and para substituents form similar hydrogen bonds, we refrained from testing these combinations. Of the combinations tested, three molecules showed improvement over their predecessors. The complex structure of *Pv*BD1-RMM21 (SI Fig. S1C) largely resembles both the BP and the BM of *Pf*BD1-RMM2 (Fig. 1B), whereas the phenyl ring is oriented such that the meta substituent points in a single direction, unlike in *Pf*BD1-MPM2, where it alternates between two orientations. In addition to the interactions described for *Pf*BD1-RMM2, the structure of RMM21 confirmed the formation of hydrogen bonds to His354 or Gln365 with its sulfonamide, which is present in two conformations. The cocrystal structures of RMM23 with *Pf*BD1 (SI. Sup. Fig. S1E) and *Pv*BD1 (Fig. 1C) are largely identical, although the PvBD1 structure is much better defined and resolved, with the only difference being the visibility of two conformations of GLN365. BP and BM are very similar to *Pf*BD1-RMM2, with most of the methyl group orientated the same way, but also with a much smaller proportion rotated by 180°. The amide does not appear to interact directly with the BP in the crystal, but this does not have to be the case in solution. Notably, His354 forms a hydrogen bond to a symmetry equivalent and may be no longer available. In the next step, we tested the possibility of introducing a second *o*-methyl group. Based on the two conformations of RMM23 in the *Pf*BD1 cocrystal structure, this should be possible for RMM23 and RMM22, but not for RMM21 (compared to RMM20). The resulting ligands, RMM24 and RMM25 (Table 1), both show an improvement in affinity based on deteriorating enthalpy (ΔH), with simultaneous stronger improvement in entropy (ΔS) (SI. Sup. Fig. ITC). This indicates that no new preferential interactions are formed, but the rotation to align the methyl group, that is necessary for RMM23, is not required. This is further supported by the structure of *Pv*BD1-RMM25 (Fig. 1D), which shows an unchanged BM and BP compared to *Pv*BD1-RMM23 (Fig. 1C).

The activity of the compounds RMM23 and RMM25 was tested against *in vitro* blood stage *P. falciparum* cultures of the wild type strains 3D7 and NF54, as well as against the multidrug resistant K1 strain (chloroquine, pyrimethamine, and sulfadoxine resistant). The BDi compounds had modest activity against all three strains (Table 2). RMM25 showed the lowest half maximal effective concentration (EC50) against the K1 strain, whereas RMM23 was most active against NF54 (Table 2).

**Table 2.** Dose response of *P. falciparum* isolates to BD inhibitors.

|  | 3D7 | NF54 | K1 |
| --- | --- | --- | --- |
| Compound | EC50 (μM) | EC50 (μM) | EC50 (μM) |
| RMM23 | 18.33 (± 2.93) | 13.96 (± 3.25) | 20.9 (± 15.37) |
| RMM25 | 19.03 | 12.25 | 7.35 (± 6.51) |

Data presented as mean and SD (standard deviation) from 1-3 biological replicates

Treatment of synchronous ring stage 3D7 and NF54 parasite cultures with RMM23 resulted in significantly reduced growth after 48 h of treatment, indicating that the compound blocked parasite development within the same IDC when treatment was initiated (SI. Fig. S2A, B). Morphologically, RMM23 treated parasites developed normally into trophozoites by 24h, but then failed to undergo DNA replication and schizogony. Conversely, treatment commenced in the trophozoite stage resulted in normal schizogony, but a block in egress and reinvasion within 24 h, causing a greatly diminished number of ring stage infected erythrocytes relative to the DMSO control (SI. Fig. S1C, D). These results are consistent with those obtained by conditional KD of *Pf*BDP1 ^<u>11</u>^ and demonstrate that RMM23 can block parasite replication rapidly within the same IDC.

To validate whether the compounds indeed target the BD of *Pf*BDP1, we utilized a conditional KD system to modulate the expression levels of *Pf*BDP1 *in situ* ^[15]^. In the 3D7::*Pf*BDP1HADD line, *Pf*BDP1 is fused to a ligand-regulatable FKBP destabilization domain (DD) that targets the fusion protein for proteasomal degradation but can be stabilized by the Shield1 ligand, and 3xHA epitope tags for detection <u>^11^</u>. Titration of Shield1 from 500 to 10 nM resulted in diminished *Pf*BDP1 levels that still supported normal parasite growth (Fig. 2A). The conditional KD parasites (10 nM Shld1) exhibited increased sensitivity against RMM23, which was reflected by a leftward shift in the dose-response curve compared to standard conditions (500 nM Shld1) (Fig. 2B). In contrast, the EC50 of chloroquine was unchanged. This observation is consistent with RMM23 acting directly on *Pf*BDP1. In contrast, the parasites only showed moderately increased sensitivity to RMM25 in this assay (SI. Fig. S3), indicating that RMM25 may preferentially target the BD of another *Pf*BDP.

**Figure 2:**
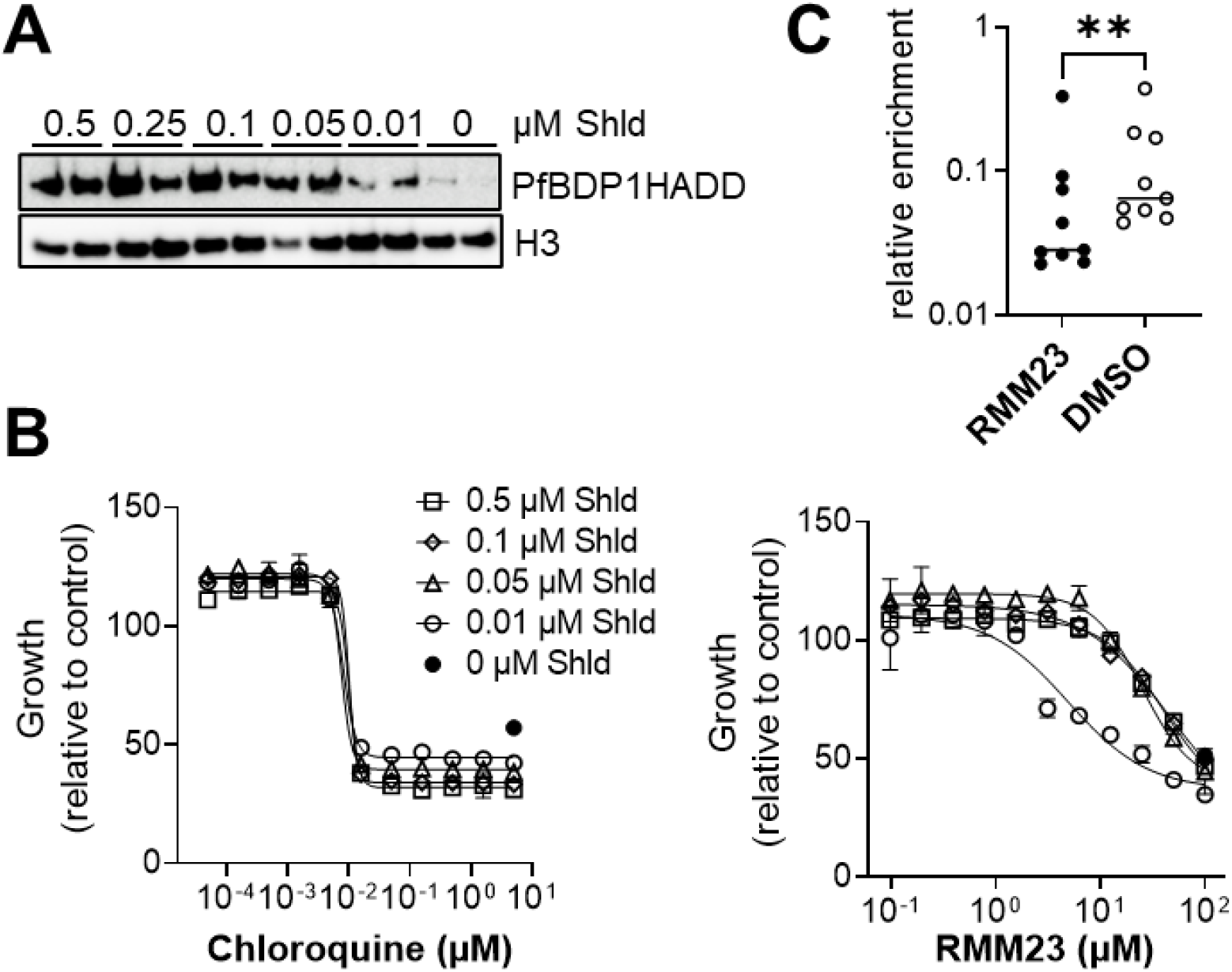
Target validation of RMM23. **A:** Western Blot analysis of 3D7::*Pf*BDP1HADD parasites cultivated in the presence of Shield1 (Shld) in concentrations ranging from 0 to 500 nM over 72 h in duplicates. *Pf*BDP1HADD was detected by anti-HA antibodies. Histone 3 (H3) was probed as a loading control. **B:** Dose-response assay of 3D7::*Pf*BDP1HADD parasites to RMM23 and chloroquine. Susceptibility to RMM23 increases under conditions of partial *Pf*BDP1 KD with 10 nM Shield1. Growth was determined relative to DMSO controls for each condition after 72 h of culture. N=4 replicates. **C**: ChIPqPCR assay of *Pf*BDP1 target regions (N=9) in 3D7::*Pf*BDP1HA parasites cultivated for 3 h in the presence of 50 μM RMM23 or DMSO as a control. Binding of *Pf*BDP1HA to target loci was calculated as enrichment relative to histone H3 ChIP performed in parallel. Shown are individual values and median of one representative experiment with technical duplicates. Paired t-test, p=0.0015.

*Pf*BDP1 is recruited to promoters of invasion-related genes by binding with its BD to acetylated histones <u>^11^</u>. To examine whether RMM23 inhibits chromatin binding of *Pf*BDP1, we performed ChIP of *Pf*BDP1HA in schizont stage parasites cultivated for 3 h with 50 μM RMM23 or DMSO control. Analysis of several known *Pf*BDP1 target loci by qPCR revealed significantly diminished enrichment of *Pf*BDP1 in the presence of RMM23 (Fig. 2C), further supporting the *Pf*BDP1 BD as the main target of RMM23 in *P. falciparum* parasites.

In summary, we developed inhibitors targeting *Pf*BDP1, an essential bromodomain protein in *P. falciparum*, and solved the first crystal structures of *Pf*BD1 and *Pv*BD1 with these inhibitors. Using *in vitro* assays combined with conditional knockdown of *Pf*BDP1, we identified RMM23 as a *Pf*BDP1-specific inhibitor that demonstrated effective binding and inhibition of parasite growth. These findings highlight the potential of therapeutically targeting bromodomain proteins in malaria parasites and suggest RMM23 as a promising candidate for further optimization towards clinical development.

## Supporting Information

The authors have cited additional references within the Supporting Information.

## Acknowledgements

M.A. was financially supported by the German Research Foundation (DFG; RTG2202). K.K.R and M.P. were supported by the German Research Foundation through the RTG2740 (Project number 447268119, Project B1).We thank the European Synchrotron Radiation Facility (ESRF) and the Swiss Light Source (Paul Scherrer Institute) for the beamtime and their technical support. We are thankful to Irene Wittmann for excellent technical support.

