## Supplementary Information for "A novel inhibitor against the bromodomain *Pf*BDP1 of the malaria pathogen *Plasmodium falciparum*"

#### Author Contributions

M.A. Conceptualization: Equal; Data curation: Lead; Formal analysis: Equal; Investigation: Lead; Methodology: Lead; Validation: Lead; Visualization: Lead; Writing—original draft: Lead; Writing—review & editing: Equal.

R.W. Conceptualization: Equal; Investigation: Lead; Methodology: Lead; Formal analysis: Equal; Writing—original draft: Equal; Writing—review & editing: Equal.

K.K.R. Investigation: Lead; Data curation: Equal; Visualization: Supporting; Writing—review & editing: Supporting.

P.H. Investigation: Supporting; Methodology: Supporting; Writing—review & editing: Supporting.

M.S. Methodology: Supporting; Writing—review & editing: Supporting.

S.C. Investigation: Supporting; Methodology: Supporting; Writing—review & editing: Supporting.

B.B. Resources: Equal; Supervision: Equal; Writing—review & editing: Supporting.

O.E. Resources: Equal; Supervision: Equal; Writing—review & editing: Supporting.

M.H. Conceptualization: Equal; Data curation: Supporting; Formal analysis: Equal; Investigation: Supporting; Methodology: Equal; Validation: Equal; Writing—original draft: Lead; Writing—review & editing: Equal.

M.P. Conceptualization: Equal; Investigation: Supporting; Methodology: Supporting; Supervision: Equal; Validation: Supporting; Writing—original draft: Equal; Writing—review & editing: Equal.

S.G. Conceptualization: Equal; Project design and administration: Lead; Supervision: Equal; Writing—original draft: Supporting; Writing—review & editing: Equal.

### Experimental procedures

##### Molecular modeling

Protein target preparation: Crystal structures were used as docking templates. Apo structure (PDB: 7M97) as well as co-crystallized structures (inhouse datasets) were prepared using the Protein Preparation Workflow (Release: 2021-4, Schrödinger, LLC, New York, NY, USA) with default parameters. During energy minimization only hydrogens were optimized using an OPLS4 force field. The conserved water molecules were kept inside the binding pocket for docking.

Docking experiment: Compounds were prepared using Ligprep (version 60137, Schrödinger, LLC) and docked using the XP (extra precision) algorithm in Glide (version 93137, Schrödinger, LLC). Top candidates were further evaluated using the MM-GBSA protocol (Schrödinger, LLC).

##### Plasmids and cloning

Genes of interest were synthesized (Azenta) and cloned into a pGEX-6P-1 vector using restriction enzyme digestion and ligated into the multiple cloning site of the vector. A TEV cleavage site between the fusion protein and the bromodomain was introduced for the PvBDP1 construct using FastCloning[ref]. All plasmids were transformed into chemical competent XL1-Blue cells, followed by plasmid preparation and sequencing to confirm the correctness of the plasmids. Supporting Table S1 shows the resulting protein sequences for all constructs.

##### Cell Growth and Purification

BL21* cells containing the construct were cultured in TB medium. A preculture was prepared in LB medium supplemented with Ampicillin (1:1000) and incubated overnight at 37 °C with shaking at 180 rpm. The next day, 1 L of TB medium (with buffer and 1 mL Ampicillin) was inoculated with 10 mL of the preculture and incubated at 37 °C and 180 rpm until an OD600 of ~2.5. The culture was cooled to 18 °C, induced with 0.2 mM IPTG, and incubated overnight. Cells were harvested by centrifugation (5000 x g, 15 min, 4 °C).

The cell pellet was resuspended in PBS (1:3 w/v). The suspension was lysed by sonication (15 min, 3 sec pulse, 10 sec pause, amplitude 70%) at 4 °C. Insoluble debris was removed by centrifugation at 80,000 x g for 45 min at 4 °C. The lysate was then sterile-filtered (0.45 µm) and purified by His affinity chromatography. The protein was cleaved using either PreScission Protease (PfBDP1-construct) or TEV Protease (PvBDP1-construct) separating the bromodomain from the fusion protein. Further separation of GST fusion protein and the bromodomain was achieved with another round of GST affinity chromatography followed by an additional size exclusion chromatography step using a HiLoad 26/600 Superdex 75 pg (GE Healthcare)

**Isothermal Titration Calorimetry (ITC)**

Affinity was measured using a MicroCal VP-ITC microcalorimeter (Malvern Instruments). Measurements were performed at 25 °C with the ligand (0.04 mM) in PBS buffer present in the cell and the protein (0.4 mM) injected via a stirred syringe in 5 min intervals (13.5 µL). Solutions were degassed prior to measurements. Automated baseline correction and peak integration were conducted using NITPIC[1,2]. Analysis, fitting, and thermodynamic calculations were performed using a binary interaction model in SEDPHAT[3], and plots were generated using GUSSI[4].

**Protein crystallization and structure determination**

Crystallization was set up using the sitting drop vapor diffusion method in Intelli-Plate 96-3 low-profile plates. The crystallization drops were dispensed using an OryxNano (Douglas Instruments) robot. For each condition, two drops were prepared: the first drop contained 33% protein solution and the second drop 50% protein solution, with a total drop size of 600 nL. Plates were stored at 277 K

X-ray diffraction data was collected at the Swiss Light Source (SLS) at the X06SA beamline as well as the European Synchrotron Radiation Facility (ESRF) at the beamlines MASSIF-3[5] and ID23-1[6]. The crystals were measured using a cryostream at 100 K.

Data were either processed using the ESRF autoprocessing pipeline with Autoproc[7]/STARANISO[8] or using the SLS processing pipeline[9,10]. Additional software used for data processing and structure determination included:

Aimless[11–13]: For scaling and merging of diffraction data.
Phaser[14]: For molecular replacement and phase determination (Initial model: apo structure of PfBD1 (PDB ID: 7m97) for PfBD1 and for PvBD1 Alphafold2 Predictions[15,16]).
PHENIX[17]: For automated refinement of the crystal structure.
Coot[18]: For manual model building and validation.
MolProbity[19]: Final structure validation

All structure illustrations were created with PyMOL (version 3.0.3)[20]

#### Cell assay

##### Parasite lines and culture

The following parasite lines were obtained from BEI resources NIAID, NIH: *Plasmodium falciparum* strain 3D7 (GL Clone), MRA-1001, and strain NF54 (Patient line E), MRA-1000, both contributed by Megan G. Dowler, as well as the multidrug-resistant strain K1, MRA-159, contributed by Dennis E. Kyle. The NF54::PfBDP1HA parasite line was generated by transfection of NF54 parasites with the plasmid pSLI_PfBDP1_3xHA and selection linked integration of the construct in the presence of G418[21]. The conditional *Pf*BDP1 knockdown parasite line 3D7::*Pf*BDP1HADD was generated as described previously by fusion of *Pf*BDP1 to a ligand-regulatable FKBP destabilization domain (DD), and 3xHA epitope tags for detection[22], and continuously cultivated in presence of 500 nM Shield1. Parasites were propagated in human red blood cells (blood group 0+) obtained from the Bavarian Red Cross service (BRK) and cultivated under standard conditions in RPMI 1640 (ThermoFisher) supplemented with 0.5% Albumax (ThermoFisher), 20 µg/ml Gentamycin, 25 mM Hepes (pH 7.3), and 0.02% hypoxanthine in an atmosphere of 1% $O_2$, 5% $CO_2$, 94% $N_2$.

##### Malaria SYBR green I based fluorescence assay

SYBR Green I based fluorescence assays to determine dose-response curves were performed essentially as described previously (Leidenberger et al., 2017). Briefly, parasite cultures were cultivated at 1% ring stage parasitemia and 2.5% haematocrit in 96 well plates in the presence of serial half-log or log2 dilutions of the compounds or solvent corresponding to the highest concentration (0.05%). Chloroquine (C6628, Sigma-Aldrich) was used as control. After 72 hours, cultures were lysed in lysis buffer (20 mM Tris-HCl, 5 mM EDTA, 0.008% saponin (Sigma-Aldrich), 0.08% Triton X (Sigma-Aldrich), 1:10,000 SYBR Green I dye (Invitrogen)) and fluorescence was measured in a Fluoroskan Ascent FL reader. The background fluorescence of wells containing uninfected red blood cells was subtracted and 50% effective concentrations (EC50s) were determined in GraphPad Prism 9.5.1 (GraphPad Software, Inc., San Diego, CA) by nonlinear regression (sigmoidal dose-response/variable slope equation).

#### Protein extraction and Western Blot

3D7::*Pf*BDP1HADD parasites were cultivated for 72 h with a starting parasitemia of 1% ring stage infection in the presence of varying concentrations of Shield1. Parasite protein extracts were prepared by first removing erythrocyte cytosolic components by saponin lysis, and after three washes with PBS at 4°C the resulting pellets were extracted with 2x Laemmli buffer at a concentration of $1x10^6$ IE/µl for 5 min at 95°C. The protein extracts were separated on 4-12% Bis-Tris bolt gels (Invitrogen) in 1x MES running buffer and transferred onto nitrocellulose membranes. Primary antibody incubation (rat anti-HA 3F10, Merck-Millipore; polyclonal rabbit anti-H3, Abcam) was performed over night at 4°C in TBS-T containing 5% non-fat milk powder, and horseradish peroxidase coupled secondary antibodies were incubated for 2 h at room temperature. Bands were visualized using Immobilon ECL Ultra substrate (Merck-Millipore).

#### Chromatin Immunoprecipitation and quantitative real time PCR (qPCR)

NF54::*Pf*BDP1HA cultures were tightly synchronized by treatment with 5% sorbitol at ring stage, and then allowed to progress to schizont stage. Around 48 hours post invasion (hpi), mature parasites were enriched by gelatine selection and allowed to reinvade for 4 hours before killing residual mature stages by sorbitol synchronization. At 40+/-2 hours post invasion, parasites were treated with 100 µM RW676 or DMSO for 3 hours before the cultures were fixed in 1% PFA for 10 min at 37°C. The reaction was quenched by addition of glycine to a final concentration of 125 mM and 5 min incubation on ice, followed by centrifugation at 2000 rpm for 5 min at 4°C. After lysis in 0.075% saponin in PBS, the released parasites were pelleted and washed three times in PBS. Nuclei were isolated by dounce homogenizing the pellets in lysis buffer (10 mM Hepes pH 7.9, 10 mM KCl, 0.1 mM EDTA pH 8.0, 0.1 mM EGTA pH 8.0). After centrifugation, the pellets were resuspended in SDS lysis buffer (1% SDS, 10 mM EDTA, 50 mM Tris pH 8.1) and sonicated with a Diagenode Bioruptor for 16 cycles, 30s on/off, set to High. Fragmented chromatin was diluted in chromatin dilution buffer (0.01% SDS, 1.1% Triton-X-100, 1.2 mM EDTA, 16.7 mM Tris-HCl pH 8.1, 150 mM NaCl) and after preclearing with BSA blocked protein G coupled sepharose 4 fast flow (GE Healthcare) for 1 hour, immunoprecipitation was conducted over night by adding rat anti-HA antibodies (clone 3F10, Roche) and protein G sepharose 4 fast flow. Washes, elution, decrosslinking, and DNA isolation using Qiagen MinElute columns were conducted as described previously (Quinn et al., 2022). The purified DNA was amplified with primers specific for *Pf*BDP1-associated genes in 10 µl reactions containing 2 µl of DNA (diluted 1:20) supplemented with 5 µl of SYBR green PCR Master Mix (Invitrogen) and 3 µl of each primer (3 µM). The qPCR was performed in duplicates in an Applied Biosystems 7900HT fast real-time PCR system. PfBDP1 enrichment was calculated as % of input DNA and normalized to H3.

#### Chemical compound synthesis

##### General Information

Air- or moisture-sensitive reaction were carried out in flame-dried glassware under argon atmosphere (Argon 5.0, *Sauerstoffwerke Friedrichshafen)* throughout the entire reaction. Therefor the glassware was flame-dried under an oil pump vacuum (0.1 mbar) then cooled back to rt and backflushed with Argon. Solvents and reagents were added under continuous argon flow. Syringes and cannulas were purged three times with argon prior to use.

##### Solvents and reagents

Reactions not sensitive to air and/or moisture were carried out using p.a. grade solvents without further purification.

Absolute solvents were dried using a M. BRAUN solvent purification system 800 before being filled into a flame-dried, argon flushed Straus flask or bought from Acros (AcroSeal) and degassed by purging with argon.

Reagents were obtained from commercial sources and used without purification unless otherwise stated.

##### Evaporation

Reaction mixtures were concentrated under reduced pressure at 40 °C or 55 °C for MeCN/H2O mixture after reversed phase chromatography on a rotation evaporator Laborota 4000 efficient from HEIDOLPH. Isolated compounds were further dried under high vacuum.

##### Chromatography

Thin layer chromatography was performed on aluminum plates coated with silica gel (MERCK, 60F254), which were visualized by UV fluorescence (λmax = 254 nm) and/or by staining with one of the following mixtures:

1% w/v Ninhydrine in EtOH + 1% AcOH

1% w/v KMnO4 in 0.5 M K2CO3

Flash Column Chromatography was performed using MACHERY-NAGEL silica gel 60® (230-400 mesh) and solvent mixtures as stated in the specific procedures.

Reversed phase column chromatography was performed on a puriFlash 430 from INTERCHIM using a PF-30C18HP-F0025 column with a flow rate of 15 mL/min. For reversed phase purifications following method was applied:

- column equilibration (90% H2O) for 5 min

- sample injection

- isocratic regime of 90% H2O for 5 min

- linear gradient from 90% H2O to pure MeCN over 45 min

- pure MeCN for 5 min

**Melting Points**

Melting points were measured on a STUART SMP10 melting point apparatus and are uncorrected.

**Nuclear Magnetic Resonance (NMR)**

NMR spectra were measured by the analytical department of the Institute of Organic Chemistry at the University of Freiburg on a BRUKER Avance 400 spectrometer (400.1 MHz and 100.6 MHz for 1H and 13C respectively) or a BRUKER Avance 500 (500.4 MHz and 125.8 MHz for 1H and 13C respectively) at a temperature of 300 or 303 K. Chemical shifts are reported in parts per million (ppm) relative to residual solvent signals: 7.26 ppm (CHCl3), 3.30 ppm (MeOD-d3), 2.04 ppm (Acetone-d5), 2.49 ppm (DMSO-d5) or 4.79 ppm (HDO). Data for 1H NMR are described as following: chemical shift (δ in ppm), multiplicity (s, singlet; d, doublet; t, triplet; q, quartet; m, multiplet; br, broad signal), coupling constant (Hz), integration. All 13C NMR are described in terms of chemical shift (δ in ppm). All 13C NMR spectra were reported in ppm relative to the solvent signal: 77.10 ppm (CDCl3), 49.00 ppm (MeOD-d4), 29.80 ppm (Acetone-d6) or 39.50 ppm (DMSO-d6) and were obtained with 1H decoupling. Multiplicities, coupling constants and integrals are reported as measured and might disagree with the expected values. All NMR data was analyzed and evaluated utilizing MestReNova by MESTRELAB RESEARCH.

**Mass Spectrometry**

Mass spectrometry was done by the analytical department of the Institute of Organic Chemistry at the University of Freiburg.

Electronspray ionization mass spectrometry (ESI) was performed on a LCQ Advantage or Exactive mass spectrometer from THERMO FISHER SCIENTIFIC. 2.5 µL/min of the sample solution were injected into a flow of 100–200 µL/min of MeOH or MeCN. The spray voltage was 4–5 kV and the ion transfer tube had a temperature of 250–300 °C.

Atmospheric pressure chemical ionization mass spectrometry (APCI) was performed on a LCQ Advantage instrument. 2.5 µL/min of the sample solution were injected into a flow of 200–400 µL/min of MeOH or MeCN. The spray current was 5 µA, the ion transfer tube had a temperature of 150–180 °C and the vaporizer had a temperature of 300–400 °C.

#### Nomenclature

Chemical structures and names were generated using ChemDraw Professional 19.1.

#### General Procedures

General procedure for Suzuki-Crosscoupling:

A mixture of 1-bromo-3-methyl-5,6,7,8-tetrahydrocyclohepta[c]pyrrol-4(2H)-one (1.0 equiv), Ar-B(OH)2 (1.5 equiv), K3PO4 (3.0 equiv), Pd2dba3 (5 mol%) and 1,3,5,7-tetramethyl-6-phenyl-2,4,8-trioxa-6-phosphaadamantane (20 mol%) in degassed dioxane/H2O (0.2 M, 1:1) was stirred at 70 °C for 3.5 h. The reaction mixture was filtered through a pad of celite, the solvents were removed under vacuum and the residue was purified by reversed phase column chromatography (MeCN/H2O). Reaction conditions based on literature.[23]

General procedure for Suzuki-Miyaura-Crosscoupling:

A mixture of Ar-Br (1.0 equiv), B2(pin)2 (1.2 equiv), KOAc (2.0 equiv) and Pd(dppf)Cl2 (6 mol%) in degassed dioxane (0.3 M with 2% DMSO) was stirred at 90 °C overnight, filtered through a pad of celite and after removal of the solvents used without further purification.

A mixture of 1-bromo-3-methyl-5,6,7,8-tetrahydrocyclohepta[c]pyrrol-4(2H)-one (1.0 equiv), crude Ar-B(pin) (1.5 equiv), K3PO4 (3.0 equiv), Pd2dba3 (5 mol%) and 1,3,5,7-tetramethyl-6-phenyl-2,4,8-trioxa-6-phosphaadamantane (20 mol%) in degassed dioxane/H2O (0.2 M, 1:1) was stirred at 70 °C for 3.5 h. The reaction mixture was filtered through a pad of celite, the solvents were removed under vacuum and the residue was purified by reversed phase column chromatography (MeCN/H2O).

The synthesis for MPM2 and MPM6 were according to literature[24].

**3-Methyl-1-phenyl-5,6,7,8-tetrahydrocyclohepta[*c*]pyrrol-4(2H)-one (208; MPM3)**

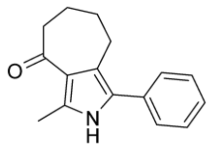

Chemical Formula: $C_{16}H_{17}NO$
Molecular Weight: 239,32

The title compound was obtained as a colourless solid (16 mg, 66 µmol, **79%**) according to the general procedure for Suzuki-Crosscoupling using 1-bromo-3-methyl-5,6,7,8-tetrahydrocyclohepta[*c*]pyrrol-4(2*H*)-one (20 mg, 83 µmol, 1.0 equiv).

**Mp:** 178-180 °C.

**$^{1}$H NMR (500 MHz, $CDCl_3$):** δ = 1.83 ($m_c$, 2H), 1.86 – 1.92 (m, 2H), 2.55 (s, 3H), 2.67 – 2.71 (m, 2H), 2.85 – 2.89 (m, 2H), 7.27 – 7.31 (m, 1H), 7.34 – 7.38 (m, 2H), 7.39 – 7.43 (m, 2H), 8.38 (s, 1H).

**$^{13}$C NMR (126 MHz, $CDCl_3$):** δ = 13.7, 22.3, 23.5, 26.0, 41.9, 121.9, 122.5, 126.5, 126.9, 127.5, 128.9, 132.6, 135.0, 200.5.

**HRMS (pos. APCI): $[M + H]^+$** calcd. for $C_{16}H_{18}NO$: 240.1383; found: 240.1383.

**3-Methyl-1-(*m*-tolyl)-5,6,7,8-tetrahydrocyclohepta[*c*]pyrrol-4(2*H*)-one (292; RMM6)**

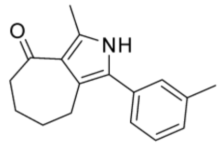

Chemical Formula: $C_{17}H_{19}NO$
Molecular Weight: 253,35

3-Methyl-1-(*m*-tolyl)-5,6,7,8-tetrahydrocyclohepta[*c*]pyrrol-4(2*H*)-one was obtained (9.7 mg, 38 µmol, **64%**) as a brown oil according to the general procedure for Suzuki-Miyaura-Crosscoupling using 1-bromo-3-methyl-5,6,7,8-tetrahydrocyclohepta[*c*]pyrrol-4(2*H*)-one (15 mg, 60 µmol, 1.0 equiv).

**$^{1}$H NMR (400 MHz, $CDCl_3$):** δ = 1.78 – 1.95 (m, 4H), 2.39 (d, *J* = 0.7 Hz, 3H), 2.55 (d, *J* = 0.4 Hz, 3H), 2.66 – 2.73 (m, 2H), 2.83 – 2.90 (m, 2H), 7.09 – 7.19 (m, 3H), 7.27 – 7.33 (m, 1H), 8.20 (s, 1H).

**$^{13}$C NMR (101 MHz, $CDCl_3$):** δ = 13.8, 21.7, 22.4, 23.7, 26.1, 42.0, 121.9, 122.6, 124.7, 126.7, 127.8, 128.2, 128.8, 132.7, 134.8, 138.6, 200.4.

**HRMS (pos. ESI): $[M + H]^+$** calcd. for $C_{17}H_{20}ON$: 254.1539; found: 254.1537.

**3-Methyl-1-(3-nitrophenyl)-5,6,7,8-tetrahydrocyclohepta[*c*]pyrrol-4(2*H*)-one (293; RMM7)**

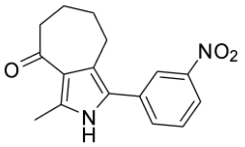

Chemical Formula: $C_{16}H_{16}N_2O_3$
Molecular Weight: 284,32

The title compound was obtained as a yellow solid (17 mg, 61 µmol, **74%**) according to the general procedure for Suzuki-Crosscoupling using 1-bromo-3-methyl-5,6,7,8-tetrahydrocyclohepta[*c*]pyrrol-4(2*H*)-one (20 mg, 83 µmol, 1.0 equiv).

**Mp:** 182-183 °C.

**$^1$H NMR (500 MHz, $CDCl_3$):** δ = 1.82 – 1.93 (m, 4H), 2.57 (s, 3H), 2.68 – 2.72 (m, 2H), 2.85 – 2.89 (m, 2H), 7.58 (t, J = 8.0 Hz, 1H), 7.69 (dt, J = 7.7, 1.4 Hz, 1H), 8.11 (ddd, J = 8.2, 2.3, 1.1 Hz, 1H), 8.21 (t, J = 2.0 Hz, 1H), 8.69 (s, 1H).

**$^{13}$C NMR (126 MHz, $CDCl_3$):** δ = 13.7, 22.2, 23.6, 25.9, 41.8, 121.3, 121.8, 122.9, 123.9, 124.2, 129.8, 133.1, 134.3, 136.3, 148.7, 200.3.

**HRMS (neg. ESI): $[M - H]^-$** calcd. for $C_{16}H_{15}N_2O_3$: 283.1088; found: 283.1086.

**3-Methyl-1-(3-(methylamino)phenyl)-5,6,7,8-tetrahydrocyclohepta[*c*]pyrrol-4(2H)-one (294) and 1-(3-(dimethylamino)phenyl)-3-methyl-5,6,7,8-tetrahydrocyclohepta[*c*]pyrrol-4(2*H*)-one (295;RMM8, RMM9)**

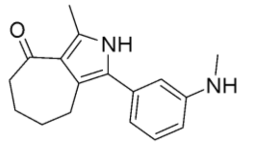

Chemical Formula: $C_{17}H_{20}N_2O$
Molecular Weight: 268,36

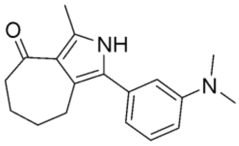

Chemical Formula: $C_{18}H_{22}N_2O$
Molecular Weight: 282,39

A mixture of 1-(3-aminophenyl)-3-methyl-5,6,7,8-tetrahydrocyclohepta[*c*]pyrrol-4(2*H*)-one **207** (21 mg, 83 µmol, 1.0 equiv), MeI (6.5 µL, 14 mg, 0.10 mmol, 1.25 equiv) and $KHCO_3$ (12 mg, 0.12 mmol, 1.5 equiv) in DMF (0.13 mL) was stirred at rt for 7.5 h. $H_2O$ and EtOAc were added and the aqueous phase was extracted three times with EtOAc, the combined organic layers were dried over $Na_2SO_4$ and evaporated. The residue was purified by reversed phase column chromatography (MeCN/$H_2O$) to obtain the desired products as brown oils 3-methyl-1-(3-(methylamino)phenyl)-5,6,7,8-tetrahydrocyclohepta[*c*]pyrrol-4(2H)-one (3.9 mg, 15 µmol, **17%**) and 1-(3-(dimethylamino)phenyl)-3-methyl-5,6,7,8-tetrahydrocyclohepta[*c*]pyrrol-4(2*H*)-one (6.3 mg, 22 µmol, **27%**).

**3-Methyl-1-(3-(methylamino)phenyl)-5,6,7,8-tetrahydrocyclohepta[*c*]pyrrol-4(2*H*)-one (RMM8):**

**$^1$H NMR (500 MHz, $CDCl_3$):** δ = 1.77 – 1.92 (m, 4H), 2.55 (s, 3H), 2.66 – 2.72 (m, 2H), 2.85 – 2.91 (m, 5H), 6.59 – 6.65 (m, 2H), 6.73 (dt, *J* = 7.6, 1.2 Hz, 1H), 7.24 (t, *J* = 7.8 Hz, 1H), 8.20 (s, 1H).

**$^{13}$C NMR (126 MHz, $CDCl_3$):** δ = 13.8, 22.4, 23.6, 26.1, 31.1, 41.9, 111.6, 111.7, 117.1, 121.8, 122.5, 127.0, 129.8, 133.7, 134.6, 149.3, 200.5.

**HRMS (neg. ESI): $[M - H]^-$** calcd. for $C_{17}H_{19}N_2O$: 267.1503; found: 267.1502.

**1-(3-(Dimethylamino)phenyl)-3-methyl-5,6,7,8-tetrahydrocyclohepta[*c*]pyrrol-4(2*H*)-one (RMM9):**

**$^{1}$H NMR (500 MHz, $CDCl_3$):** δ = 1.79 – 1.93 (m, 4H), 2.55 (s, 3H), 2.67 – 2.72 (m, 2H), 2.87 – 2.92 (m, 2H), 3.00 (s, 6H), 6.69 – 6.75 (m, 3H), 7.29 (t, *J* = 8.1 Hz, 1H), 8.15 – 8.24 (m, 1H).

**$^{13}$C NMR (126 MHz, $CDCl_3$):** δ = 13.8, 22.4, 23.6, 26.1, 40.9, 41.9, 111.6, 112.0, 116.4, 121.8, 122.5, 127.3, 129.7, 133.5, 134.6, 150.7, 200.5.

**HRMS (pos. ESI): $[M + H]^+$** calcd. for $C_{18}H_{23}N_2O$: 283.1805; found: 283.1802.

***N*-(3-(3-Methyl-4-oxo-2,4,5,6,7,8-hexahydrocyclohepta[*c*]pyrrol-1-yl)phenyl)acetamide (296; RMM10)**

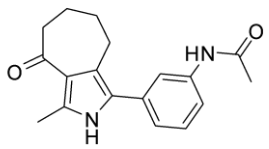

Chemical Formula: $C_{18}H_{20}N_2O_2$
Molecular Weight: 296,37

Acetyl chloride (4.5 µL, 5.0 mg, 63 µmol, 1.5 equiv) was added dropwise to a solution of **207** (11 mg, 42 µmol, 1.0 equiv) and $NEt_3$ (12 µL, 8.6 mg, 85 µmol, 2.0 equiv) in dry DCM (0.42 mL). The reaction mixture was stirred at rt overnight. Aqueous HCl (1 M) was added and the aqueous phase was extracted three times with DCM, the combined organic layers were washed with $H_2O$, dried over $Na_2SO_4$ and concentrated in vacuo. The residue was purified by reversed phase column chromatography (MeCN/$H_2O$) to obtain the title compound as a colourless solid (6.2 mg, 21 µmol, **49%**).

**Mp:** < 200 °C.

**$^{1}$H NMR (500 MHz, $MeOD_4/CDCl_3$, 1:1):** δ = 1.77 – 1.89 (m, 4H), 2.13 (s, 3H), 2.47 (s, 3H), 2.62 – 2.66 (m, 2H), 2.84 – 2.88 (m, 2H), 7.11 (dt, J = 7.3, 1.6 Hz, 1H), 7.27 (dt, J = 8.1, 1.6 Hz, 1H), 7.29 – 7.33 (m, 1H), 7.63 (t, J = 1.7 Hz, 1H).

**$^{13}$C NMR (126 MHz, $MeOD_4/CDCl_3$, 1:1):** δ = 13.5, 22.9, 23.9, 23.9, 26.4, 41.9, 119.0, 120.1, 122.4, 122.8, 124.0, 127.6, 129.6, 134.0, 136.9, 139.2, 171.2, 202.6.

**HRMS (pos. APCI): $[M + H]^+$** calcd. for $C_{18}H_{21}N_2O_2$: 297.1598; found: 297.1597.

**N-(3-(3-Methyl-4-oxo-2,4,5,6,7,8-hexahydrocyclohepta[c]pyrrol-1-yl)phenyl)methanesulfonamide (297; RMM1)**

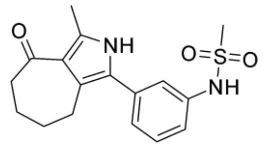

Chemical Formula: $C_{17}H_{20}N_2O_3S$
Molecular Weight: 332,42

Methanesulfonyl chloride (3.9 µL, 5.7 mg, 50 µmol, 1.0 equiv) was added to a solution of 1-(3-aminophenyl)-3-methyl-5,6,7,8-tetrahydrocyclohepta[*c*]pyrrol-4(2*H*)-one **207** (13 mg, 50 µmol, 1.0 equiv) and pyridine (6.0 µL, 5.9 mg, 75 µmol, 1.5 equiv) in DCM (0.1 mL) at 0 °C. The mixture was allowed to return to rt overnight. Aq. HCl (0.5 M) was added and the aqueous phase was extracted three times with DCM. The combined organic layers were dried over $Na_2SO_4$, evaporated and the residue was purified by reversed phase column chromatography (MeCN/$H_2O$) to obtain the desired product as a beige solid (10 mg, 32 µmol, **63%**).

**Mp:** >200 °C.

**$^1$H NMR (500 MHz, Acetone-d$_6$):** δ = 1.79 – 1.86 (m, 4H), 2.47 (d, *J* = 0.5 Hz, 3H), 2.56 – 2.62 (m, 2H), 2.86 – 2.91 (m, 2H), 3.01 (s, 3H), 7.20 – 7.24 (m, 2H), 7.39 (t, *J* = 7.9 Hz, 1H), 7.43 (t, *J* = 1.9 Hz, 1H), 8.63 (s, 1H), 10.49 (s, 1H).

**$^{13}$C NMR (126 MHz, Acetone-d$_6$):** δ = 13.2, 22.8, 24.0, 26.7, 39.5, 42.1, 118.8, 119.8, 122.5, 123.1, 123.9, 126.8, 130.4, 135.0, 135.6, 139.7, 199.0.

**HRMS (pos. ESI): $[M + H]^+$** calcd. for $C_{17}H_{21}N_2O_3S$: 333.1267; found: 333.1265.

**3-(3-Methyl-4-oxo-2,4,5,6,7,8-hexahydrocyclohepta[*c*]pyrrol-1-yl)benzamide (298; MPM8)**

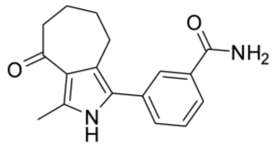

Chemical Formula: $C_{17}H_{18}N_2O_2$
Molecular Weight: 282,34

The title compound was obtained as a colourless gum (6.9 mg, 25 µmol, **30%**) according to the general procedure for Suzuki-Crosscoupling using 1-bromo-3-methyl-5,6,7,8-tetrahydrocyclohepta[*c*]pyrrol-4(2*H*)-one (20 mg, 83 µmol, 1.0 equiv).

**$^1$H NMR (500 MHz, MeOD$_4$/CDCl$_3$, 2:1):** δ = 1.78 – 1.83 (m, 2H), 1.87 (tdd, J = 9.8, 7.9, 3.0 Hz, 2H), 2.48 (s, 3H), 2.64 – 2.68 (m, 2H), 2.84 – 2.88 (m, 2H), 7.47 (td, J = 7.7, 0.4 Hz, 1H), 7.54 (ddd, J = 7.7, 1.8, 1.2 Hz, 1H), 7.72 (ddd, J = 7.7, 1.8, 1.2 Hz, 1H), 7.87 – 7.89 (m, 1H).

**$^{13}$C NMR (126 MHz, MeOD$_4$/CDCl$_3$, 2:1):** δ = 13.5, 23.1, 24.1, 26.6, 42.1, 48.5, 122.7, 123.4, 126.1, 127.2, 127.5, 129.5, 131.7, 134.1, 134.8, 137.4, 172.0, 202.7.

**HRMS (neg. ESI): [M - H]$^-$** calcd. for $C_{17}H_{17}N_2O_2$: 281.1296; found: 281.1295.

**3-(3-Methyl-4-oxo-2,4,5,6,7,8-hexahydrocyclohepta[*c*]pyrrol-1-yl)benzenesulfonamide (299; RMM13)**

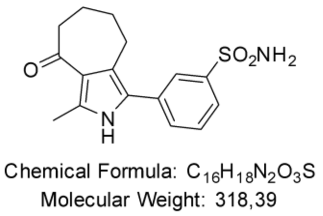

3-(3-Methyl-4-oxo-2,4,5,6,7,8-hexahydrocyclohepta[*c*]pyrrol-1-yl)benzenesulfonamide was obtained as a colorless wax (20 mg, 63 µmol, **76%**) according to the general procedure for Suzuki-Crosscoupling using 1-bromo-3-methyl-5,6,7,8-tetrahydrocyclohepta[*c*]pyrrol-4(2*H*)-one (20 mg, 83 µmol, 1.0 equiv).

**$^1$H NMR (500 MHz, MeOD-*d*$_4$):** δ = 1.86 ($m_c$, 2H), 2.48 (s, 3H), 2.67 ($m_c$, 2H), 2.88 ($m_c$, 2H), 7.56 – 7.60 (m, 2H), 7.79 (dt, *J* = 6.9, 1.9Hz, 1H), 7.92 – 7.93 (m, 1H).

**$^{13}$C NMR (126 MHz, MeOD-*d*$_4$):** δ = 13.4, 23.4, 24.3, 26.9, 42.3, 123.1, 124.3, 124.9, 125.9, 127.0, 130.4, 131.9, 135.0, 138.0, 145.5, 202.9.

**HRMS (neg. ESI) (*m/z*): [M−H]$^-$** calcd. for $C_{16}H_{17}N_2O_3S$: 317.0965; found: 317.0970.

**3-Methyl-1-(*o*-tolyl)-5,6,7,8-tetrahydrocyclohepta[*c*]pyrrol-4(2*H*)-one (305; RMM2)**

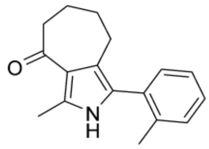

Chemical Formula: $C_{17}H_{19}NO$
Molecular Weight: 253,35

The title compound was obtained (18 mg, 71 µmol, **86%**) as a yellow oil according to the general procedure for Suzuki-Crosscoupling using 1-bromo-3-methyl-5,6,7,8-tetrahydrocyclohepta[*c*]pyrrol-4(2*H*)-one (20 mg, 83 µmol, 1.0 equiv).

**$^{1}$H NMR (400 MHz, $CDCl_3$):** δ = 1.76 ($m_c$, 2H), 1.83 – 1.90 (m, 2H), 2.25 (s, 3H), 2.53 – 2.54 (m, 3H), 2.54 – 2.58 (m, 2H), 2.63 – 2.69 (m, 2H), 7.20 – 7.26 (m, 2H), 7.26 – 7.28 (m, 1H), 8.18 (s, 1H).

**$^{13}$C NMR (101 MHz, $CDCl_3$):** δ = 13.8, 20.2, 22.3, 23.6, 25.7, 42.0, 121.3, 122.4, 125.6, 125.7, 128.1, 130.4, 131.1, 132.1, 134.3, 137.7, 200.5.

**HRMS (neg. ESI): $[M – H]^-$** calcd. for $C_{17}H_{18}NO$: 252.1394; found: 252.1395.

**1-(2-Ethylphenyl)-3-methyl-5,6,7,8-tetrahydrocyclohepta[*c*]pyrrol-4(2*H*)-one (306; RMM3)**

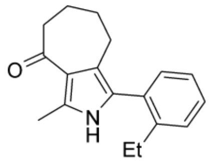

Chemical Formula: $C_{18}H_{21}NO$
Molecular Weight: 267,37

The title compound was obtained as a brown solid (22 mg, 83 µmol, **quant.**) according to the general procedure for Suzuki-Crosscoupling using 1-bromo-3-methyl-5,6,7,8-tetrahydrocyclohepta[*c*]pyrrol-4(2*H*)-one (20 mg, 83 µmol, 1.0 equiv).

**Mp:** 126-128 °C.

**$^{1}$H NMR (500 MHz, $CDCl_3$):** δ = 1.09 (t, J = 7.6 Hz, 3H), 1.72 – 1.78 (m, 2H), 1.83 – 1.89 (m, 2H), 2.53 – 2.59 (m, 7H), 2.64 – 2.67 (m, 2H), 7.18 – 7.24 (m, 2H), 7.29 – 7.35 (m, 2H), 8.13 (s, 1H).

**$^{13}$C NMR (126 MHz, $CDCl_3$):** δ = 13.8, 15.5, 22.3, 23.5, 25.7, 26.3, 42.0, 121.2, 122.3, 125.5, 125.7, 128.6, 128.6, 131.4, 131.5, 134.2, 144.2, 200.6.

**HRMS (pos. APCI): $[M + H]^+$** calcd. for $C_{16}H_{22}NO$: 268.1696; found: 268.1699.

**1-(2-Chlorophenyl)-3-methyl-5,6,7,8-tetrahydrocyclohepta[c]pyrrol-4(2H)-one (307; RMM1)**

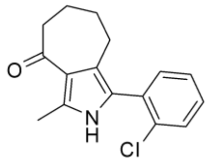

Chemical Formula: $C_{16}H_{16}ClNO$
Molecular Weight: 273,76

The title compound was obtained as a beige solid (18 mg, 66 µmol, **80%**) according to the general procedure for Suzuki-Crosscoupling using 1-bromo-3-methyl-5,6,7,8-tetrahydrocyclohepta[*c*]pyrrol-4(2*H*)-one (20 mg, 83 µmol, 1.0 equiv).

**Mp:** 136-138 °C.

**$^{1}$H NMR (500 MHz, $CDCl_3$):** δ = 1.78 – 1.84 (m, 2H), 1.84 – 1.91 (m, 2H), 2.55 (s, 3H), 2.65 – 2.69 (m, 4H), 7.25 – 7.33 (m, 3H), 7.43 – 7.49 (m, 1H), 8.44 (s, 1H).

**$^{13}$C NMR (126 MHz, $CDCl_3$):** δ = 13.9, 22.2, 23.7, 25.6, 42.0, 121.5, 123.2, 123.9, 126.9, 129.0, 130.2, 131.1, 132.4, 133.6, 135.0, 200.3.

**HRMS (pos. APCI): [M + H]$^+$** calcd. for $C_{16}H_{17}NOCl$: 274.0993; found: 274.0995.

**3-Methyl-1-(2-(trifluoromethyl)phenyl)-5,6,7,8-tetrahydrocyclohepta[*c*]pyrrol-4(2*H*)-one (308; RMM4)**

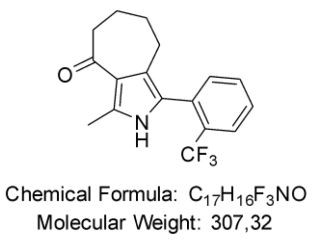

The title compound was obtained as a colourless solid (18 mg, 58 µmol, **70%**) according to the general procedure for Suzuki-Crosscoupling using 1-bromo-3-methyl-5,6,7,8-tetrahydrocyclohepta[*c*]pyrrol-4(2*H*)-one (20 mg, 83 µmol, 1.0 equiv).

**Mp:** 148-150 °C.

**$^{1}$H NMR (500 MHz, $CDCl_3$):** δ = 1.72 – 1.78 (m, 2H), 1.82 – 1.89 (m, 2H), 2.53 (s, 3H), 2.55 – 2.58 (m, 2H), 2.64 – 2.68 (m, 2H), 7.36 – 7.40 (m, 1H), 7.46 – 7.51 (m, 1H), 7.55 – 7.60 (m, 1H), 7.74 – 7.78 (m, 1H), 8.20 (s, 1H).

**$^{13}$C NMR (126 MHz, $CDCl_3$):** δ = 13.7, 22.1, 23.3, 25.7, 42.0, 121.2, 122.7, 123.8, 125.2 (q, *J* = 273.5 Hz), 126.5 (q, *J* = 5.3 Hz), 128.3, 129.9 (q, *J* = 29.7 Hz), 131.0, 131.7, 133.9, 134.8, 200.4.

**$^{19}$F-NMR (471 MHz, $CDCl_3$):** δ = -59.2.

**HRMS (neg. ESI): $[M - H]^-$** calcd. for $C_{17}H_{15}NOF_3$: 306.1111; found: 306.1116.

**1-(2-Methoxyphenyl)-3-methyl-5,6,7,8-tetrahydrocyclohepta[*c*]pyrrol-4(2*H*)-one (309; RMM5)**

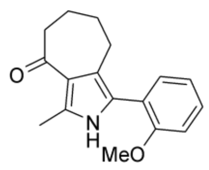

Chemical Formula: $C_{17}H_{19}NO_2$
Molecular Weight: 269,34

The title compound was obtained as a yellow-brown solid (21 mg, 78 µmol, **95%**) according to the general procedure for Suzuki-Crosscoupling using 1-bromo-3-methyl-5,6,7,8-tetrahydrocyclohepta[*c*]pyrrol-4(2*H*)-one (20 mg, 83 µmol, 1.0 equiv).

**Mp:** 147-149 °C.

**$^{1}$H NMR (500 MHz, $CDCl_3$):** δ = 1.80 – 1.86 (m, 2H), 1.86 – 1.93 (m, 2H), 2.56 (s, 3H), 2.67 – 2.71 (m, 2H), 2.80 – 2.85 (m, 2H), 3.87 (s, 3H), 6.98 (dd, J = 8.2, 1.0 Hz, 1H), 7.02 (td, J = 7.5, 1.1 Hz, 1H), 7.26 – 7.32 (m, 2H), 8.69 (s, 1H).

**$^{13}$C NMR (126 MHz, $CDCl_3$):** δ = 14.0, 22.3, 24.0, 26.0, 41.9, 55.7, 111.3, 120.9, 121.0, 121.7, 122.9, 123.2, 128.3, 130.6, 134.3, 156.4, 200.2.

**HRMS (pos. APCI): [M + H]$^+$** calcd. for $C_{17}H_{20}NO_2$: 270.1489; found: 270.1491.

**4-(3-Methyl-4-oxo-2,4,5,6,7,8-hexahydrocyclohepta[*c*]pyrrol-1-yl)benzamide (310; RMM15)**

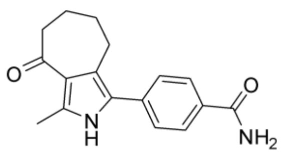

Chemical Formula: $C_{17}H_{18}N_2O_2$
Molecular Weight: 282,34

4-(3-Methyl-4-oxo-2,4,5,6,7,8-hexahydrocyclohepta[*c*]pyrrol-1-yl)benzamide was obtained (14 mg, 50 µmol, **62%**) as a colourless solid according to the general procedure for Suzuki-Crosscoupling using 1-bromo-3-methyl-5,6,7,8-tetrahydrocyclohepta[*c*]pyrrol-4(2*H*)-one (19 mg, 80 µmol, 1.0 equiv).

**Mp:** >200 °C.

**$^1$H NMR (400 MHz, MeOD-*d*$_4$):** δ = 1.77 – 1.92 (m, 3H), 2.48 (s, 4H), 2.65 – 2.69 (m, 2H), 2.87 – 2.93 (m, 2H), 7.46 – 7.50 (m, 2H), 7.89 – 7.93 (m, 2H).

**$^{13}$C NMR (101 MHz, MeOD-*d*$_4$):** δ = 13.4, 23.4, 24.5, 26.9, 42.3, 123.3, 124.5, 128.2, 129.1, 129.9, 130.0, 132.5, 133.0, 133.1, 137.6, 172.0, 202.9.

**HRMS (neg. ESI): [M – H]$^+$** calcd. for $C_{17}H_{17}N_2O_2$: 281.1296; found: 281.1294.

**N-Methyl-4-(3-methyl-4-oxo-2,4,5,6,7,8-hexahydrocyclohepta[c]pyrrol-1-yl)benzamide (311; RMM16)**

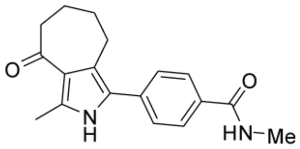

Chemical Formula: $C_{18}H_{20}N_2O_2$
Molecular Weight: 296,37

The title compound was obtained as a colourless solid (4.8 mg, 16 µmol, **20%**) according to the general procedure for Suzuki-Crosscoupling using 1-bromo-3-methyl-5,6,7,8-tetrahydrocyclohepta[*c*]pyrrol-4(2*H*)-one (20 mg, 83 µmol, 1.0 equiv) and 1.0 equiv of the corresponding arylboronic acid.

**Mp:** >200 °C.

**$^1$H NMR (500 MHz, $CDCl_3$/$MeOD_4$, 5:1):** δ = 1.78 – 1.91 (m, 5H), 2.48 (s, 3H), 2.64 – 2.68 (m, 2H), 2.86 – 2.91 (m, 2H), 2.93 (s, 3H), 7.44 – 7.48 (m, 2H), 7.82 – 7.86 (m, 2H).

**$^{13}$C NMR (126 MHz, $CDCl_3$/$MeOD_4$, 5:1):** δ = 13.5, 23.2, 24.4, 26.8, 26.9, 42.2, 123.1, 124.3, 127.3, 128.1, 128.4, 132.9, 137.0, 138.0, 170.2, 202.8.

**HRMS (neg. APCI): [M - H]$^-$** calcd. for $C_{18}H_{19}N_2O_2$: 295.1452; found: 295.1452.

**_N,N_-Dimethyl-4-(3-methyl-4-oxo-2,4,5,6,7,8-hexahydrocyclohepta[_c_]pyrrol-1-yl)benzamide (312; RMM17)**

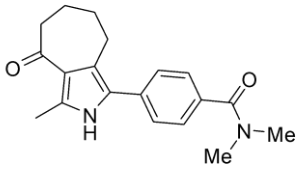

Chemical Formula: $C_{19}H_{22}N_2O_2$
Molecular Weight: 310,40

The title compound was obtained as a colourless solid (19 mg, 60 µmol, **73%**) according to the general procedure for Suzuki-Crosscoupling using 1-bromo-3-methyl-5,6,7,8-tetrahydrocyclohepta[*c*]pyrrol-4(2*H*)-one (20 mg, 83 µmol, 1.0 equiv).

**Mp:** >200 °C.

**$^1$H NMR (500 MHz, CDCl$_3$):** δ = 1.80 – 1.86 (m, 2H), 1.90 (mc, 2H), 2.59 (s, 3H), 2.68 – 2.72 (m, 2H), 2.83 – 2.88 (m, 2H), 3.03 (s, 3H), 3.14 (s, 3H), 7.31 – 7.35 (m, 2H), 7.36 – 7.39 (m, 2H), 9.00 (s, 1H).

**$^{13}$C NMR (126 MHz, CDCl$_3$):** δ = 13.7, 22.3, 23.6, 26.0, 35.6, 39.8, 41.9, 122.7, 122.8, 125.8, 127.1, 127.7, 133.9, 134.1, 135.8, 171.6, 200.2.

**HRMS (pos. ESI): [M + H]$^+$** calcd. for $C_{19}H_{23}N_2O_2$: 311.1754; found: 311.1756.

**4-(3-Methyl-4-oxo-2,4,5,6,7,8-hexahydrocyclohepta[*c*]pyrrol-1-yl)benzonitrile (313; RMM18)**

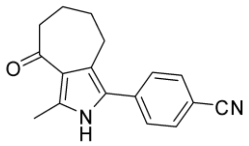

Chemical Formula: $C_{17}H_{16}N_2O$
Molecular Weight: 264,33

The title compound was obtained as a slightly yellow solid (11 mg, 41 µmol, **49%**) according to the general procedure for Suzuki-Crosscoupling using 1-bromo-3-methyl-5,6,7,8-tetrahydrocyclohepta[*c*]pyrrol-4(2*H*)-one (20 mg, 83 µmol, 1.0 equiv).

**Mp:** 129-131 ℃.

**$^1$H NMR (400 MHz, $CDCl_3$):** δ = 1.76 – 1.96 (m, 4H), 2.56 (s, 3H), 2.70 ($m_c$, 2H), 2.87 ($m_c$, 2H), 7.45 (d, *J* = 8.5 Hz, 2H), 7.68 (d, *J* = 8.5 Hz, 2H), 8.38 (s, 1H).

**$^{13}$C NMR (101 MHz, $CDCl_3$):** δ = 13.8, 22.2, 23.8, 26.0, 41.9, 109.9, 118.9, 123.3, 124.6, 124.8, 127.4 (2×), 132.7 (2×), 136.6, 137.0, 200.0.

**HRMS (pos. APCI) (*m/z*): [M+H]** calcd. for $C_{17}H_{17}N_2O$: 265.1335; found: 265.1344.

**4-Bromobenzenesulfonamide (317)**

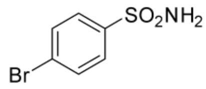

Chemical Formula: $C_6H_6BrNO_2S$
Molecular Weight: 236,08

A solution of 4-bromobenzenesulfonyl chloride (1.0 g, 3.9 mmol, 1.0 equiv) in EtOAc (3.9 mL) was added dropwise to aqueous ammonia (8.3 mL) at 0 °C. The reaction mixture was allowed to return to rt and stirred for 3 days. $H_2O$ was added and the aqueous phase was extracted three times with EtOAc. The combined organic layers were washed with $H_2O$, dried over $Na_2SO_4$ and concentrated under reduced pressure. The residue was purified by column chromatography (DCM/MeOH, 15:1) to obtain a yellow solid. The solid was digested with DCM to obtain the desired product was a colourless solid (0.62 g, 2.6 mmol, **68%**).

**$^1$H NMR (500 MHz, MeOD$_4$):** δ = 7.69 – 7.72 (m, 2H), 7.78 – 7.81 (m, 2H).

**$^{13}$C NMR (126 MHz, MeOD$_4$):** δ = 127.5, 129.0, 133.2, 144.4.

**4-(3-Methyl-4-oxo-2,4,5,6,7,8-hexahydrocyclohepta[c]pyrrol-1-yl)benzenesulfonamide (314; RMM19)**

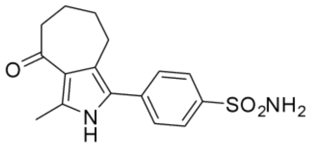

Chemical Formula: $C_{16}H_{18}N_2O_3S$
Molecular Weight: 318,39

The title compound was obtained as a colourless solid (19 mg, 41 µmol, **50%**) according to the general procedure for Suzuki-Miyaura-Crosscoupling using 1-bromo-3-methyl-5,6,7,8-tetrahydrocyclohepta[*c*]pyrrol-4(2*H*)-one (20 mg, 83 µmol, 1.0 equiv).

**Mp:** has turned brown >170 °C.

**$^1$H NMR (500 MHz, $MeOD_4$/$CDCl_3$, 3:2):** δ = 1.79 – 1.91 (m, 4H), 2.49 (s, 3H), 2.64 – 2.68 (m, 2H), 2.85 – 2.89 (m, 2H), 7.49 – 7.52 (m, 2H), 7.88 – 7.92 (m, 2H).

**$^{13}$C NMR (126 MHz, $MeOD_4$/$CDCl_3$, 3:2):** δ = 13.5, 22.9, 24.2, 26.5, 42.0, 123.0, 124.6, 126.4, 127.1, 128.1, 137.4, 138.1, 141.4, 202.4.

**HRMS (neg. APCI): [M - H]$^-$** calcd. for $C_{16}H_{17}N_2O_3S$: 317.0965; found: 317.0965.

**3-Methyl-1-(*p*-tolyl)-5,6,7,8-tetrahydrocyclohepta[*c*]pyrrol-4(2*H*)-one (315; RMM14)**

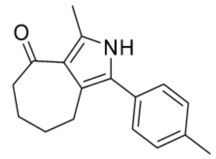

Chemical Formula: $C_{17}H_{19}NO$
Molecular Weight: 253,35

3-Methyl-1-(*p*-tolyl)-5,6,7,8-tetrahydrocyclohepta[*c*]pyrrol-4(2*H*)-one was obtained (17 mg, 66 µmol, **83%**) as a brown solid according to the general procedure for Suzuki-Crosscoupling using 1-bromo-3-methyl-5,6,7,8-tetrahydrocyclohepta[*c*]pyrrol-4(2*H*)-one (20 mg, 80 µmol, 1.0 equiv).

**Mp:** 153-155 °C.

**$^1$H NMR (500 MHz, $CDCl_3$):** δ = 1.77 – 1.92 (m, 4H), 2.38 (s, 3H), 2.55 (s, 2H), 2.66 – 2.72 (m, 2H), 2.82 – 2.88 (m, 2H), 7.19 – 7.28 (m, 4H), 8.26 (s, 1H).

**$^{13}$C NMR (126 MHz, $CDCl_3$):** δ = 13.8, 21.3, 22.4, 23.6, 26.1, 41.9, 121.6, 122.4, 126.6, 127.5, 129.6, 129.8, 134.7, 136.8, 200.5.

**HRMS (pos. ESI): $[M + H]^+$** calcd. for $C_{17}H_{20}ON$: 254.1539; found: 254.1537.

**N-(3-Bromo-4-methylphenyl)methanesulfonamide (339)**

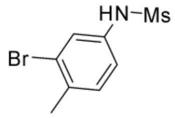

Chemical Formula: $C_8H_{10}BrNO_2S$
Molecular Weight: 264,14

Methanesulfonyl chloride (83 µL, 0.12 g, 1.1 mmol, 1.2 equiv) was added dropwise to a solution of 3-bromo-4-methylaniline (0.17 g, 0.89 mmol, 1.0 equiv) and pyridine (0.14 mL, 0.14 g, 1.8 mmol, 2.0 equiv) in dry DCM (1.8 mL). The reaction mixture was stirred at rt overnight. Aqueous HCl (0.5 M) was added, the phases separated and the aqueous phase was extracted three times with DCM. The combined organic layers were dried over $Na_2SO_4$, concentrated in vacuo and the residue was purified by column chromatography (pentane/EtOAc, 7:3) to obtain the title compound as a colourless solid (0.22 g, 0.85 mmol, **95%**).

**$^1$H NMR (400 MHz, $CDCl_3$):** δ = 2.37 (s, 3H), 3.01 (s, 3H), 6.70 (s, 1H), 7.11 (dd, J = 8.2, 2.3 Hz, 1H), 7.21 (d, J = 8.2 Hz, 1H), 7.44 (d, J = 2.3 Hz, 1H).

**$^{13}$C NMR (101 MHz, $CDCl_3$):** δ = 22.3, 39.6, 120.2, 124.9, 125.5, 131.6, 135.4, 135.5.

**N-(4-Methyl-3-(3-methyl-4-oxo-2,4,5,6,7,8-hexahydrocyclohepta[*c*]pyrrol-1-yl)phenyl)methanesulfonamide (329; RMM20)**

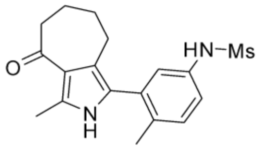

Chemical Formula: $C_{18}H_{22}N_2O_3S$
Molecular Weight: 346,45

The title compound was obtained as a brown oil (24 mg, 70 µmol, **85%**) according to the general procedure for Suzuki-Miyaura-Crosscoupling using 1-bromo-3-methyl-5,6,7,8-tetrahydrocyclohepta[*c*]pyrrol-4(2*H*)-one (20 mg, 83 µmol, 1.0 equiv).

**$^1$H NMR (400 MHz, Acetone-*d*$_6$):** δ = 1.72 – 1.86 (m, 4H), 2.21 (s, 3H), 2.47 (d, J = 0.5 Hz, 3H), 2.58 (dd, J = 12.0, 5.7 Hz, 4H), 2.96 (s, 3H), 7.21 – 7.31 (m, 3H), 8.46 (s, 1H), 10.26 (s, 1H

**$^{13}$C NMR (101 MHz, Acetone-*d*$_6$):** δ = 13.4, 19.7, 22.8, 24.0, 26.4, 39.3, 42.3, 121.0, 121.8, 122.3, 122.7, 124.0, 126.1, 131.9, 134.5, 135.0, 136.9, 199.1.

**HRMS (neg. ESI): $[M - H]^-$** calcd. for $C_{18}H_{21}N_2O_3S$: 345.1278; found: 345.1279.

##### *N*-(3-Bromo-2-methylphenyl)methanesulfonamide (340)

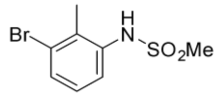

Chemical Formula: $C_8H_{10}BrNO_2S$
Molecular Weight: 264,14

*N*-(3-Bromo-2-methylphenyl)methanesulfonamide was prepared ajar to literature: [25]

Methanesulfonyl chloride (75 µL, 0.11 g, 0.97 mmol, 1.2 equiv) was added dropwise to a solution of 3-bromo-2-methylaniline (99 µL, 0.15 g, 0.81 mmol, 1.0 equiv) and pyridine (0.13 mL, 0.13 g, 1.6 mmol, 2.0 equiv) in DCM (1.6 mL) at rt. The reaction mixture was stirred at rt overnight. Aqueous HCl (0.5 M) and DCM were added. The phases were separated and the aqueous phase was extracted three times with DCM. The combined organic layers were washed with brine, dried over $Na_2SO_4$ and evaporated. The residue was purified by column chromatography (pentane/EtOAc, 4:1). The title compound was obtained as a colorless solid (0.18 g, 0.71 mmol, **88%**).

**$^1$H NMR (300 MHz, $CDCl_3$):** δ = 2.44 (s, 3H), 3.03 (s, 3H), 6.30 (br. s, 1H), 7.10 (t, *J* = 8.0 Hz, 1H), 7.43 (dd, *J* = 8.1, 1.2 Hz, 1H), 7.47 (dd, *J* = 8.1, 1.2 Hz, 1H) ppm.

**N-(2-Methyl-3-(3-methyl-4-oxo-2,4,5,6,7,8-hexahydrocyclohepta[*c*]pyrrol-1-yl)phenyl)methanesulfonamide (332; RMM21)**

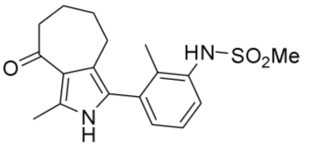

Chemical Formula: $C_{18}H_{22}N_2O_3S$
Molecular Weight: 346,45

The title compound was obtained as a yellow oil (7.1 mg, 21 µmol, **25%**) according to the general procedure for Suzuki-Miyaura-Crosscoupling using 1-bromo-3-methyl-5,6,7,8-tetrahydrocyclohepta[*c*]pyrrol-4(2*H*)-one (20 mg, 83 µmol, 1.0 equiv).

**$^1$H NMR (400 MHz, MeOD-$d_4$):** δ = 1.76 ($m_c$, 2H), 1.85 ($m_c$, 2H), 2.23 (s, 3H), 2.46 (s, 3H), 2.54 ($m_c$, 2H), 2.64 ($m_c$, 2H), 3.01 (s, 3H), 7.14 (dd, *J* = 7.6, 1.4 Hz, 1H), 7.24 (dd, *J* = 7.8, 7.7 Hz, 1H), 7.37 (dd, *J*=8.0, 1.4 Hz, 1H).

**$^{13}$C NMR (101 MHz, MeOD-$d_4$):** δ = 13.6, 16.2, 23.3, 24.2, 26.5, 40.3, 42.4, 121.6, 123.6, 127.0, 127.0, 127.2, 130.8, 135.5, 135.9, 136.9, 137.2, 203.0.

**HRMS (pos. ESI) (*m/z*): [M+H]$^+$** calcd. for $C_{18}H_{23}N_2O_3S$: 347.1424; found: 347.1427.

**3-Methyl-4-(3-methyl-4-oxo-2,4,5,6,7,8-hexahydrocyclohepta[*c*]pyrrol-1-yl)benzamide (335; RMM23)**

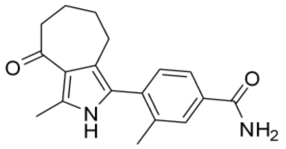

Chemical Formula: $C_{18}H_{20}N_2O_2$
Molecular Weight: 296,37

The title compound was obtained as a colourless solid (19 mg, 64 µmol, **78%**) according to the general procedure for Suzuki-Miyaura-Crosscoupling using 1-bromo-3-methyl-5,6,7,8-tetrahydrocyclohepta[*c*]pyrrol-4(2*H*)-one (20 mg, 83 µmol, 1.0 equiv).

**Mp:** >200 °C.

**$^1$H NMR (500 MHz, $MeOD_4/CDCl_3$, 2:1):** δ = 1.75 ($m_c$, 2H), 1.83 – 1.89 (m, 2H), 2.28 (s, 3H), 2.47 (s, 3H), 2.53 – 2.57 (m, 2H), 2.61 – 2.66 (m, 2H), 7.26 (d, *J* = 7.9 Hz, 1H), 7.66 – 7.70 (m, 1H), 7.76 – 7.78 (m, 1H).

**$^{13}$C NMR (126 MHz, $MeOD_4/CDCl_3$, 2:1):** δ = 13.6, 20.4, 22.9, 24.0, 26.2, 42.1, 121.5, 123.7, 125.4, 126.2, 130.2, 131.8, 133.4, 137.0, 137.0, 138.7, 171.8, 202.7.

**HRMS (pos. APCI): $[M + H]^+$** calcd. for $C_{18}H_{21}N_2O_2$: 297.1598; found: 297.1598.

**3-Methyl-4-(3-methyl-4-oxo-2,4,5,6,7,8-hexahydrocyclohepta[*c*]pyrrol-1-yl)benzonitrile (336; RMM22)**

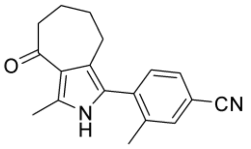

Chemical Formula: $C_{18}H_{18}N_2O$
Molecular Weight: 278,36

3-Methyl-4-(3-methyl-4-oxo-2,4,5,6,7,8-hexahydrocyclohepta[*c*]pyrrol-1-yl)benzonitrile was obtained as a colorless wax (6.4 mg, 23 µmol, **28%**) according to the general procedure for Suzuki-Miyaura-Crosscoupling using 1-bromo-3-methyl-5,6,7,8-tetrahydrocyclohepta[*c*]pyrrol-4(2*H*)-one (20 mg, 83 µmol, 1.0 equiv).

**$^{1}$H NMR (400 MHz, $CDCl_3$):** δ = 1.67 – 1.97 (m, 4H), 2.31 (s, 3H), 2.44 – 2.60 (m, 5H), 2.69 ($m_c$, 2H), 7.31 (d, *J* = 7.9 Hz, 1H), 7.51 ($m_c$, 1H), 7.56 – 7.57 (m, 1H), 7.97 (s, 1H).

**$^{13}$C NMR (101 MHz, $CDCl_3$):** δ = 13.8, 20.2, 22.2, 23.7, 25.6, 42.0, 111.7, 118.8, 121.9, 123.7, 123.9, 129.5, 131.6, 134.1, 135.5, 137.0, 138.8, 200.1.

**HRMS (neg. ESI) (*m/z*): $[M-H]^-$** calcd. for $C_{18}H_{17}N_2O$: 277.1346; found: 277.1349.

**4-Bromo-3,5-dimethylbenzamide (352)**

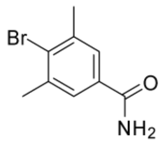

Chemical Formula: $C_9H_{10}BrNO$
Molecular Weight: 228,09

4-Bromo-3,5-dimethylbenzonitrile (0.10 g, 0.48 mmol, 1.0 equiv) in $H_2SO_4$ (0.95 mL) was stirred at rt for 4 h. The reaction mixture was poured on ice cooled $H_2O$. EtOAc was added and the phases were separated and the aqueous phase was extracted with EtOAc three times. The combined organic layers were washed with brine, dried over $Na_2SO_4$ and evaporated. The residue was purified by column chromatography (pentane/EtOAc, 5:1 up to 1:1). The title compound was obtained as a colorless solid (92 mg, 0.40 mmol, **83%**).

**$^1$H NMR (300 MHz, $CDCl_3$):** δ = 2.47 (s, 6H), 7.51 (s, 2H) ppm.

Analytical data matches literature.[26]

**3,5-dimethyl-4-(3-methyl-4-oxo-2,4,5,6,7,8-hexahydrocyclohepta[c]pyrrol-1-yl)benzonitrile (356)**

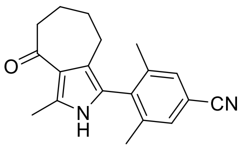

Chemical Formula: $C_{19}H_{20}N_2O$
Molecular Weight: 292,38

The title compound was obtained as a yellow wax (4.1 mg, 14 µmol, **17%**) according to the general procedure for Suzuki-Miyaura-Crosscoupling using 1-bromo-3-methyl-5,6,7,8-tetrahydrocyclohepta[*c*]pyrrol-4(2*H*)-one (20 mg, 83 µmol, 1.0 equiv).

**$^1$H NMR (400 MHz, CDCl$_3$):** δ = 1.71 – 1.79 (m, 2H), 1.80 – 1.91 (m, 3H), 2.14 (t, J = 0.6 Hz, 6H), 2.36 – 2.41 (m, 2H), 2.55 (s, 3H), 2.65 – 2.71 (m, 2H), 7.37 – 7.42 (m, 2H), 7.82 (s, 1H).

**$^{13}$C NMR (101 MHz, CDCl$_3$):** δ = 13.8, 20.3, 22.3, 23.4, 25.5, 42.2, 112.2, 118.9, 121.4, 121.8, 122.7, 130.7, 135.2, 137.0, 140.9, 200.3.

**HRMS (neg. ESI) (*m/z*): [M−H]$^-$** calcd. for $C_{19}H_{19}N_2O$: 291.1503; found: 291.1504.

**3,5-Dimethyl-4-(3-methyl-4-oxo-2,4,5,6,7,8-hexahydrocyclohepta[c]pyrrol-1-yl)benzamide (355; RMM25)**

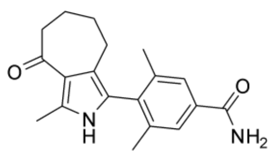

Chemical Formula: $C_{19}H_{22}N_2O_2$
Molecular Weight: 310,40

The title compound was obtained as a colourless solid (9.0 mg, 29 µmol, **24%**) according to the general procedure for Suzuki-Miyaura-Crosscoupling using 1-bromo-3-methyl-5,6,7,8-tetrahydrocyclohepta[*c*]pyrrol-4(2*H*)-one (20 mg, 83 µmol, 1.0 equiv) except that for the borylation 10% DMSO were added and the reaction was stirred at 100 °C for 44 h and for the Suzuki-coupling the reaction conditions were altered to 80 °C overnight.

**Mp:** 131-133 ℃.

**$^1$H NMR (300 MHz, $CDCl_3$/MeOD-*d*$_4$ 2:1):** δ = 1.70 ($m_c$, 2H), 1.82 ($m_c$, 2H), 2.10 (s, 6H), 2.36 ($m_c$, 2H), 2.46 (s, 3H), 7.53 (s, 2H).

**$^{13}$C NMR (126 MHz, $CDCl_3$/MeOD-*d*$_4$ 2:1):** δ = 13.7, 20.5 (2×), 22.5, 22.6, 23.6, 25.6, 30.0, 78.0, 120.8, 122.6, 123.9, 126.5, 133.1, 136.5, 140.1, 171.5, 202.4.

**HRMS (neg. ESI) (*m/z*): [M−H]$^-$** calcd. for $C_{19}H_{21}N_2O_2$: 309.1609; found: 309.1609.

**NMR Spectra**

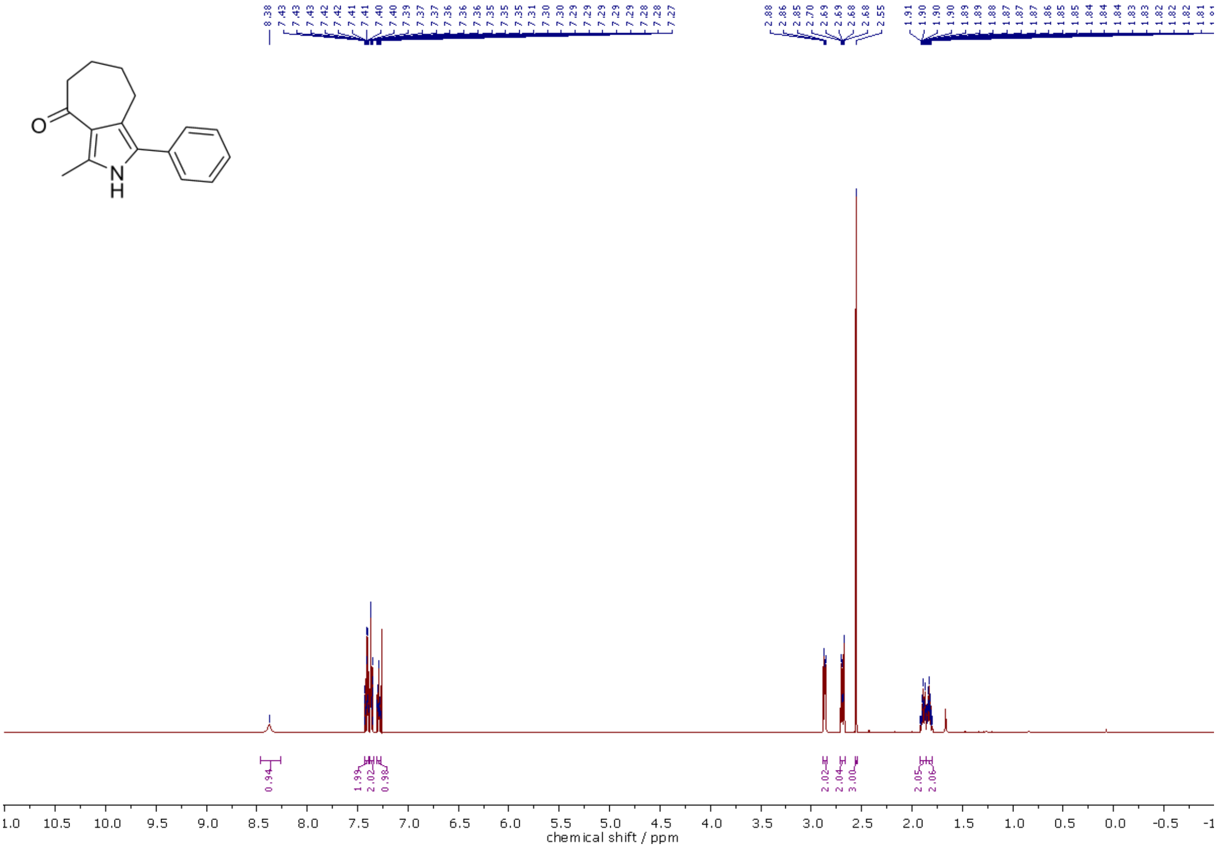

[1]H NMR (500 MHz, $CDCl_3$) spectra of **208** MPM3.

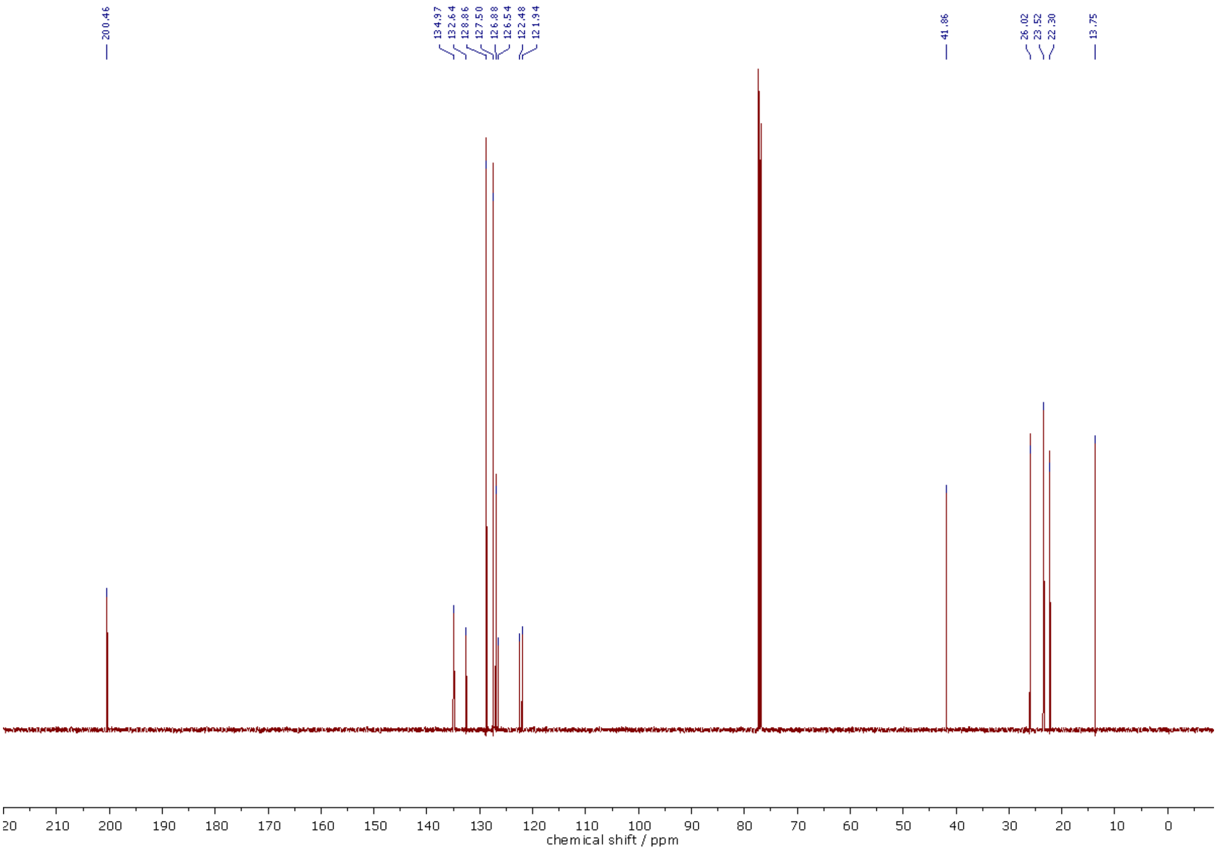

[13]C NMR (126 MHz, $CDCl_3$) spectra of **208** MPM3.

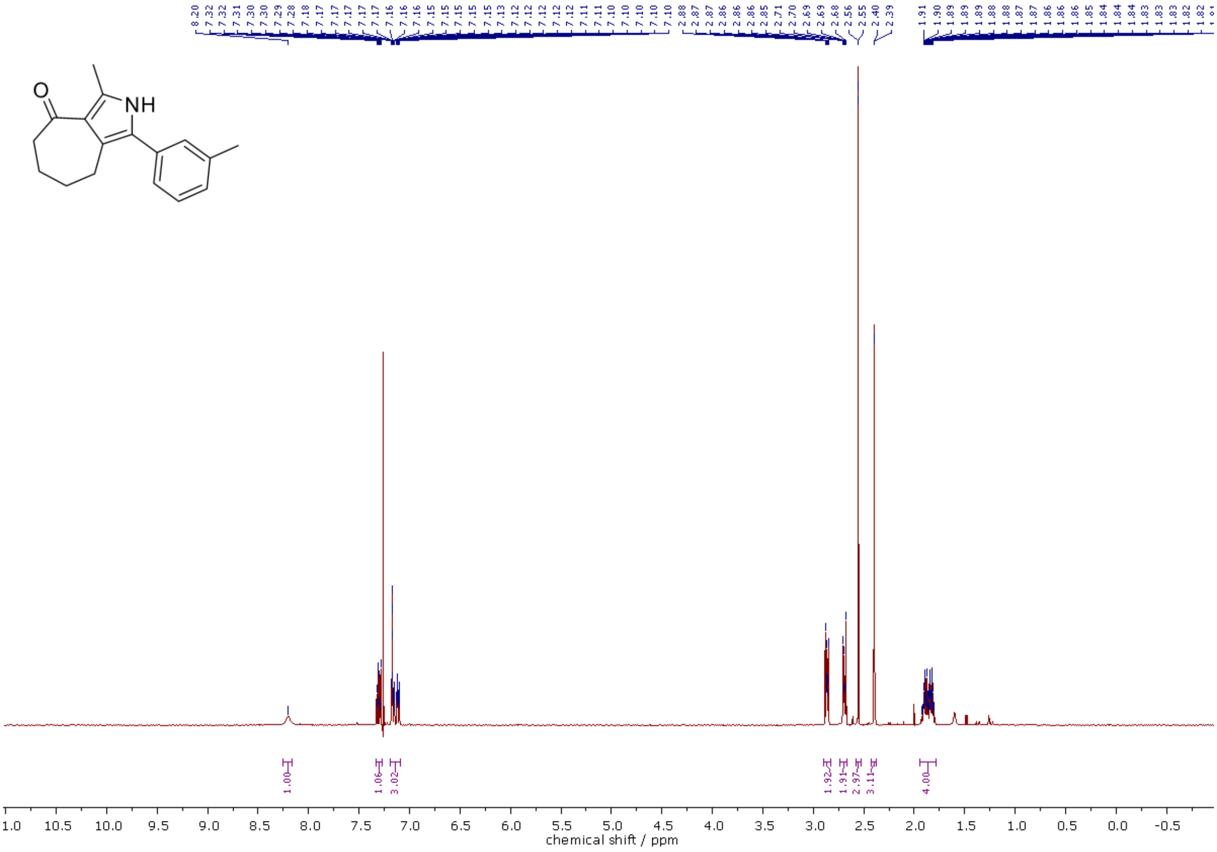

[1]H NMR (400 MHz, $CDCl_3$) spectra of **292**.

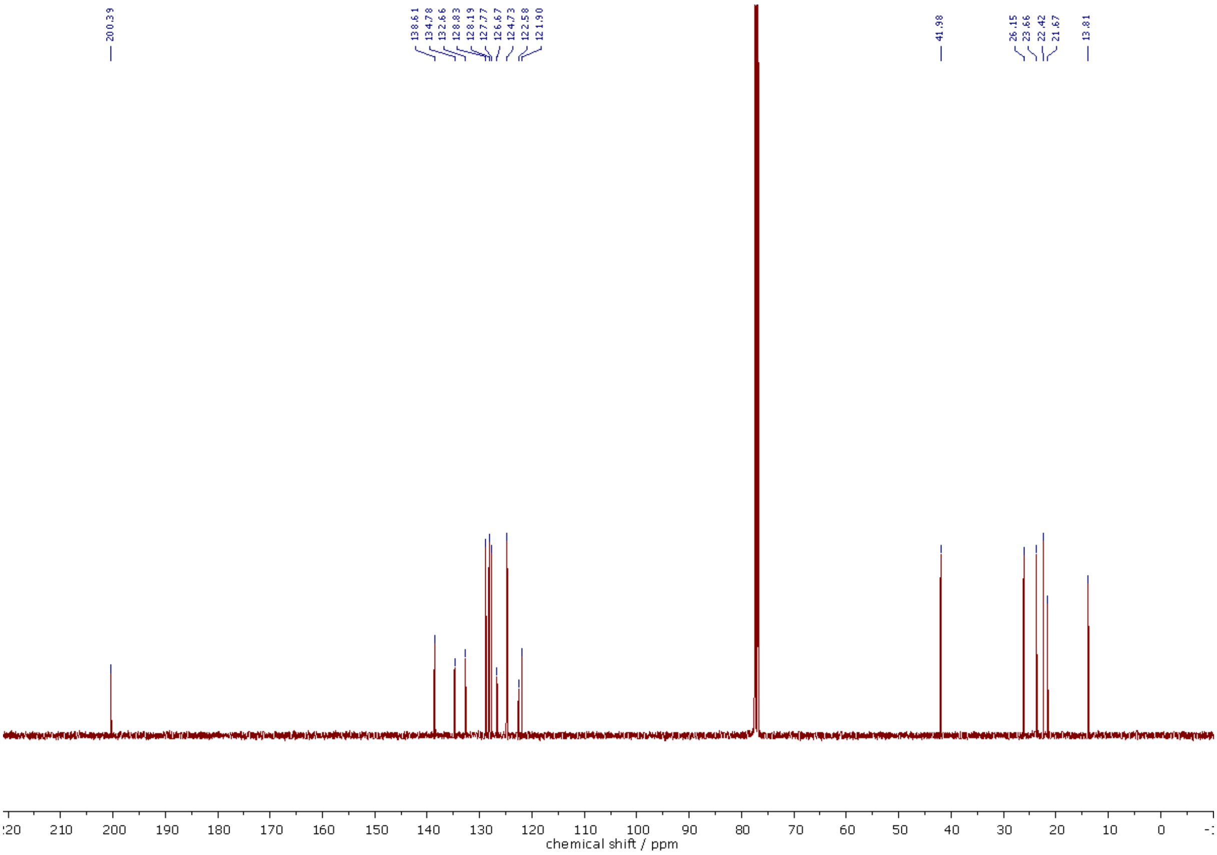

$^{13}C$ NMR (101 MHz, $CDCl_3$) spectra of **292**.

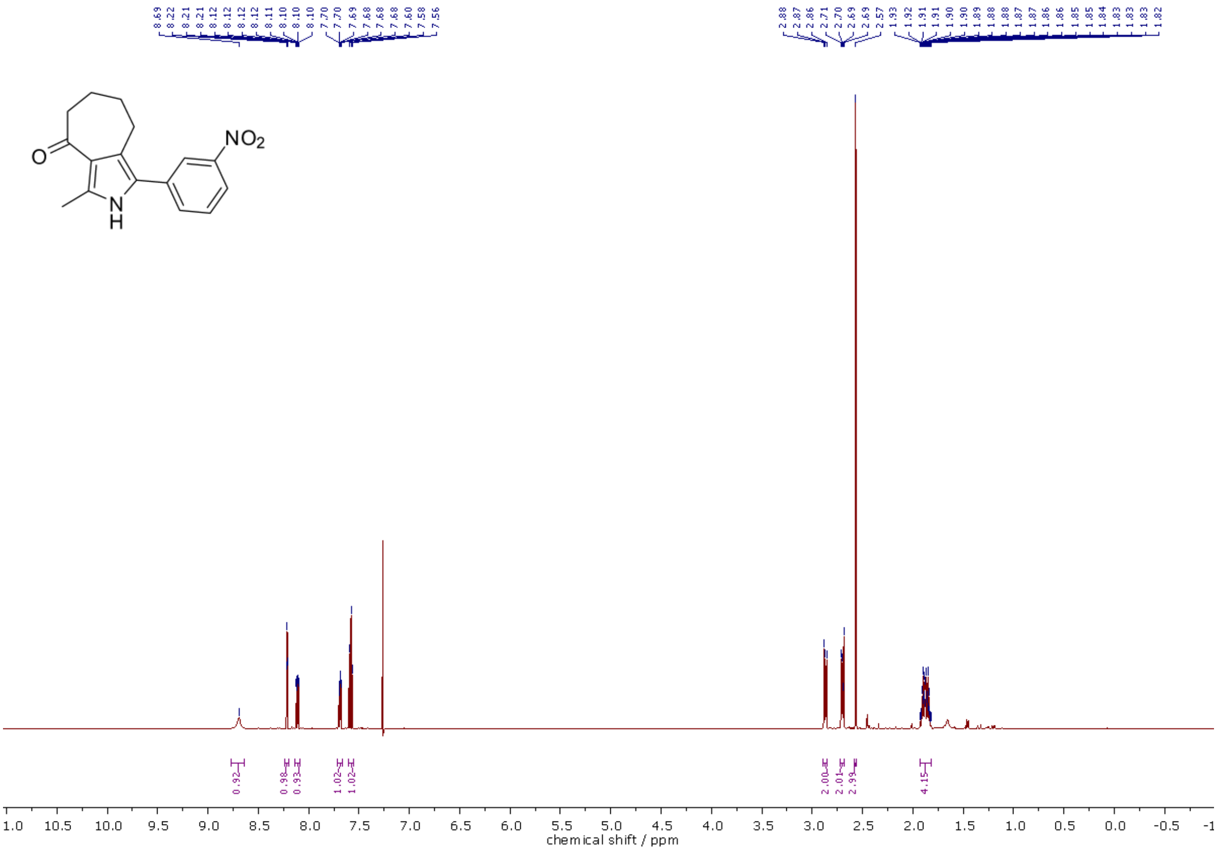

$^{1}H$ NMR (500 MHz, $CDCl_3$) spectra of **293**.

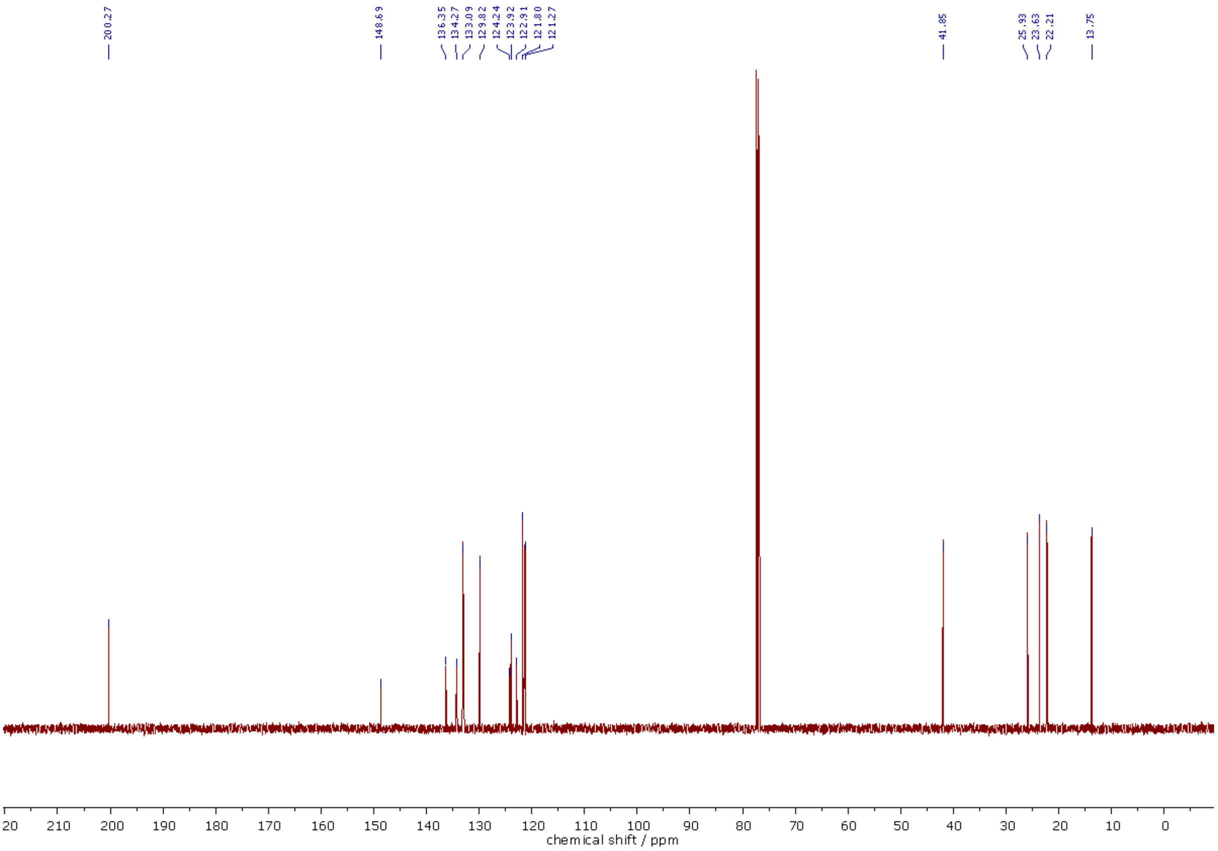

$^{13}$C NMR (126 MHz, $CDCl_3$) spectra of **293**.

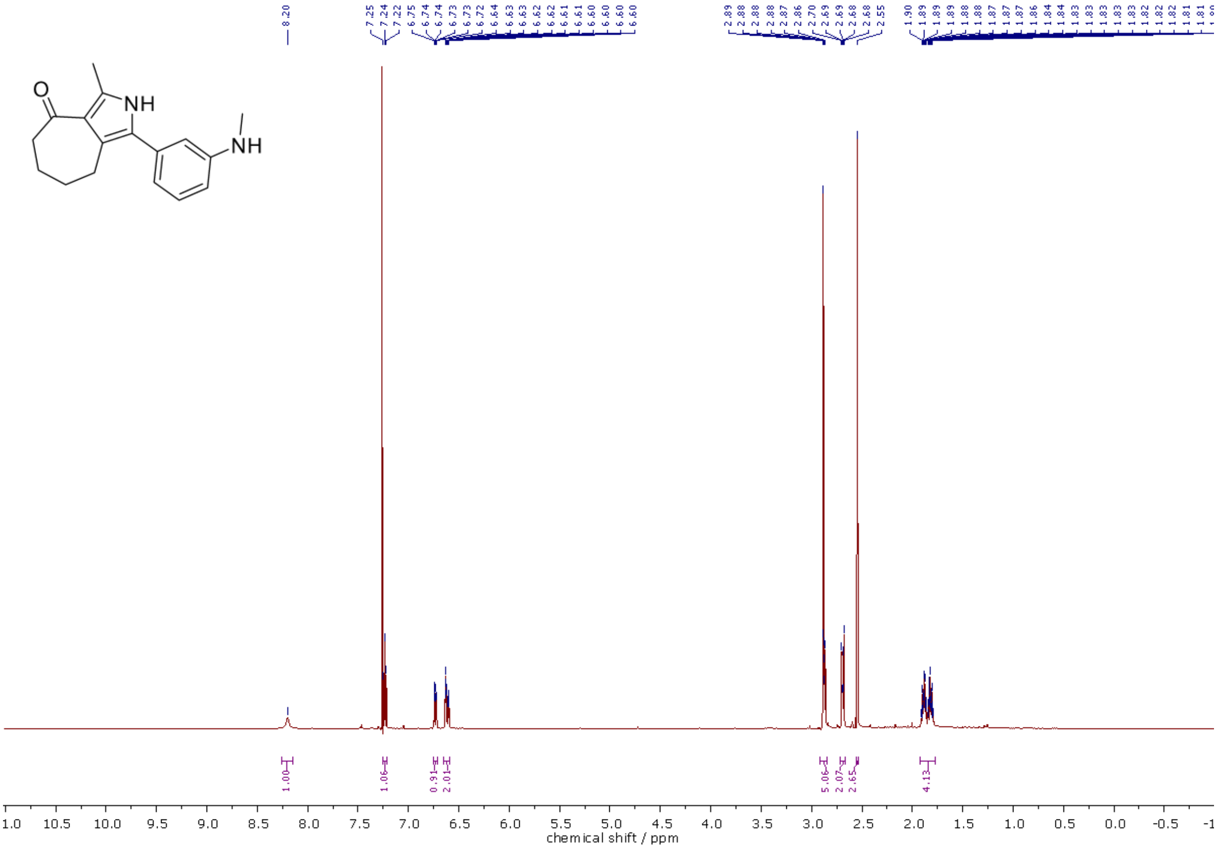

$^{1}$H NMR (500 MHz, $CDCl_3$) spectra of **294**.

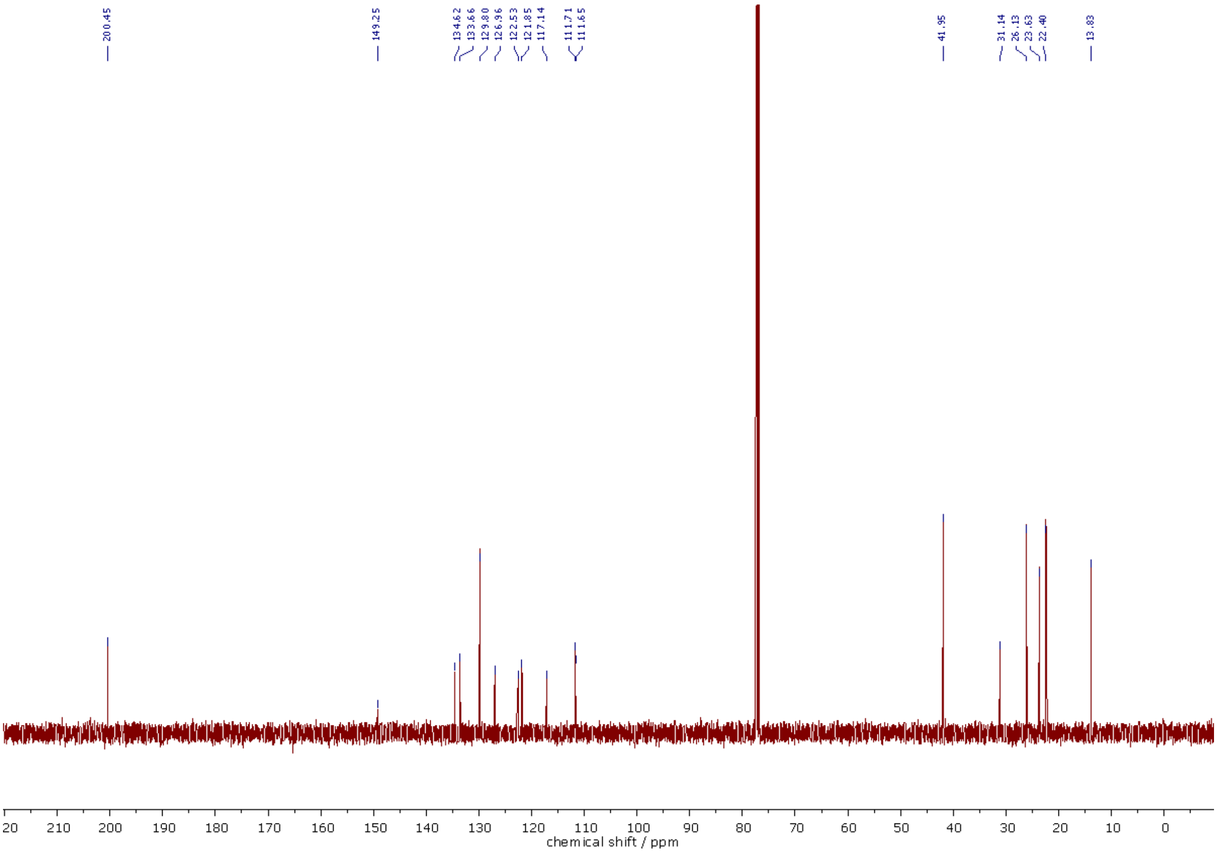

[13]C NMR (126 MHz, $CDCl_3$) spectra of **294**.

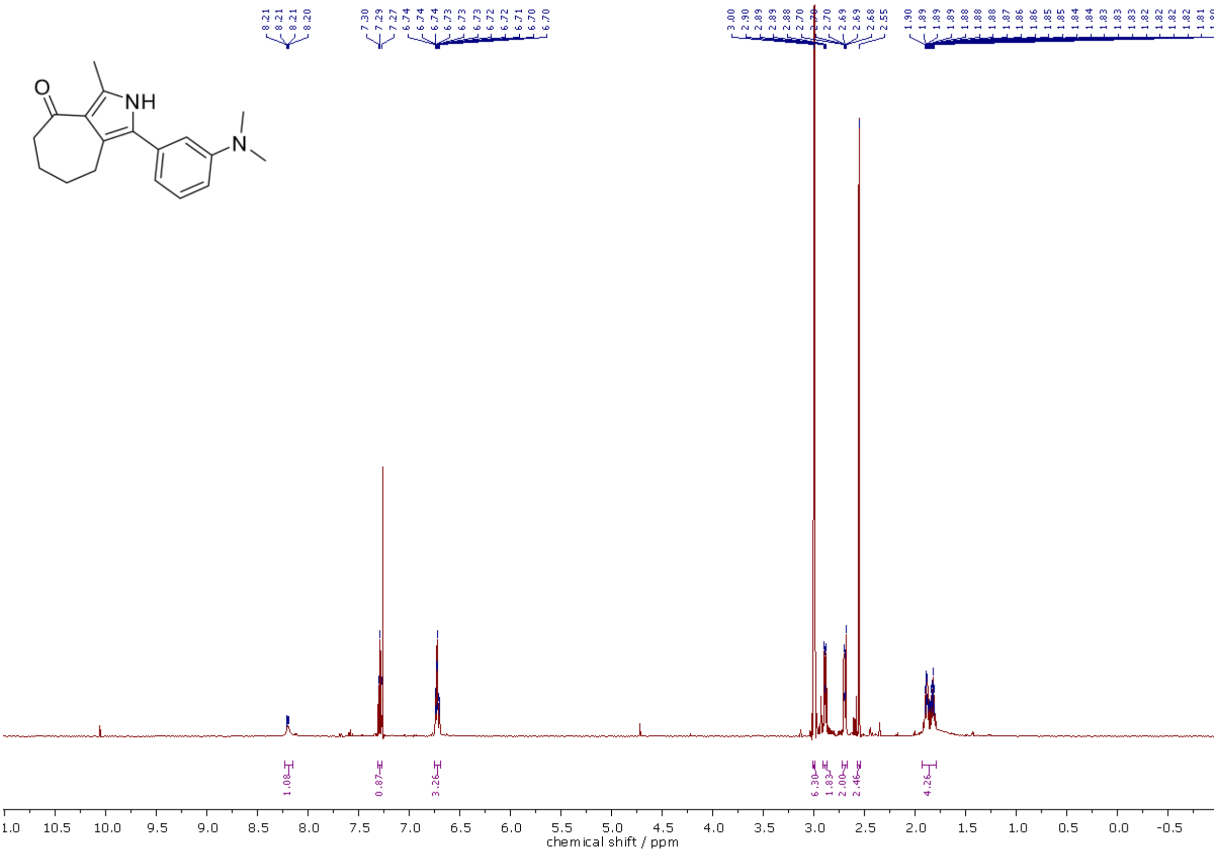

[1]H NMR (500 MHz, $CDCl_3$) spectra of **295**.

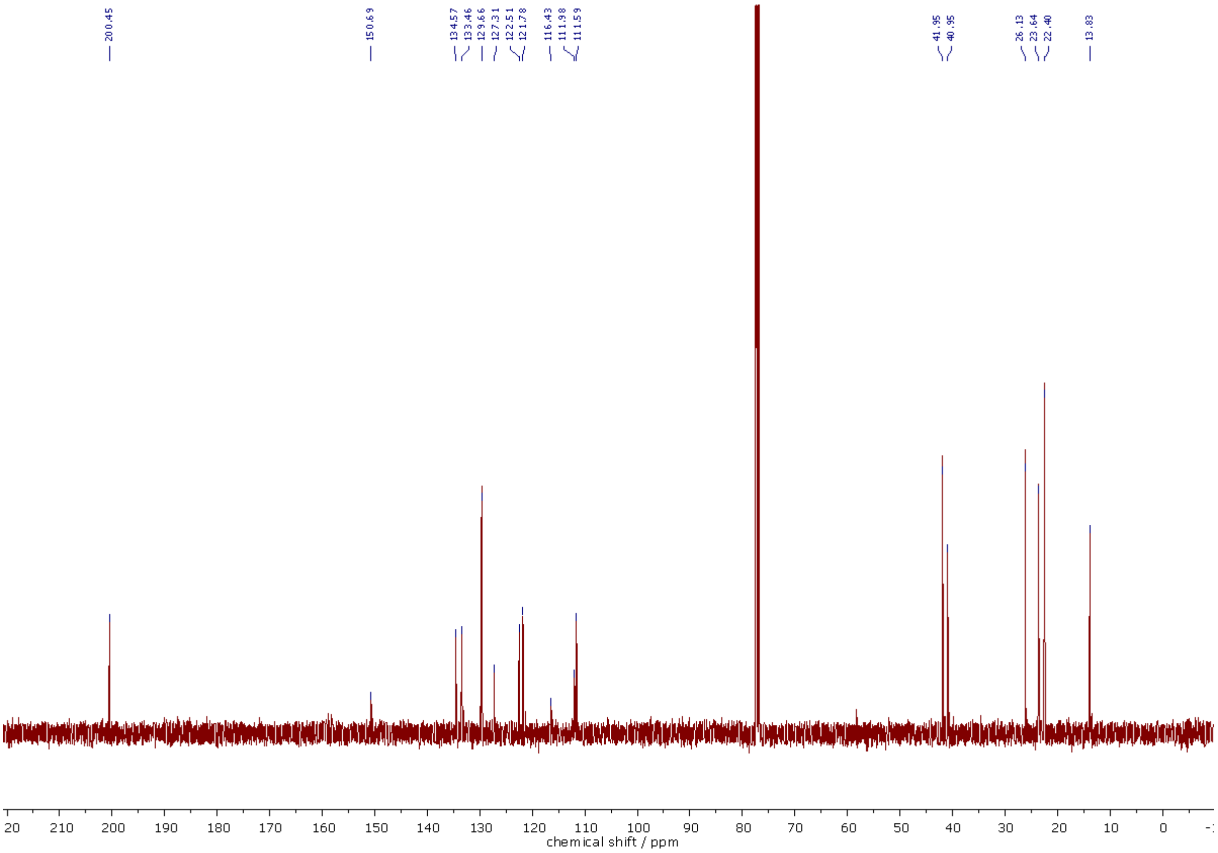

$^{13}$C NMR (126 MHz, $CDCl_3$) spectra of **295**.

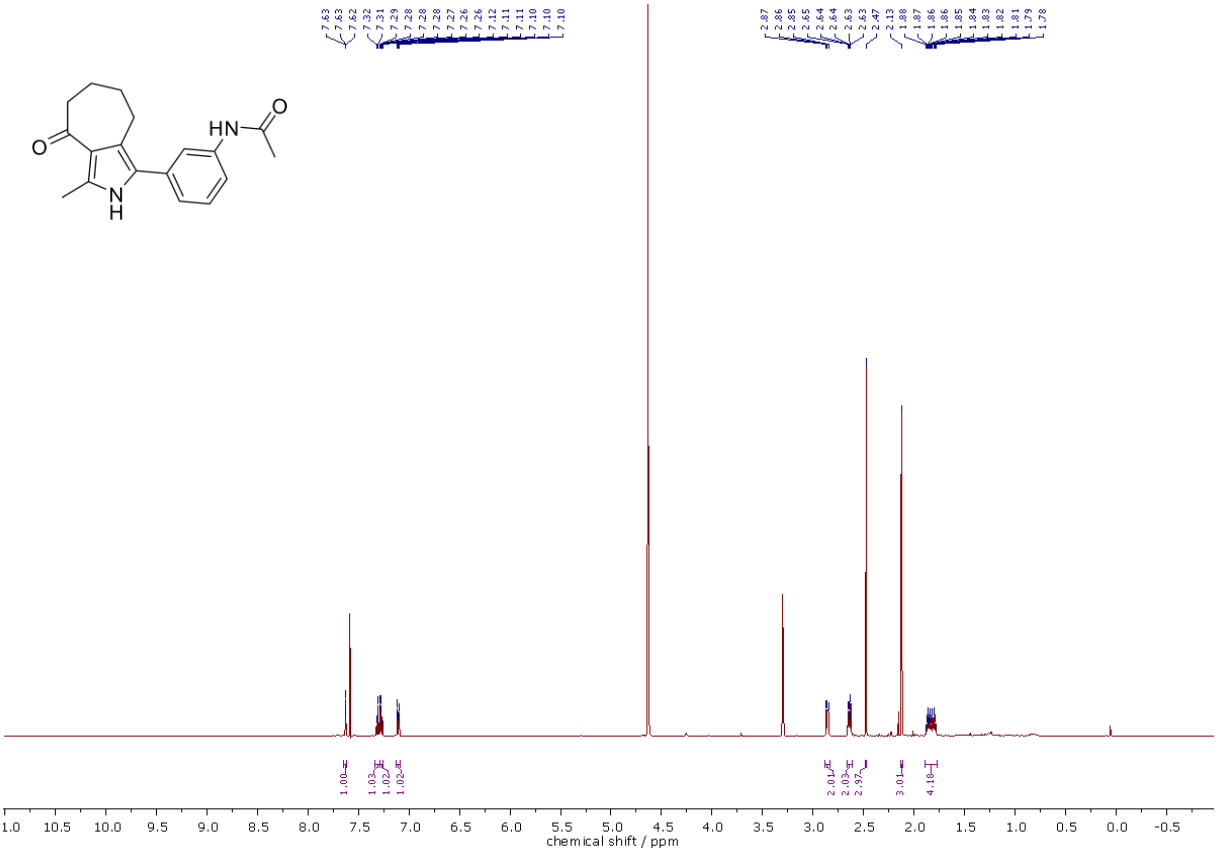

$^{1}$H NMR (500 MHz, MeOD-$d_4$/$CDCl_3$, 1:1) spectra of **296**.

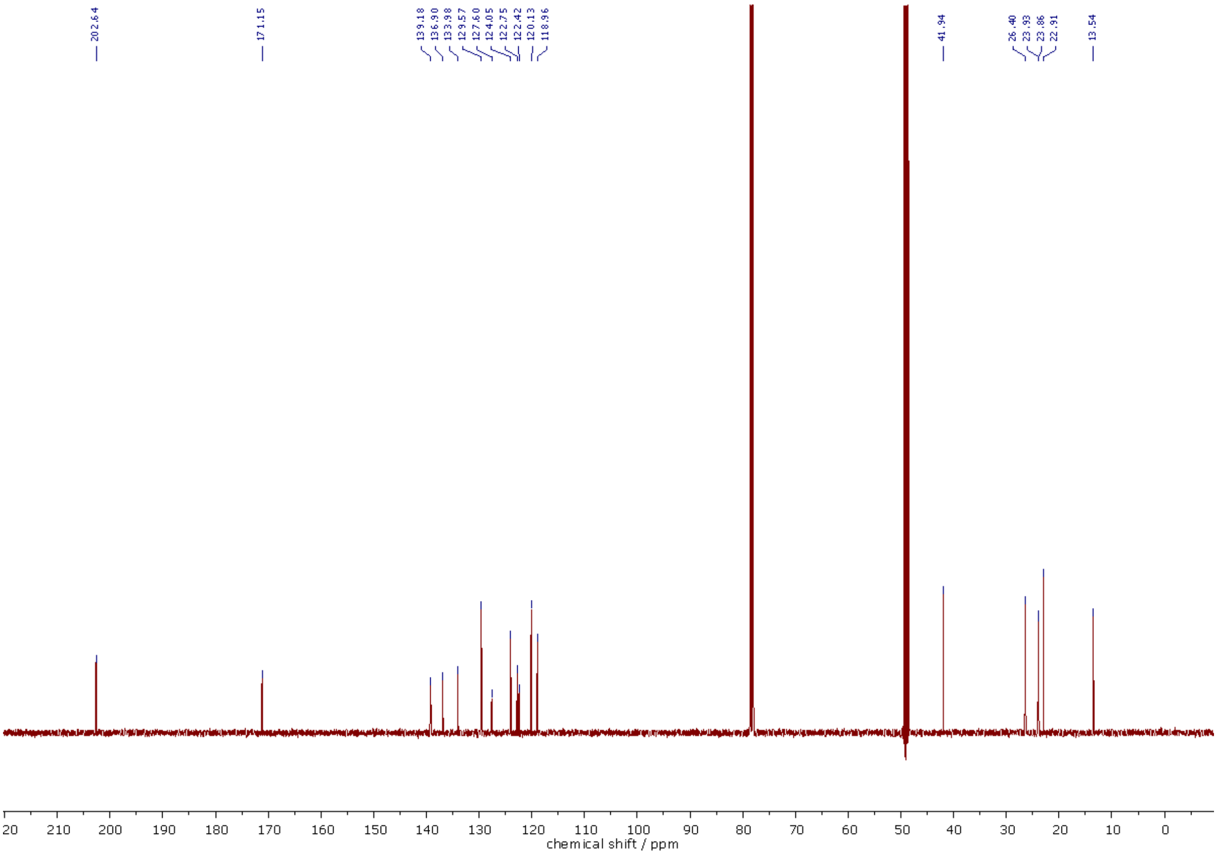

$^{13}C$ NMR (126 MHz, MeOD-d$_4$/CDCl$_3$, 1:1) spectra of **296**.

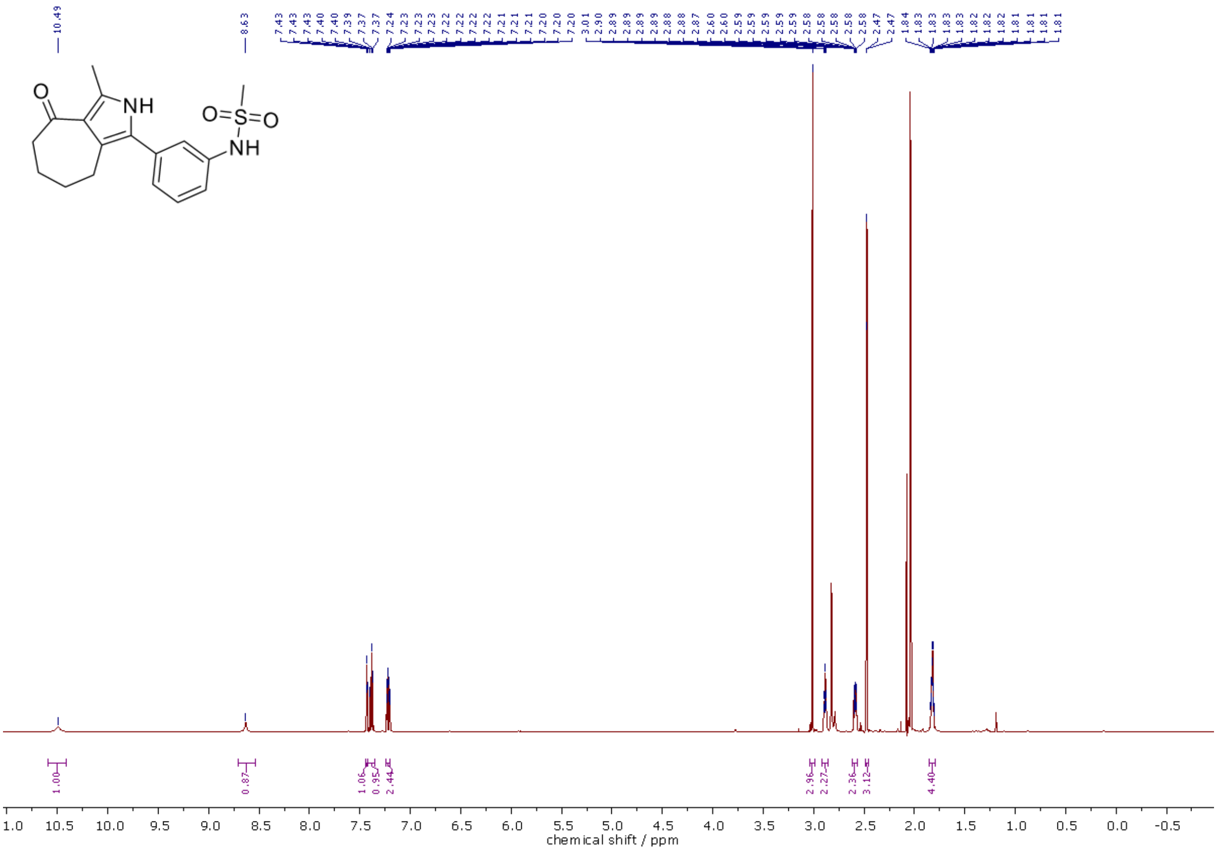

$^{1}H$ NMR (500 MHz, Acetone-d$_6$) spectra of **297**.

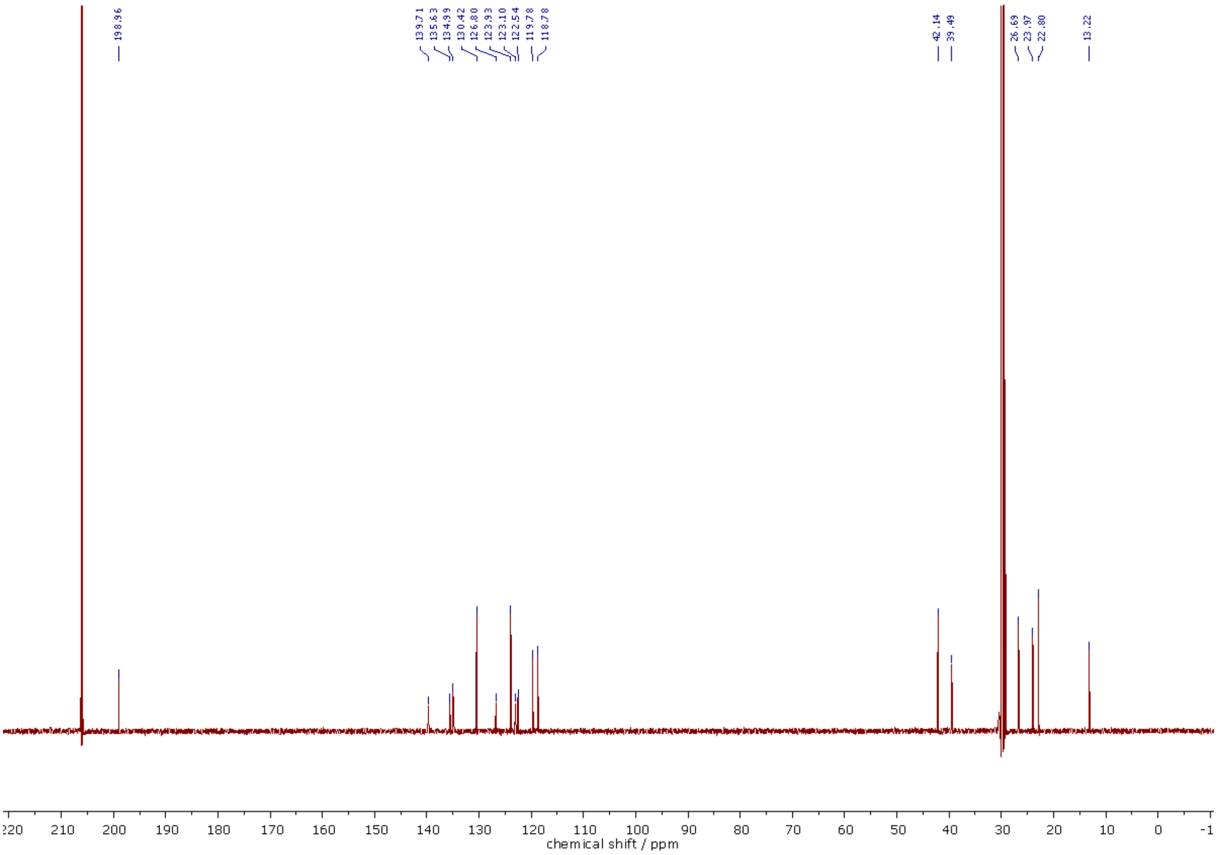

$^{13}C$ NMR (126 MHz, Acetone-$d_6$) spectra of **297**.

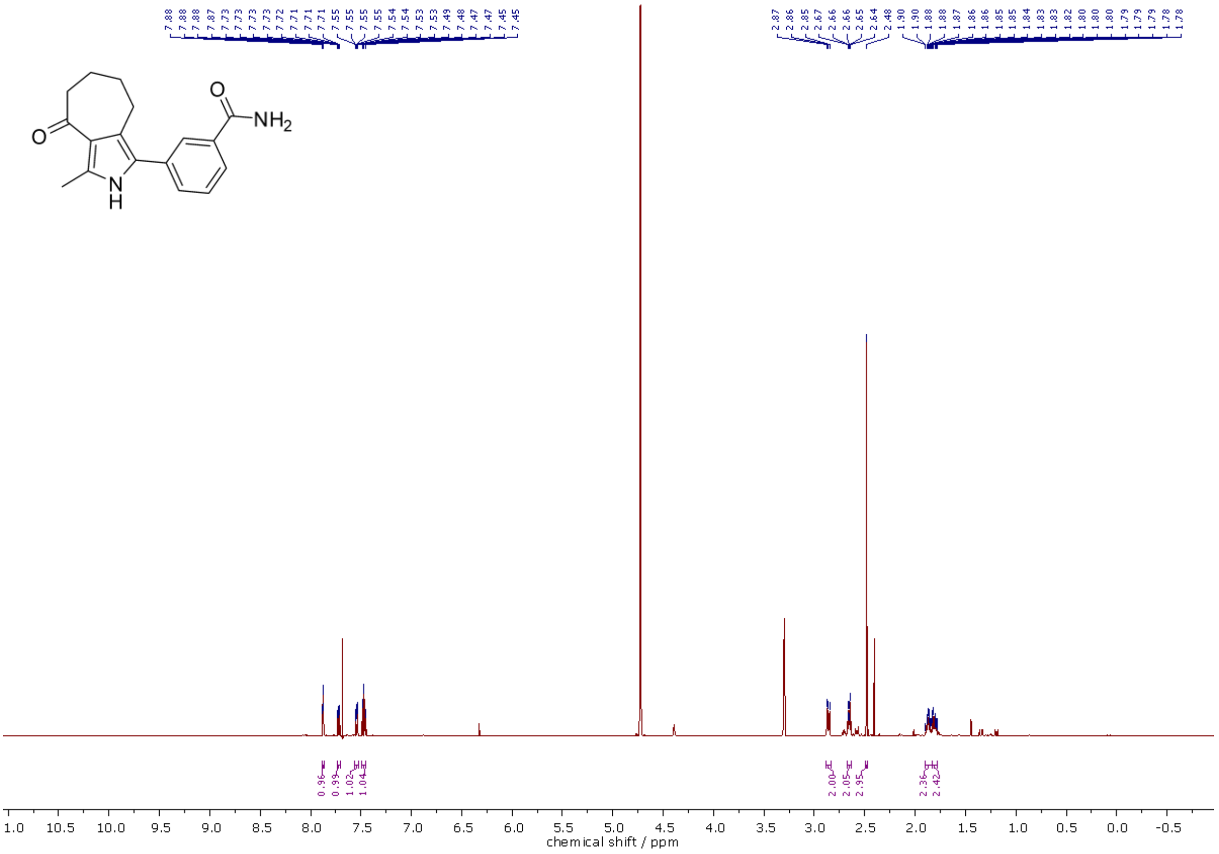

$^{1}H$ NMR (500 MHz, MeOD-$d_4$/$CDCl_3$, 2:1) spectra of **298**.

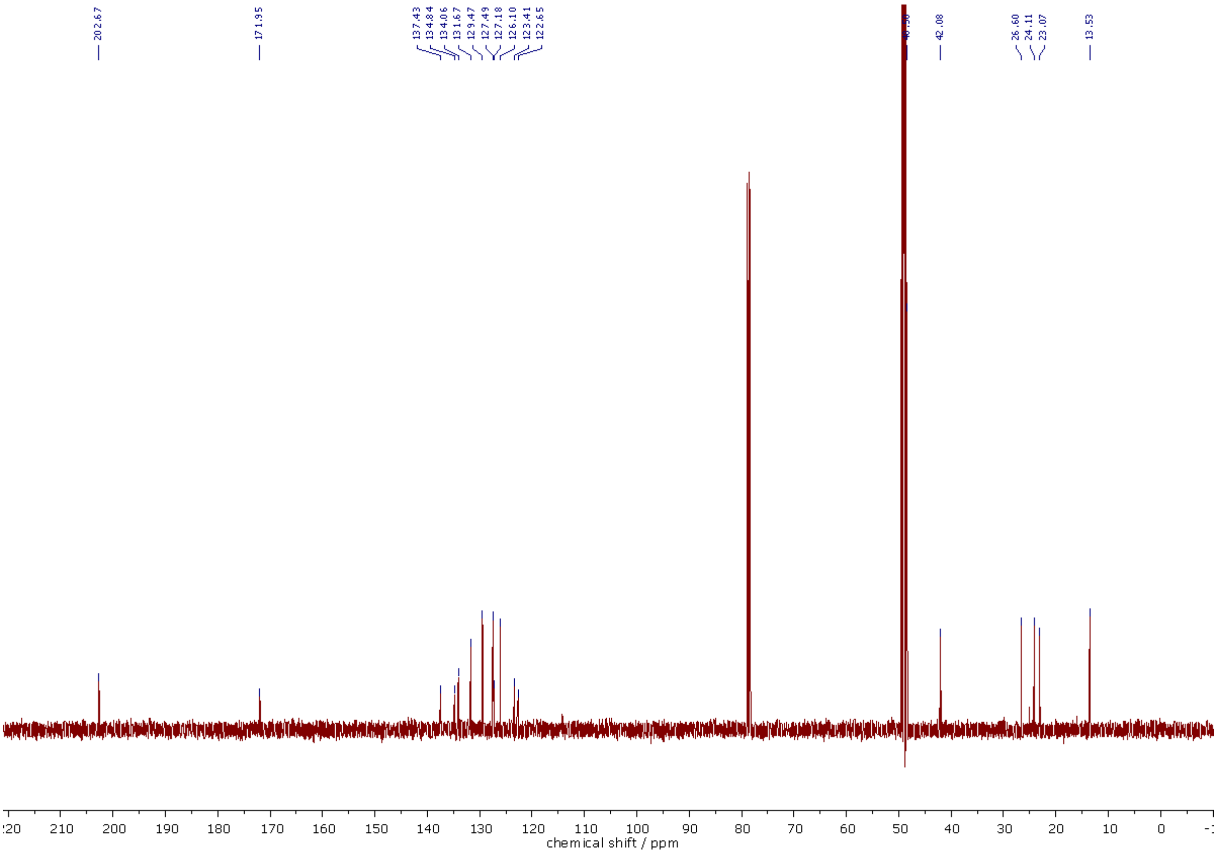

$^{13}C$ NMR (126 MHz, MeOD-d$_4$/$CDCl_3$, 2:1) spectra of **298**.

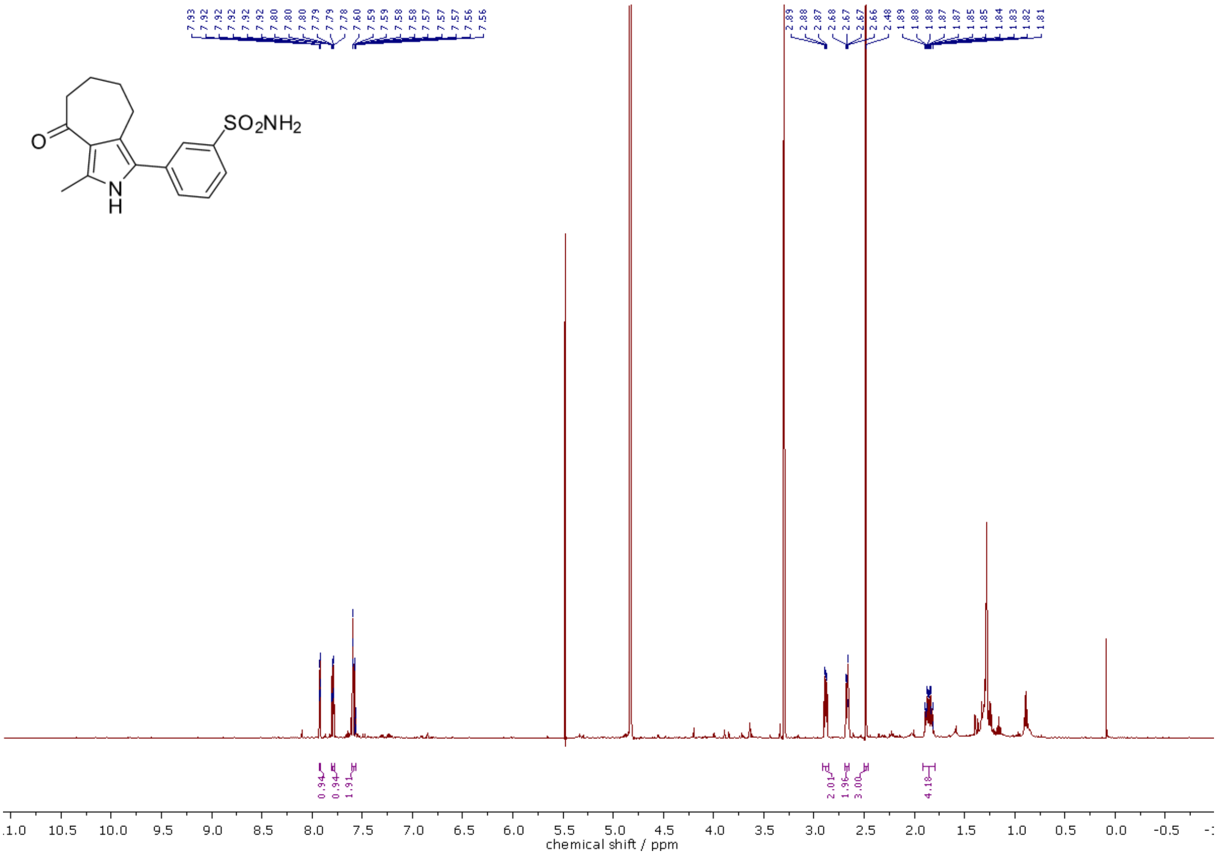

$^{1}H$ NMR (500 MHz, MeOD-d$_4$) spectra of **299**.

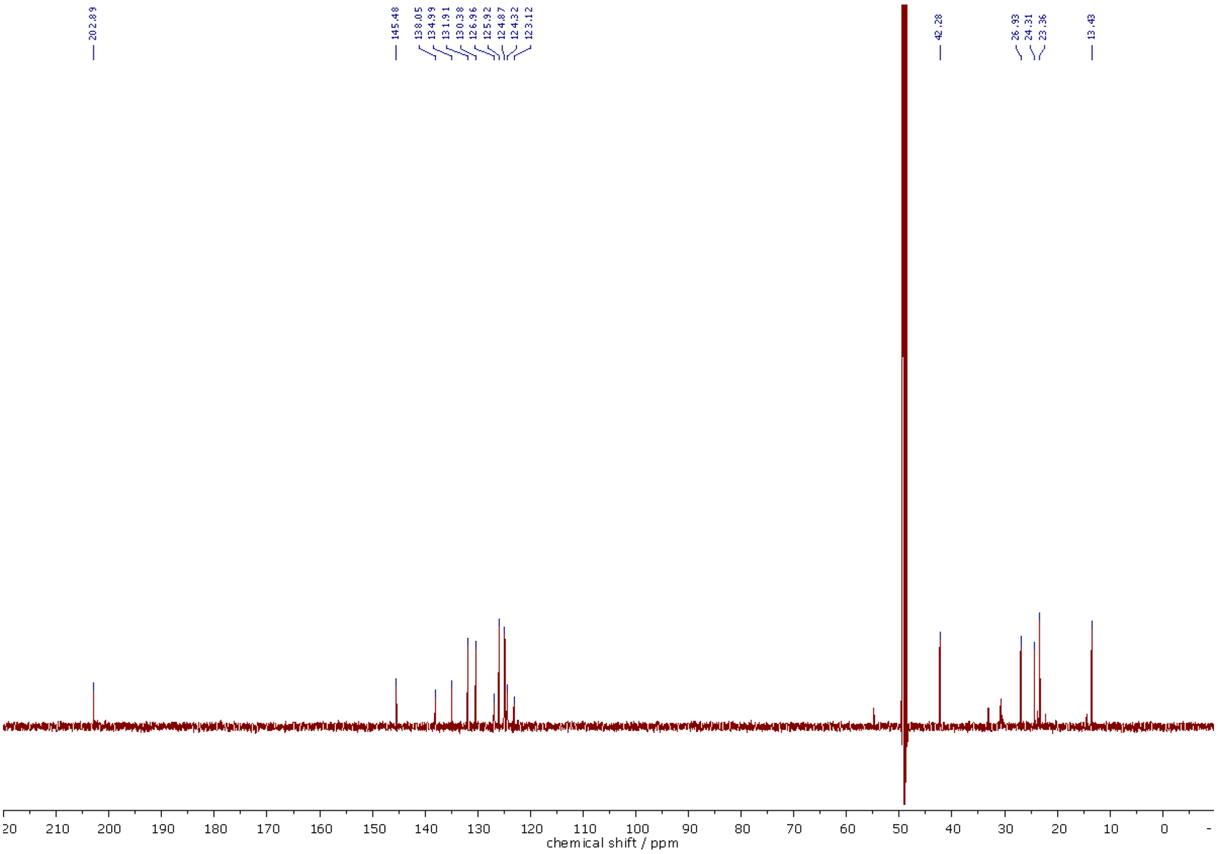

$^{13}C$ NMR (126 MHz, MeOD-$d_4$) spectra of **299**.

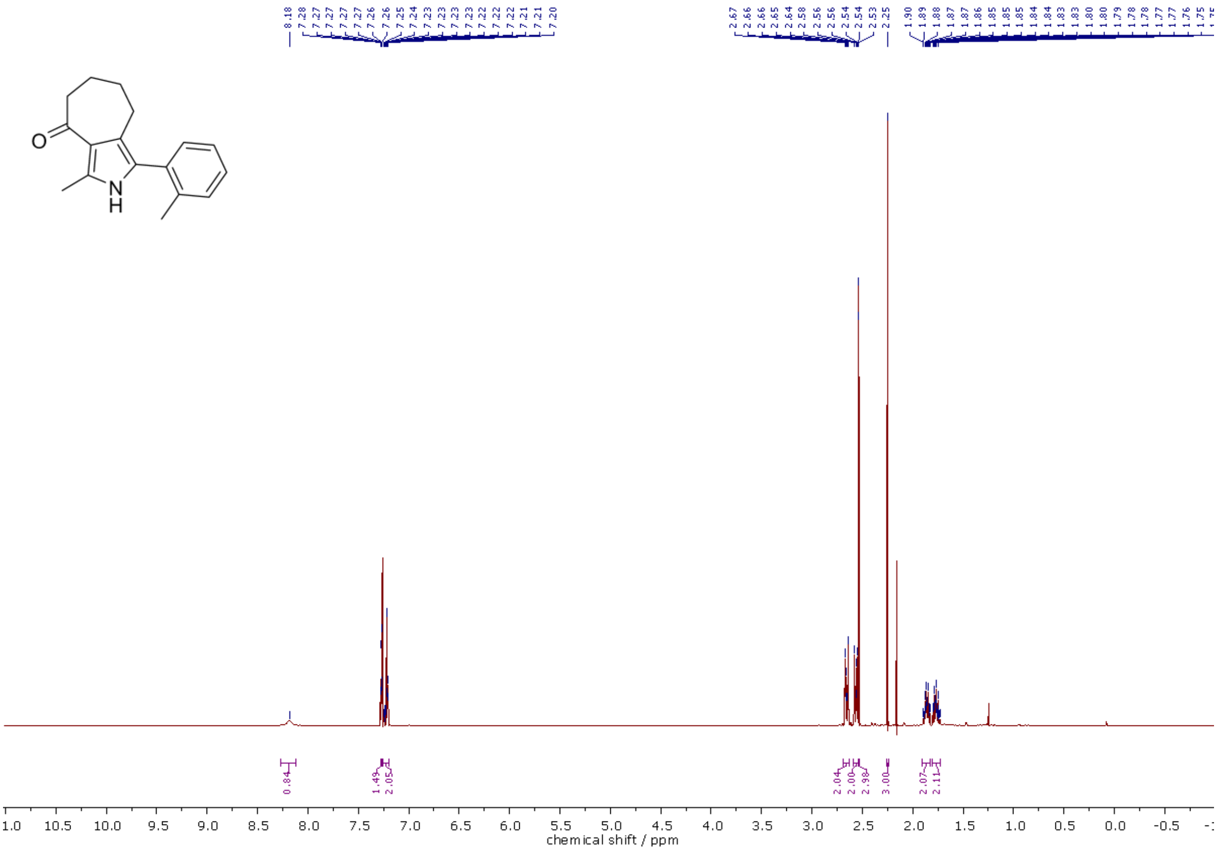

$^{1}H$ NMR (400 MHz, $CDCl_3$) spectra of **305**.

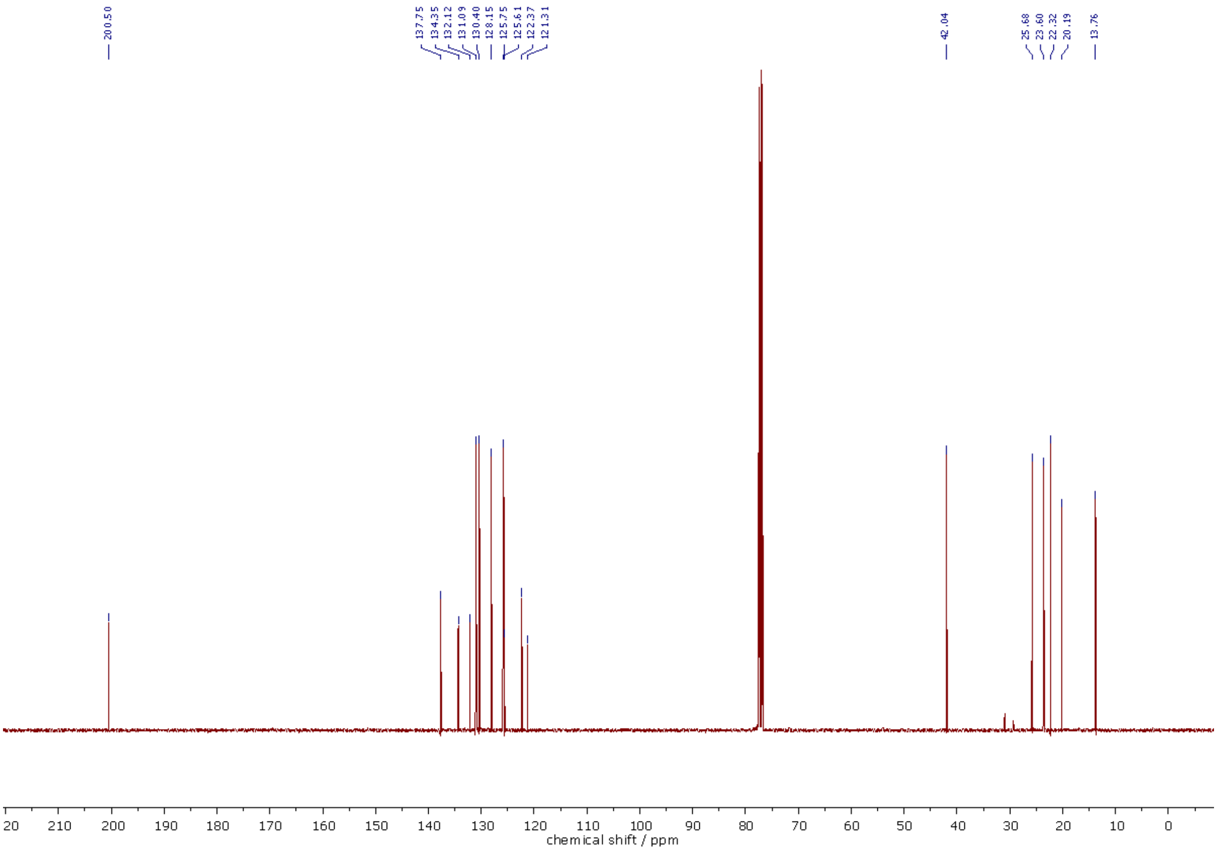

$^{13}C$ NMR (101 MHz, $CDCl_3$) spectra of **305**.

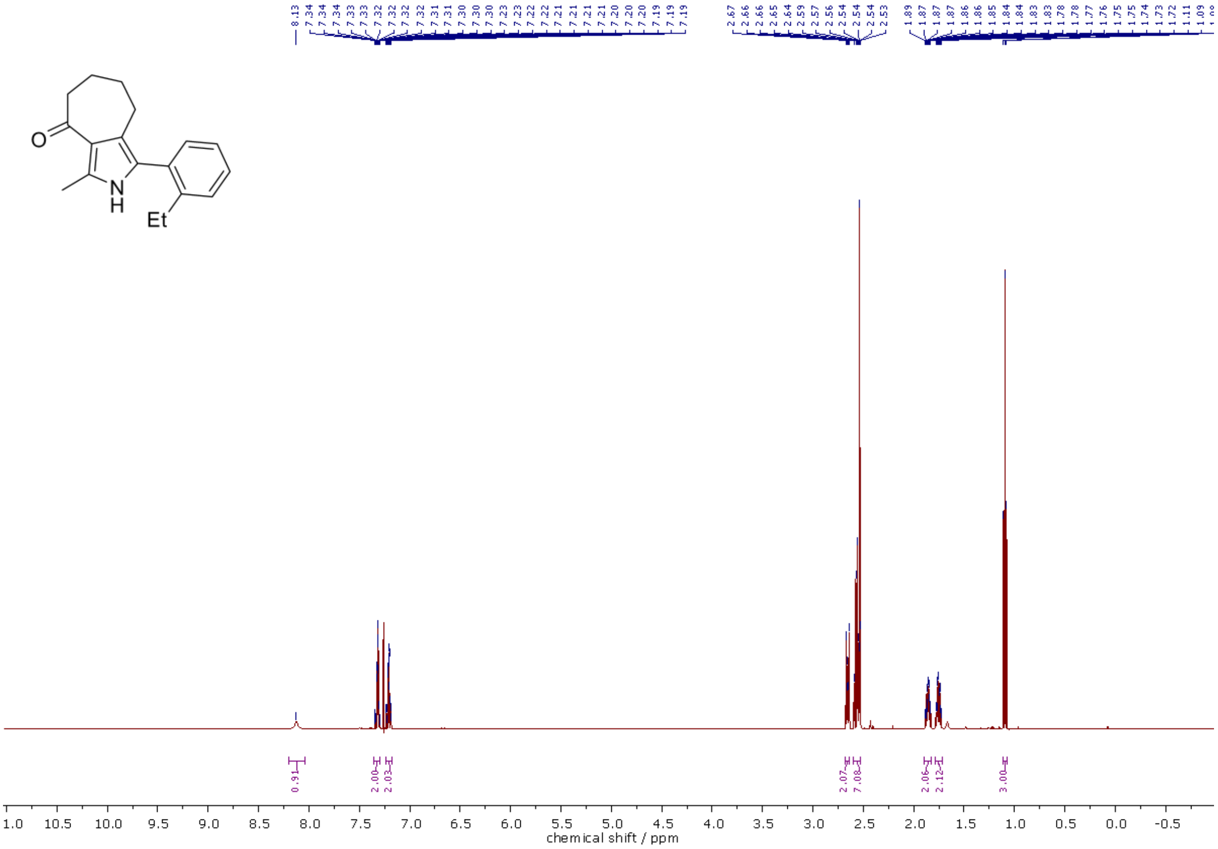

$^{1}H$ NMR (500 MHz, $CDCl_3$) spectra of **306**.

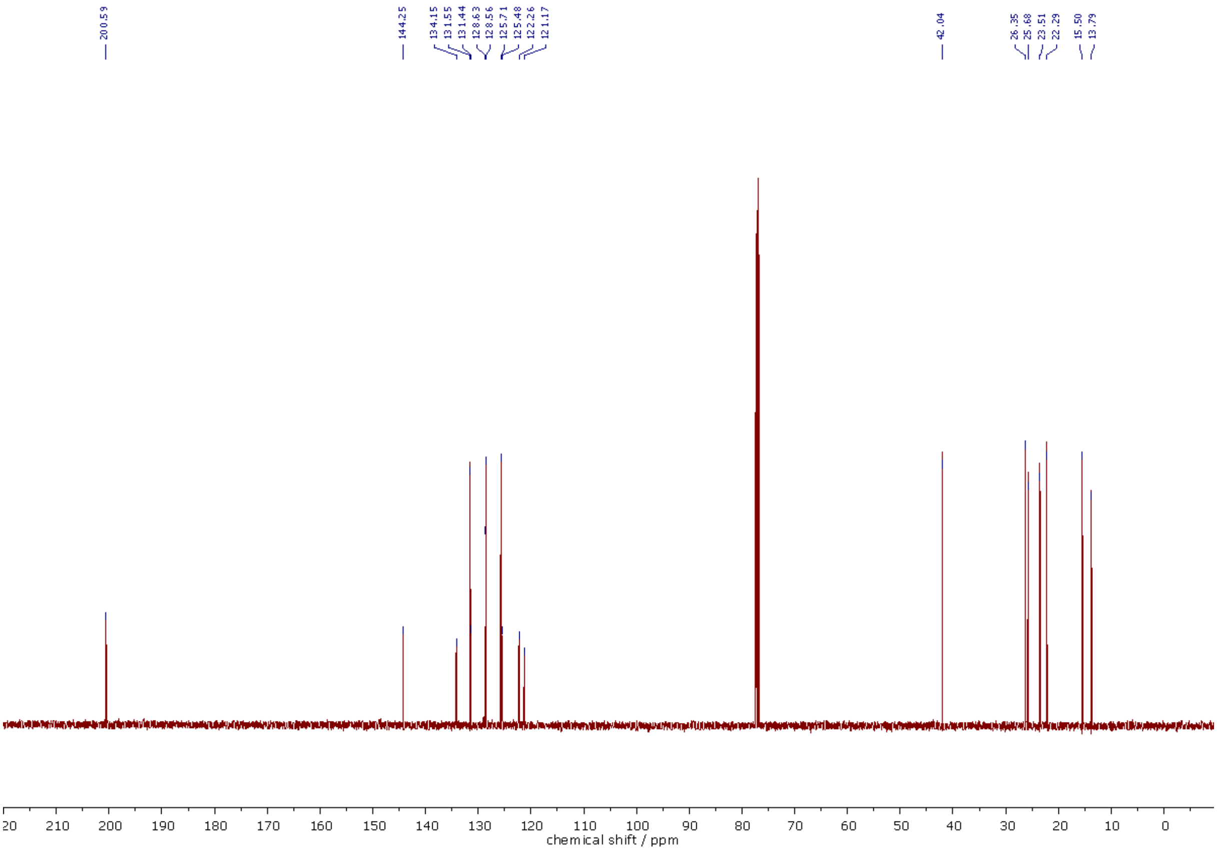

$^{13}C$ NMR (126 MHz, $CDCl_3$) spectra of **306**.

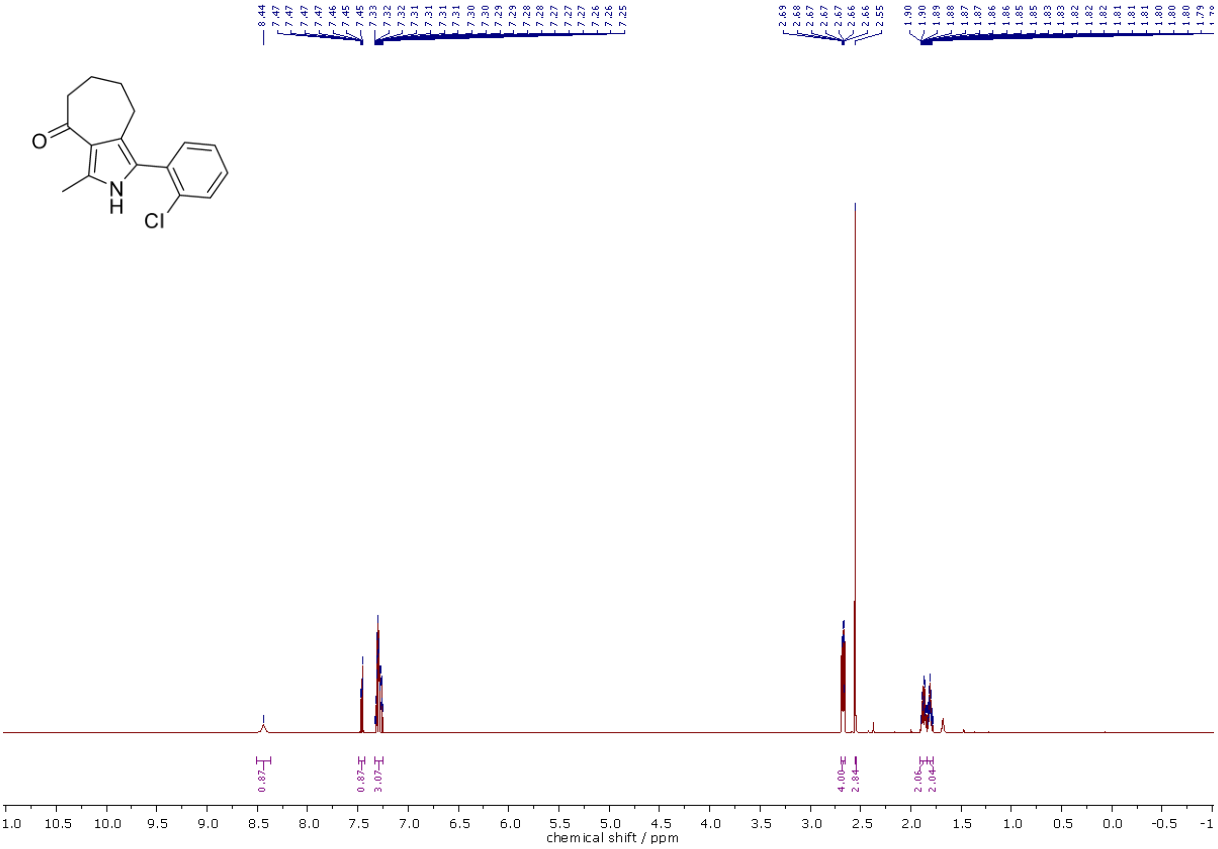

$^{1}H$ NMR (500 MHz, $CDCl_3$) spectra of **307**.

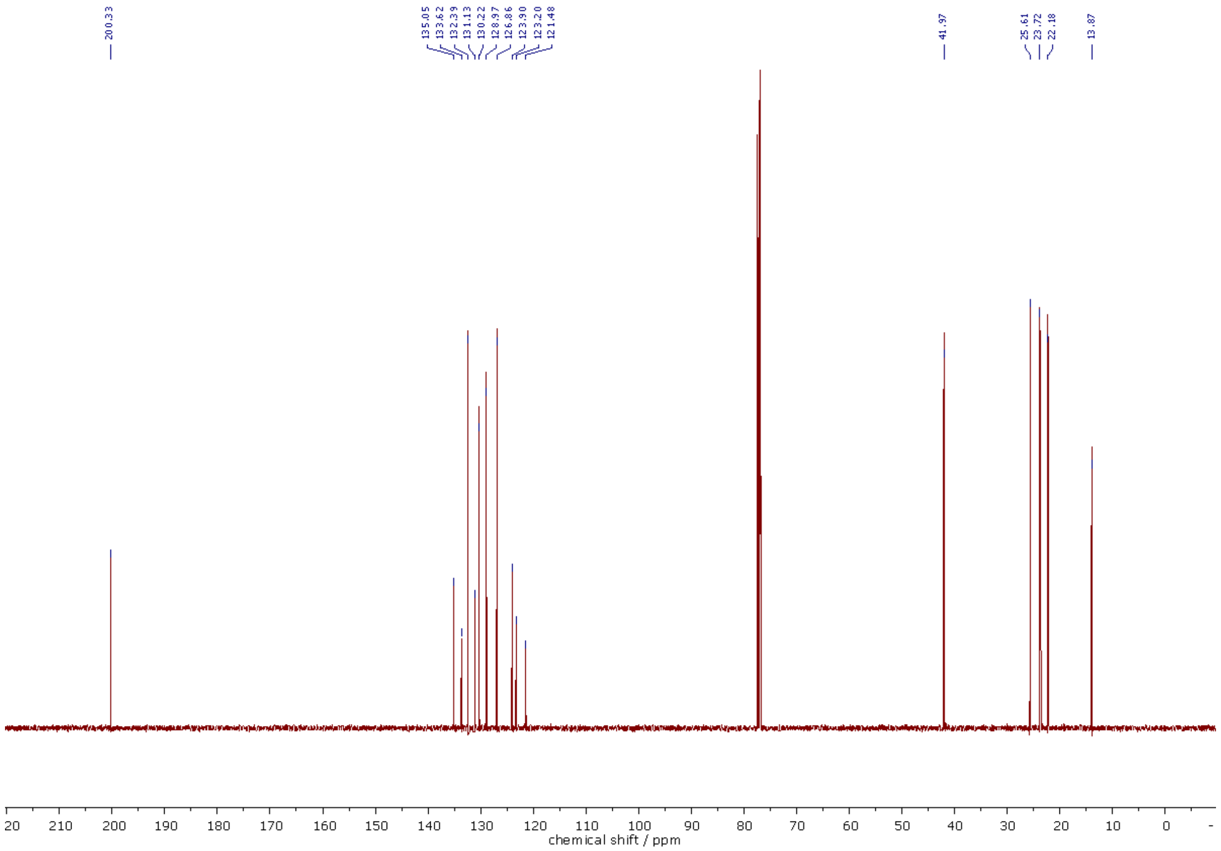

$^{13}C$ NMR (126 MHz, $CDCl_3$) spectra of **307**.

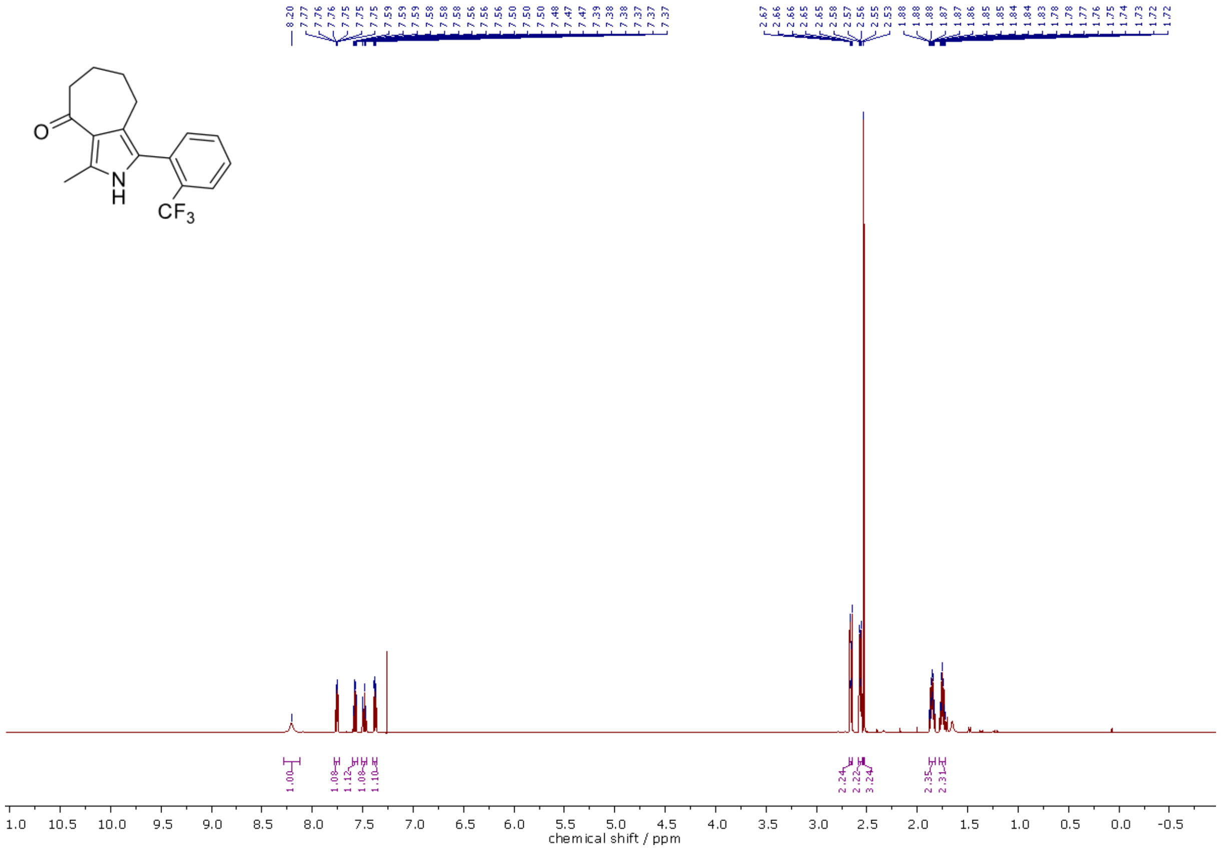

$^{1}H$ NMR (500 MHz, $CDCl_3$) spectra of **308**.

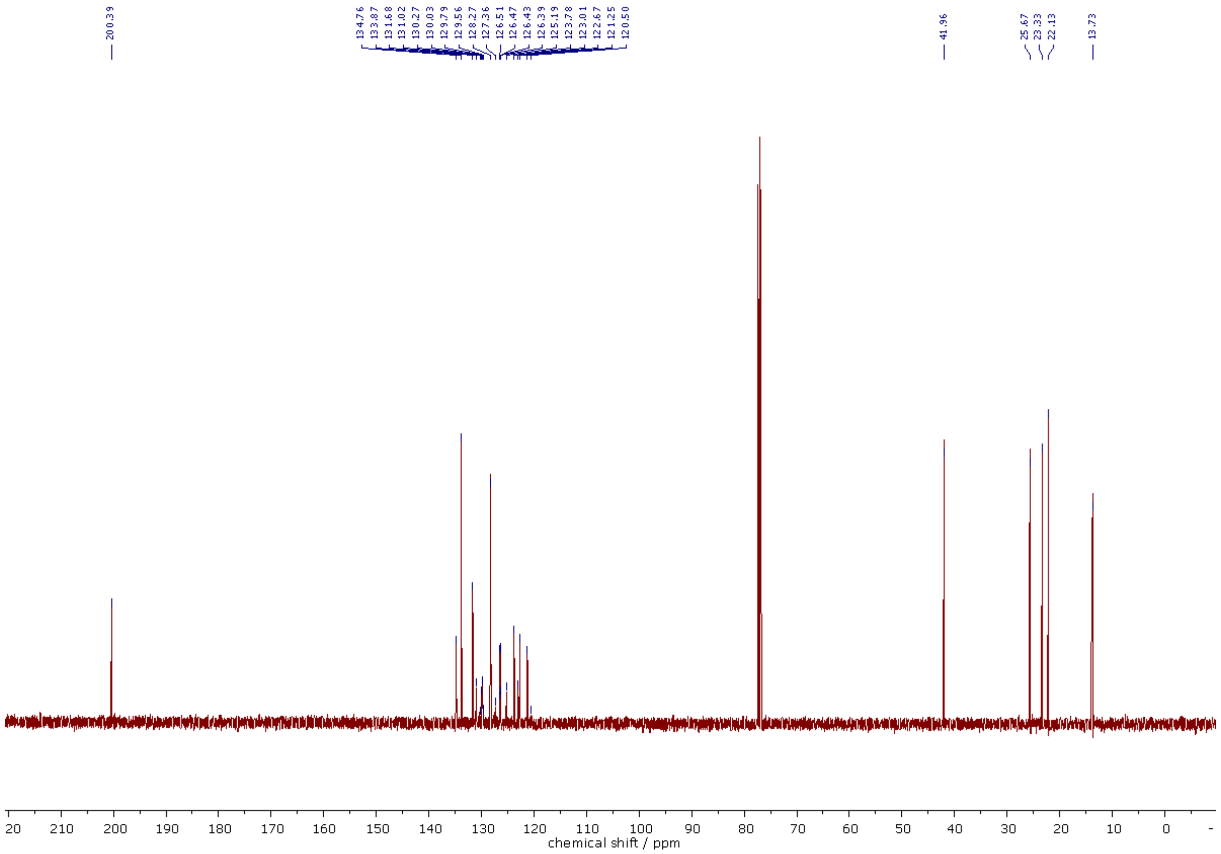

$^{13}C$ NMR (126 MHz, $CDCl_3$) spectra of **308**.

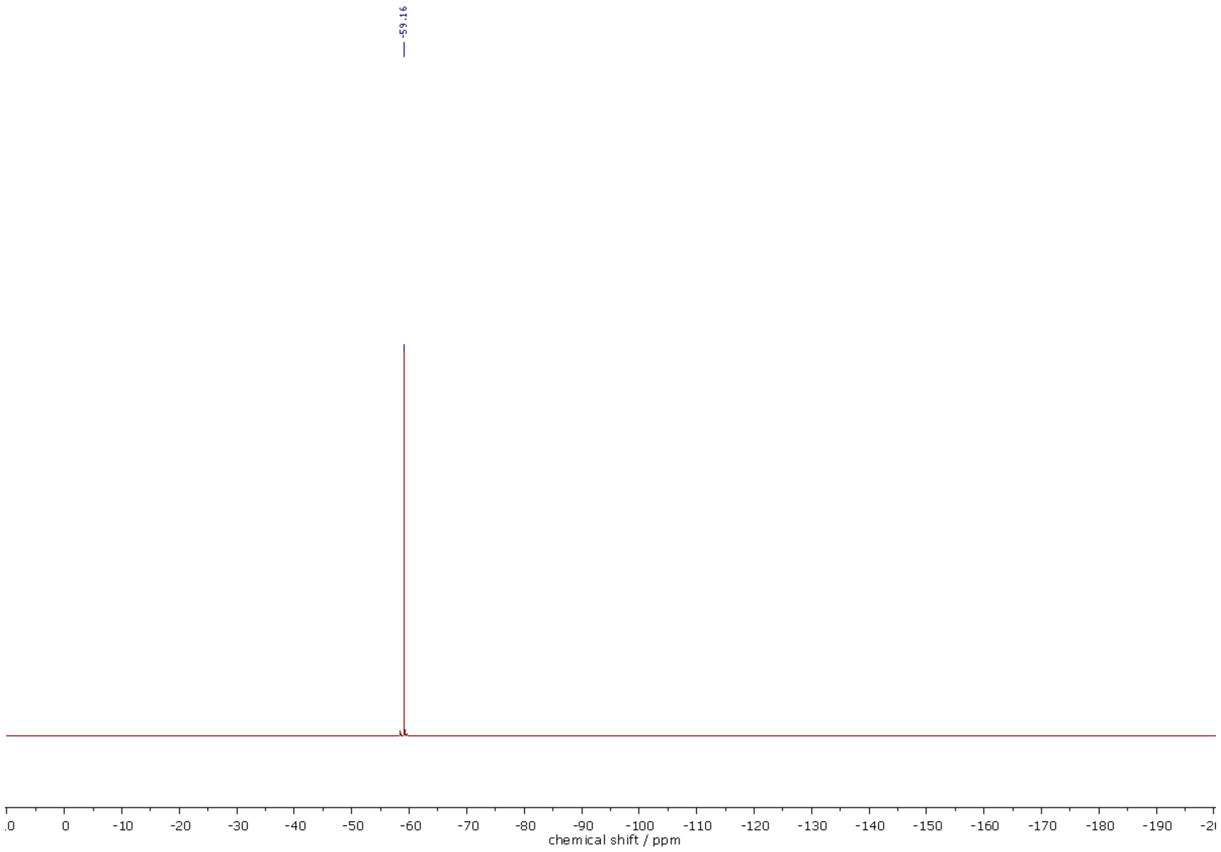

$^{19}F$ NMR (471 MHz, $CDCl_3$) spectra of **308**.

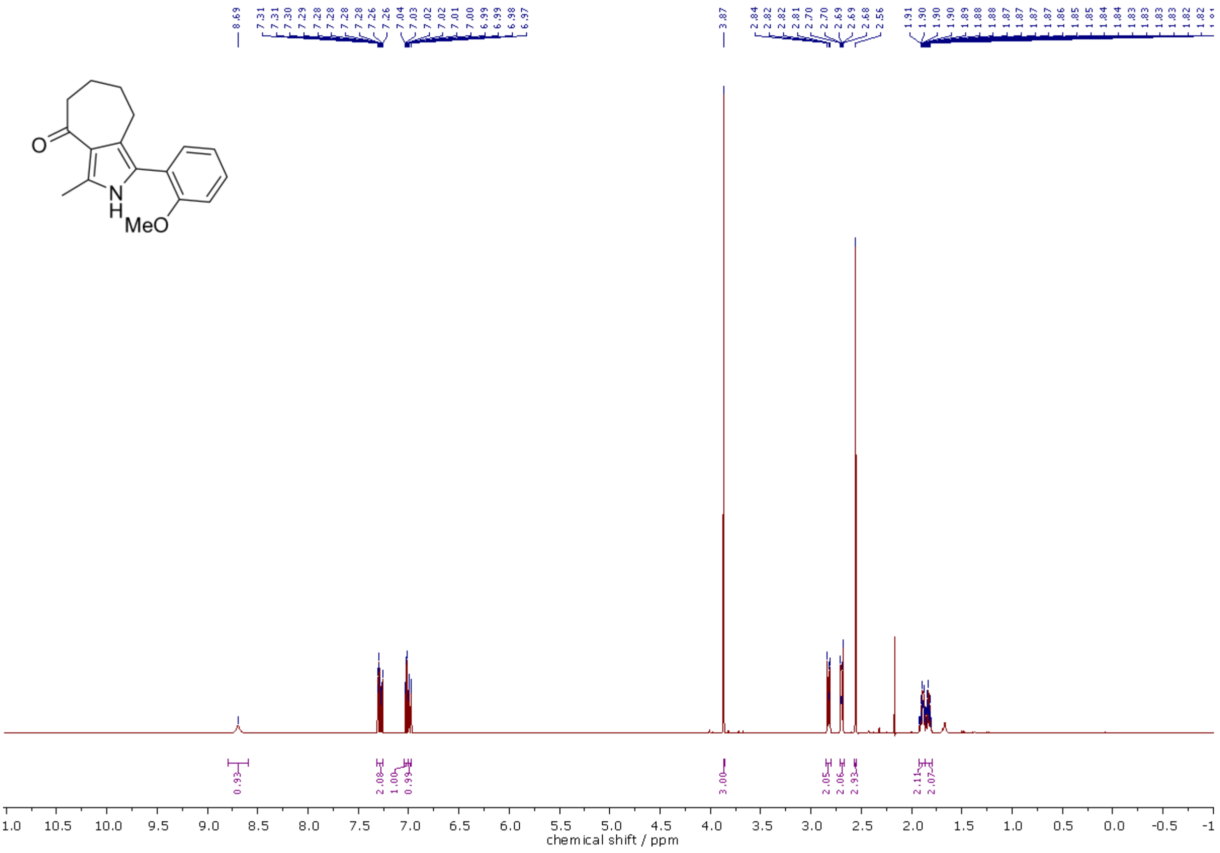

[1]H NMR (500 MHz, $CDCl_3$) spectra of **309**.

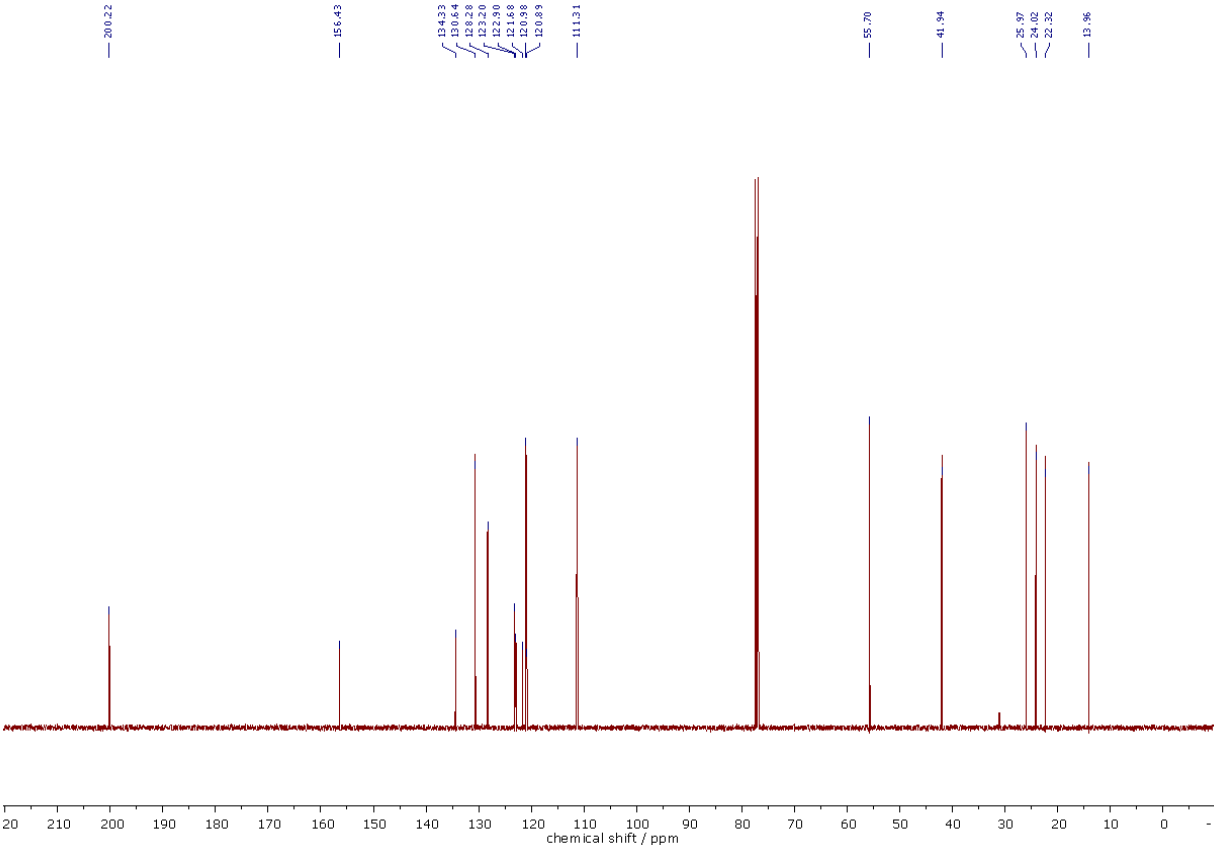

[13]C NMR (126 MHz, $CDCl_3$) spectra of **309**.

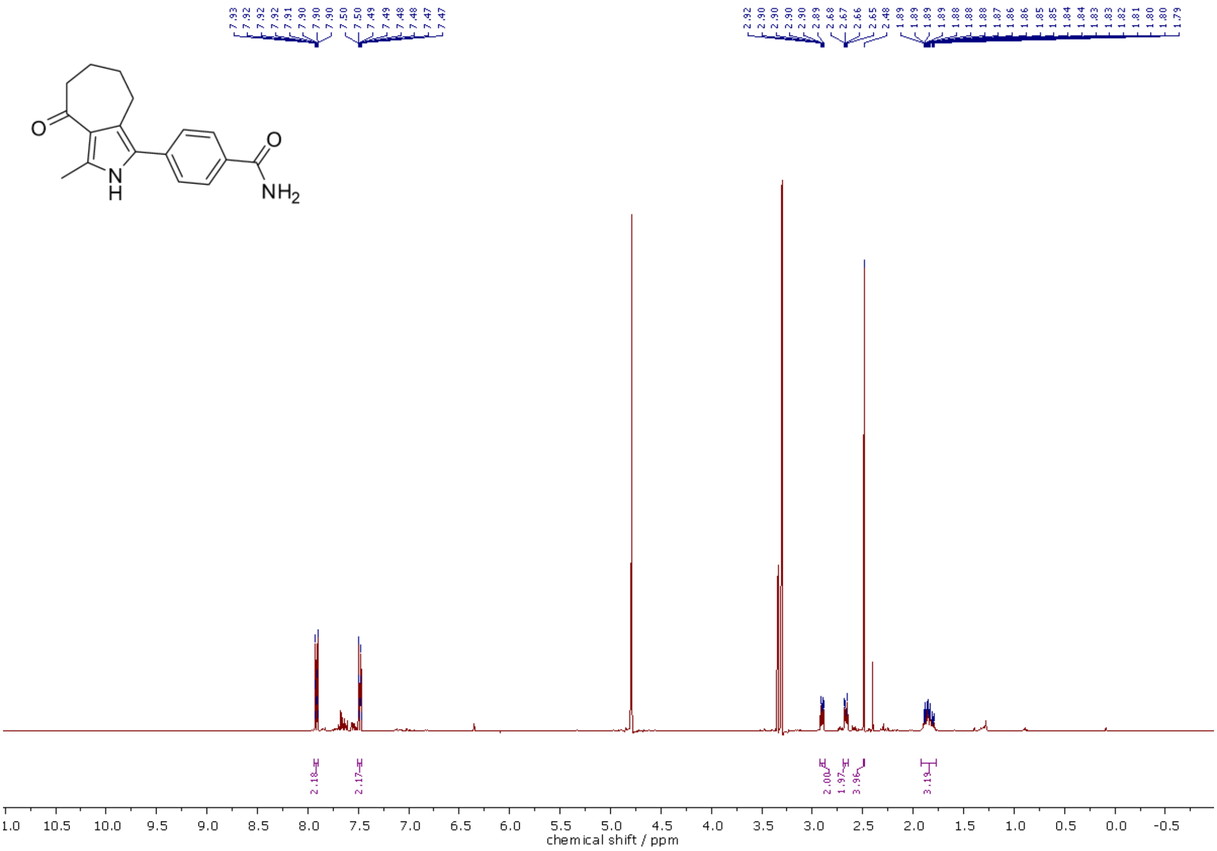

[1]H NMR (400 MHz, MeOD-d4) spectra of **310**.

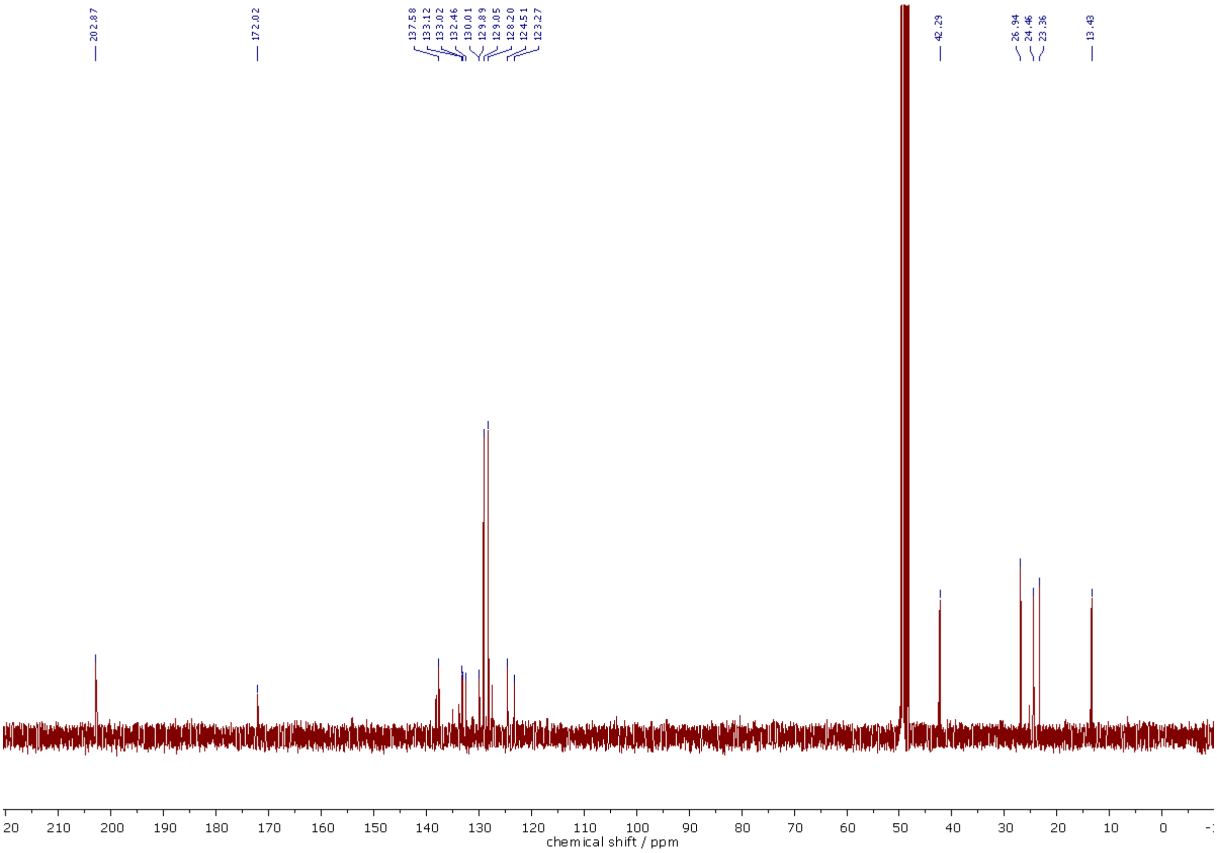

[13]C NMR (101 MHz, MeOD-d4) spectra of **310**.

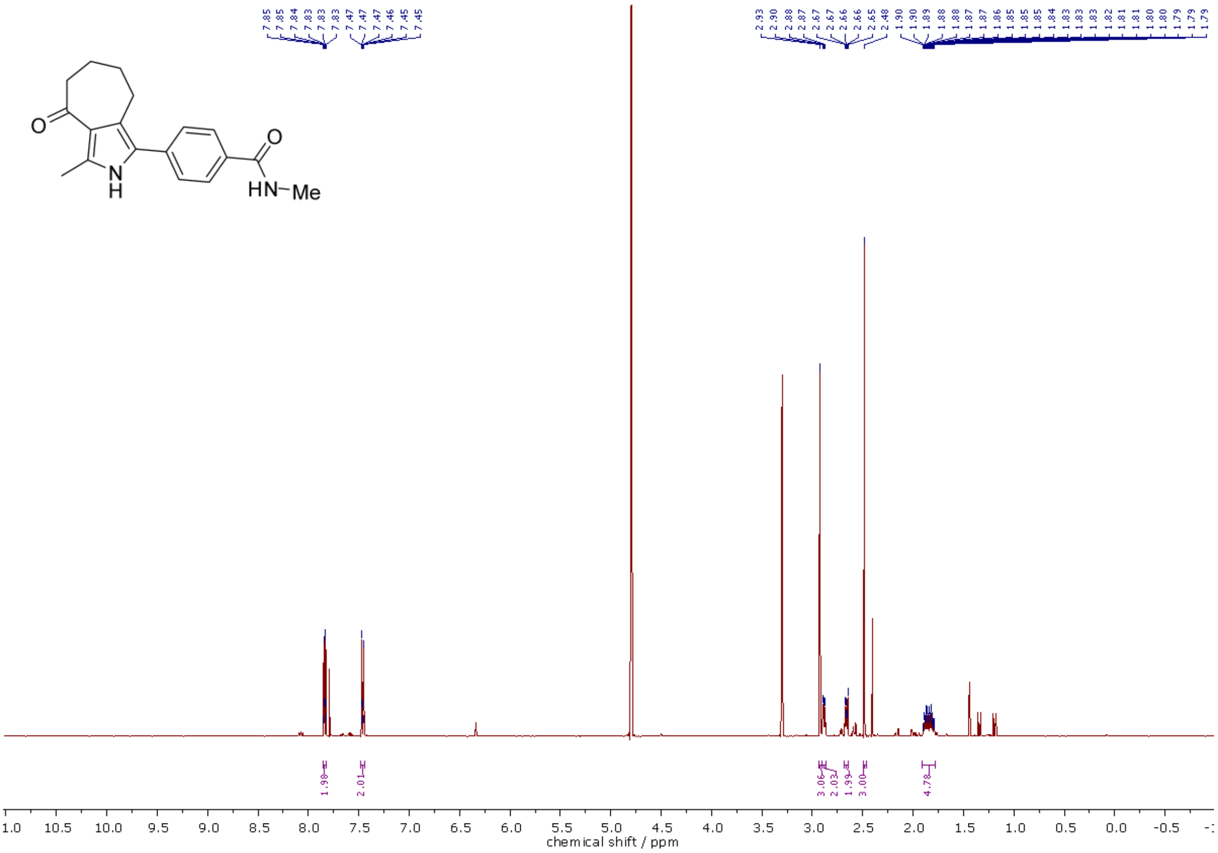

[1]H NMR (500 MHz, $CDCl_3$/MeOD-$d_4$, 5:1) spectra of **311**.

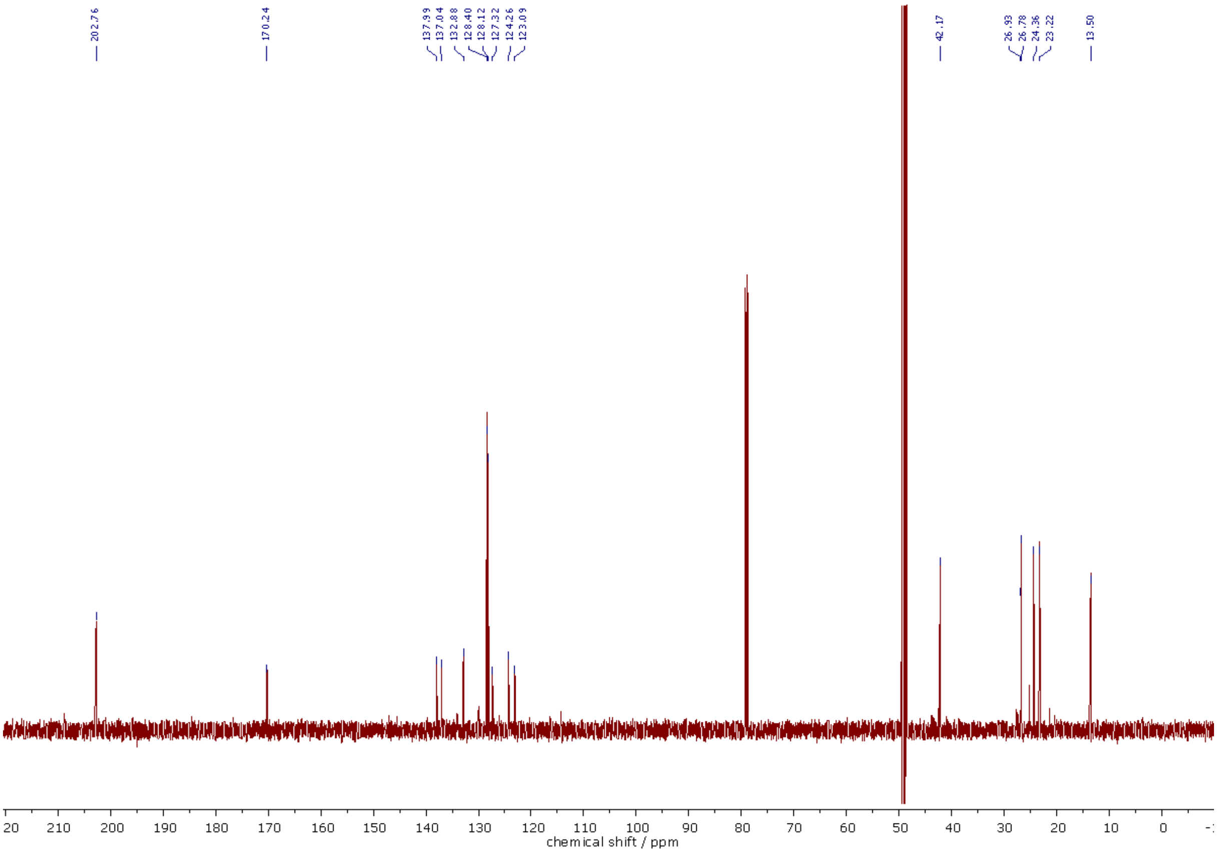

[13]C NMR (126 MHz, $CDCl_3$/MeOD-$d_4$, 5:1) spectra of **311**.

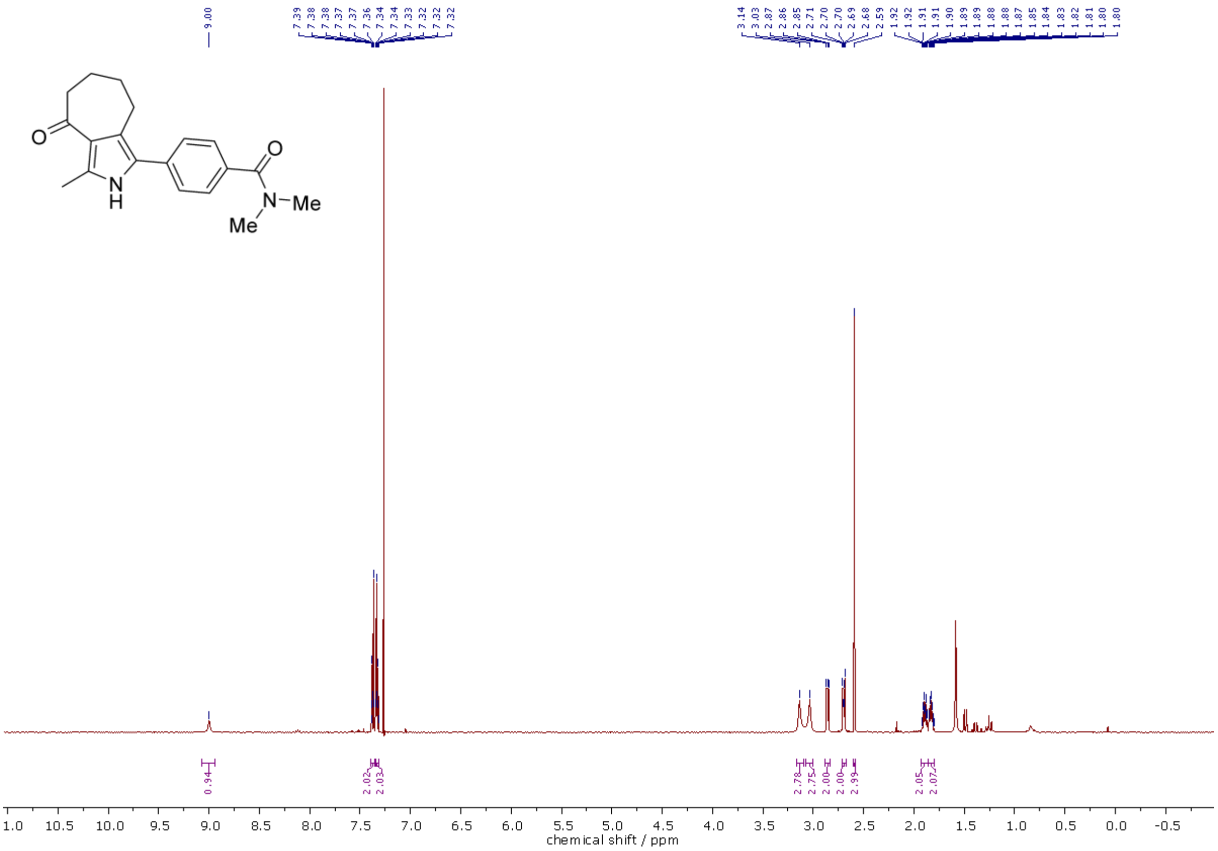

$^{1}$H NMR (500 MHz, $CDCl_3$) spectra of **312**.

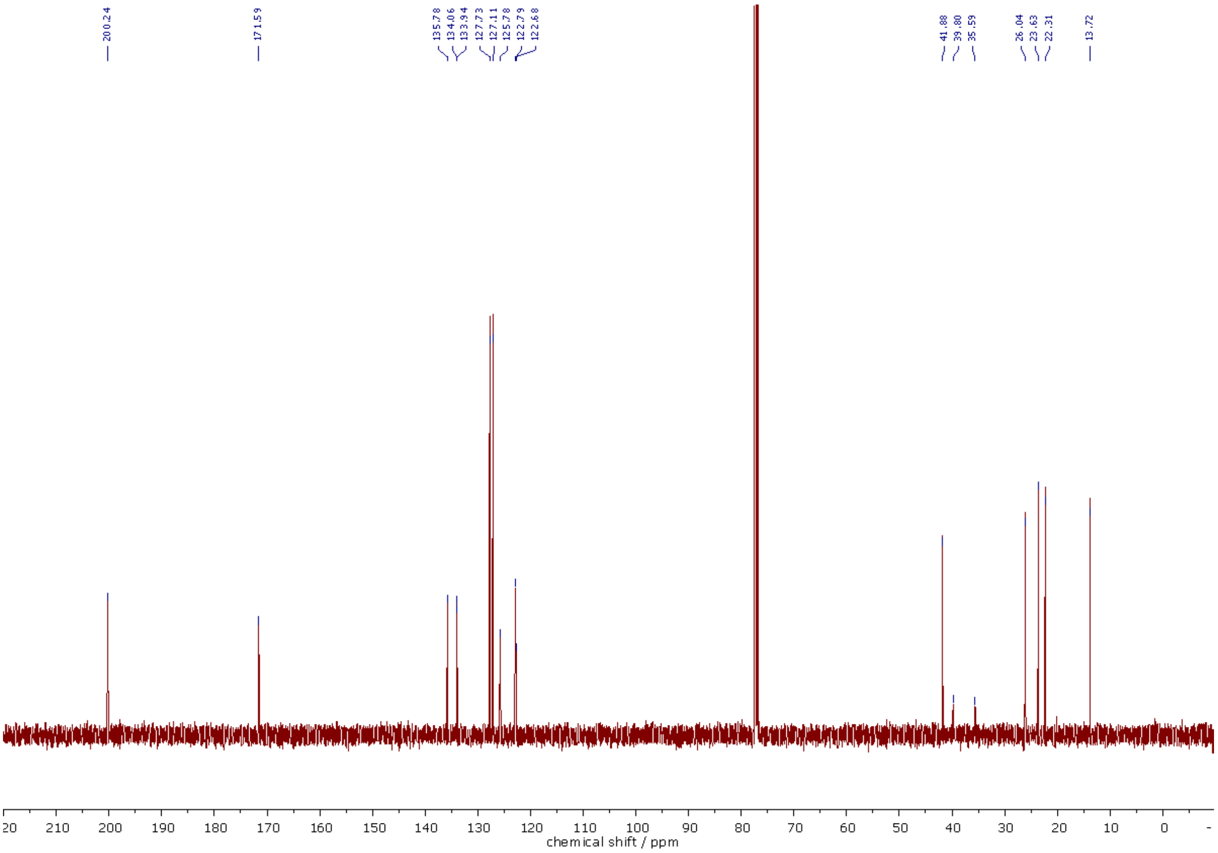

$^{13}$C NMR (126 MHz, $CDCl_3$) spectra of **312**.

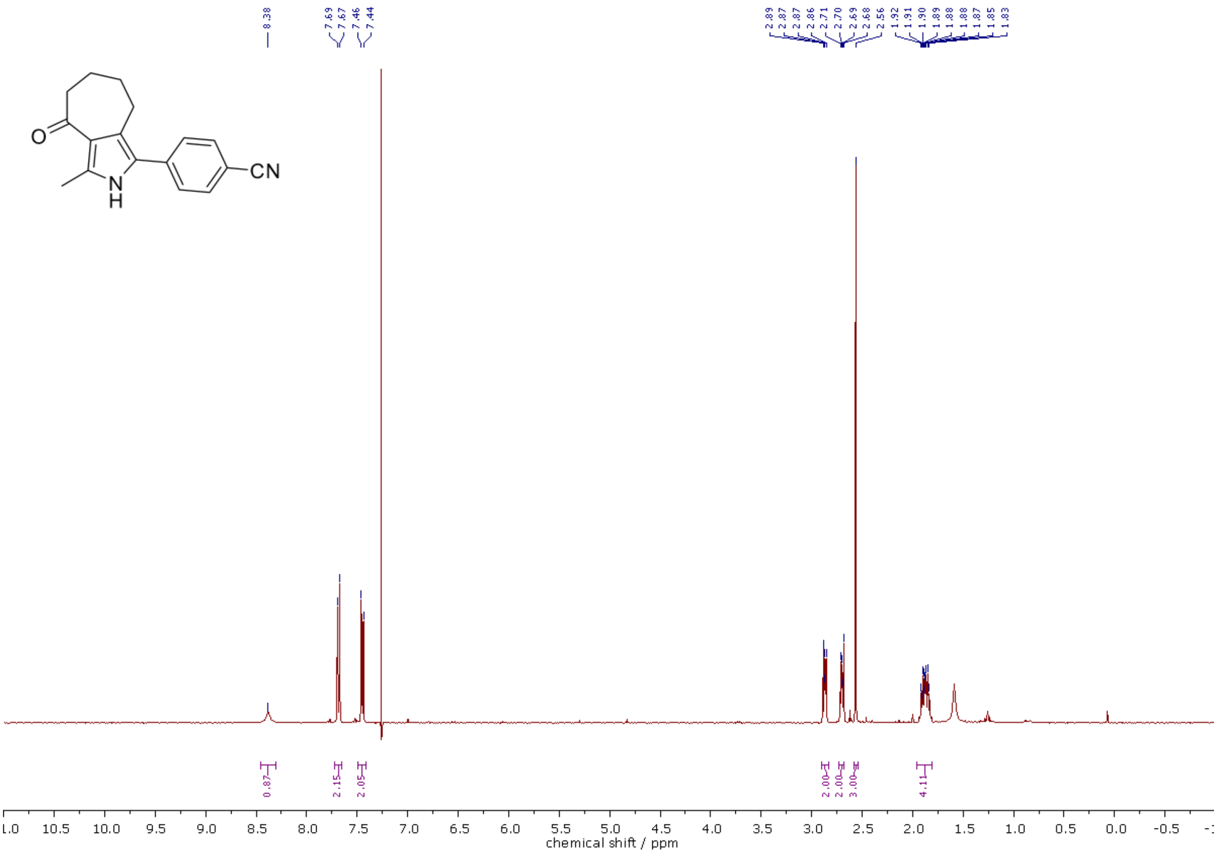

[1]H NMR (400 MHz, $CDCl_3$) spectra of **313**.

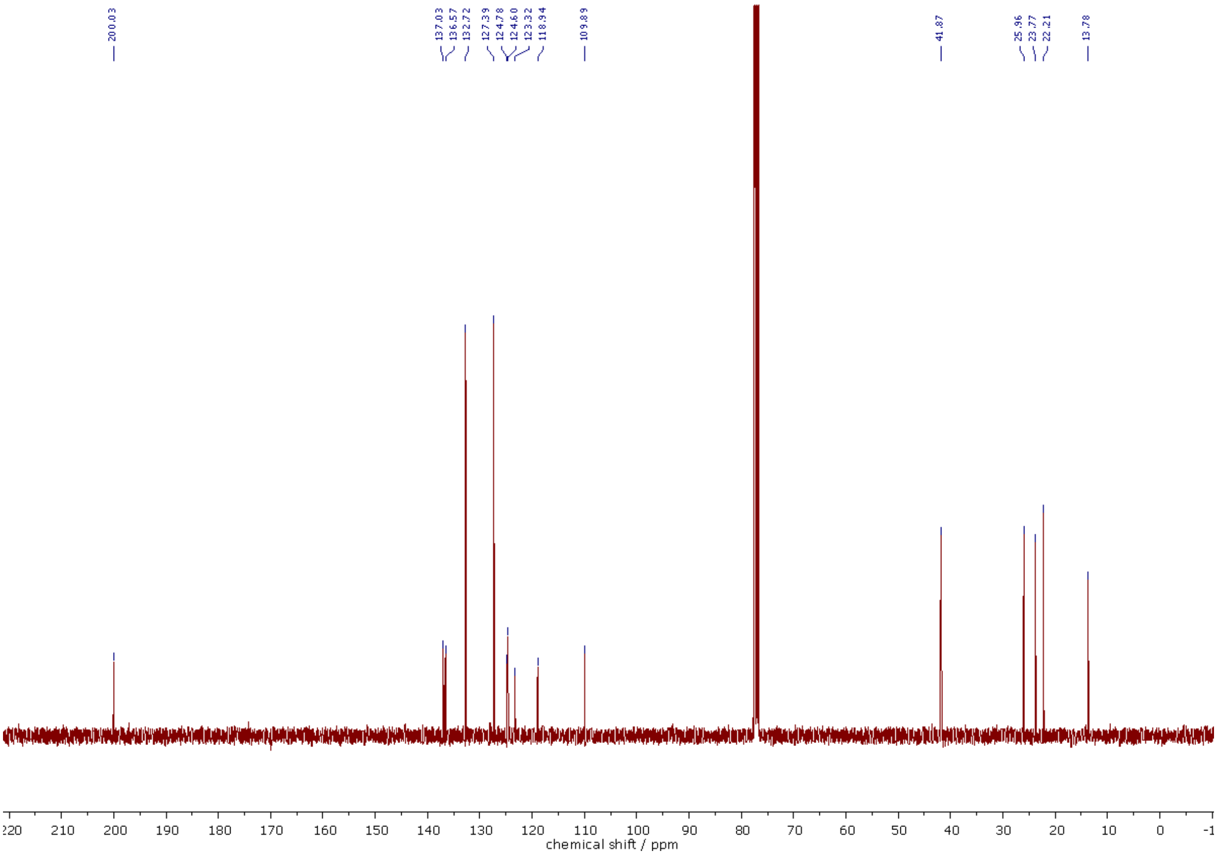

[13]C NMR (101 MHz, $CDCl_3$) spectra of **313**.

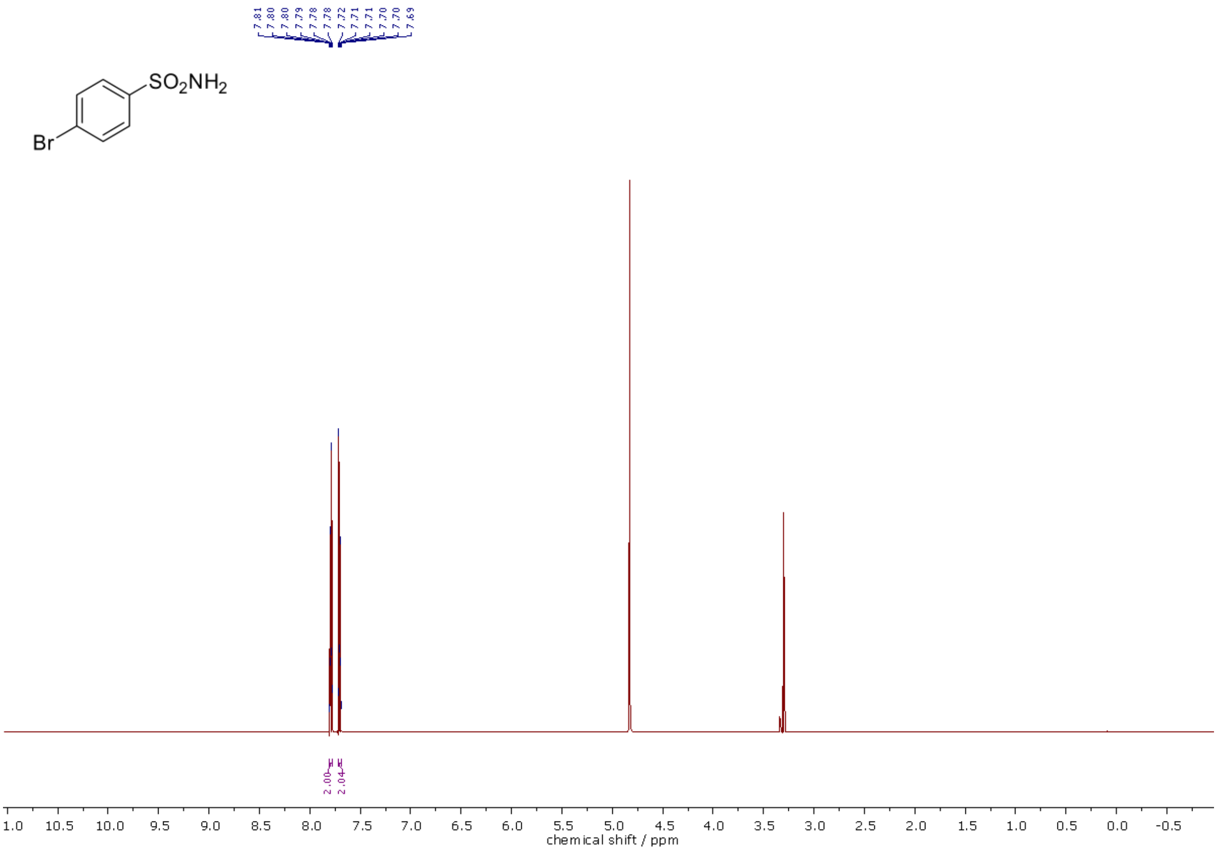

$^{1}H$ NMR (500 MHz, MeOD-d$_4$) spectra of **317**.

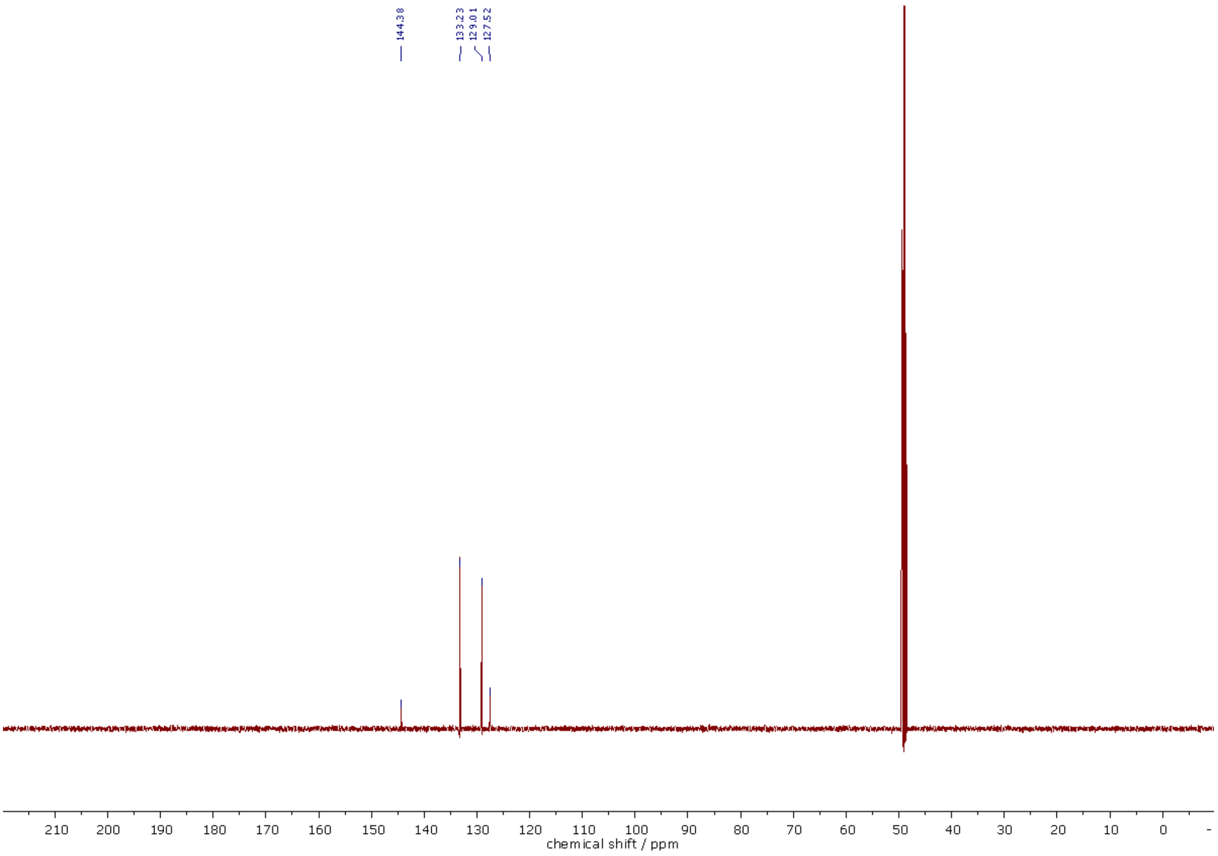

$^{13}C$ NMR (101 MHz, MeOD-d$_4$) spectra of **317**.

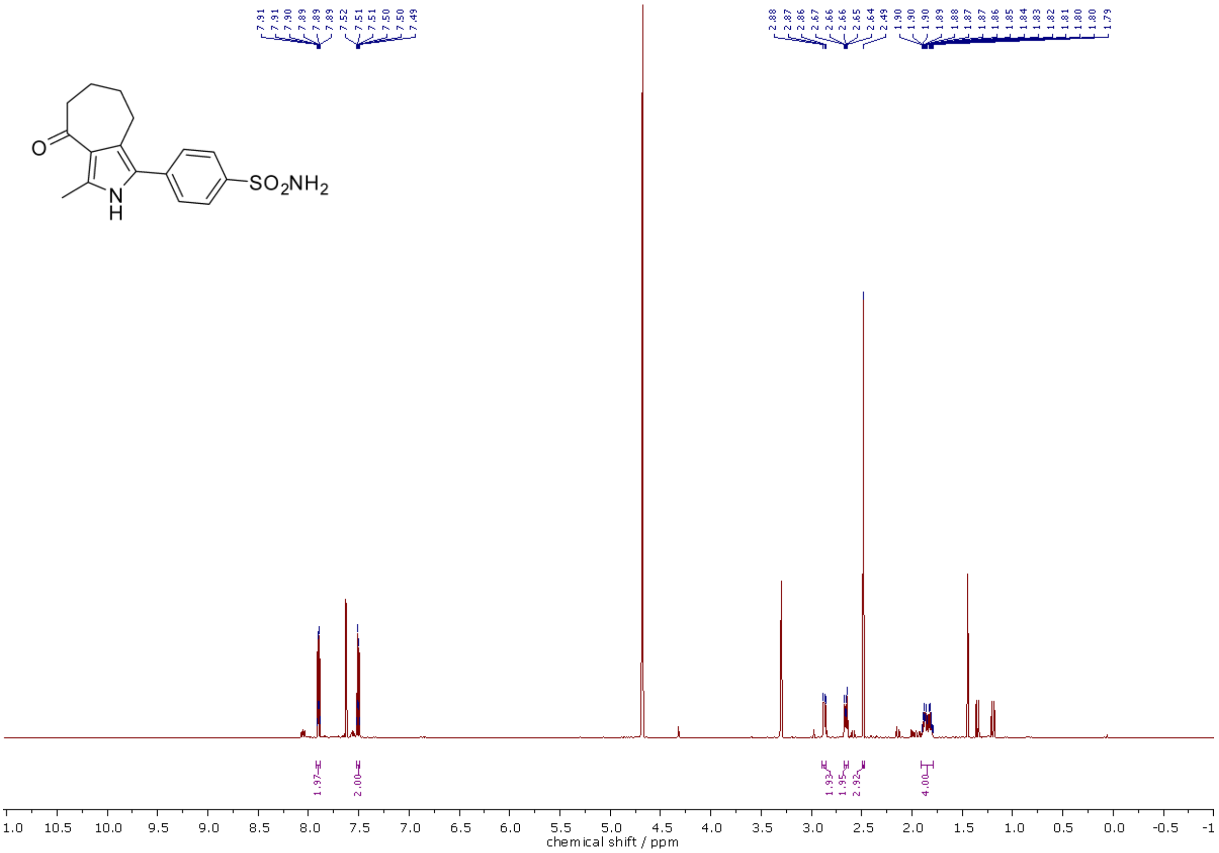

[1]H NMR (500 MHz, MeOD-d4/CDCl3, 3:2) spectra of **314**.

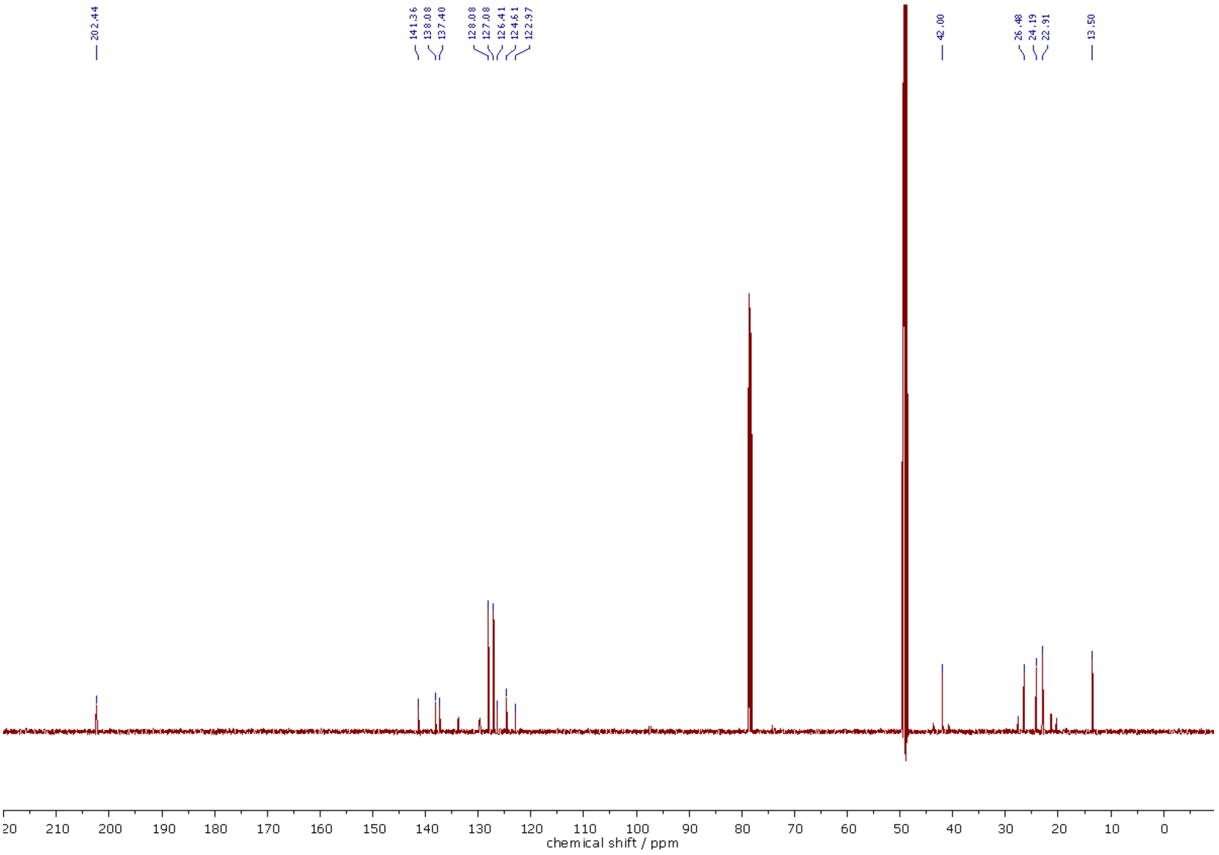

[13]C NMR (126 MHz, MeOD-d4/CDCl3, 3:2) spectra of **314**.

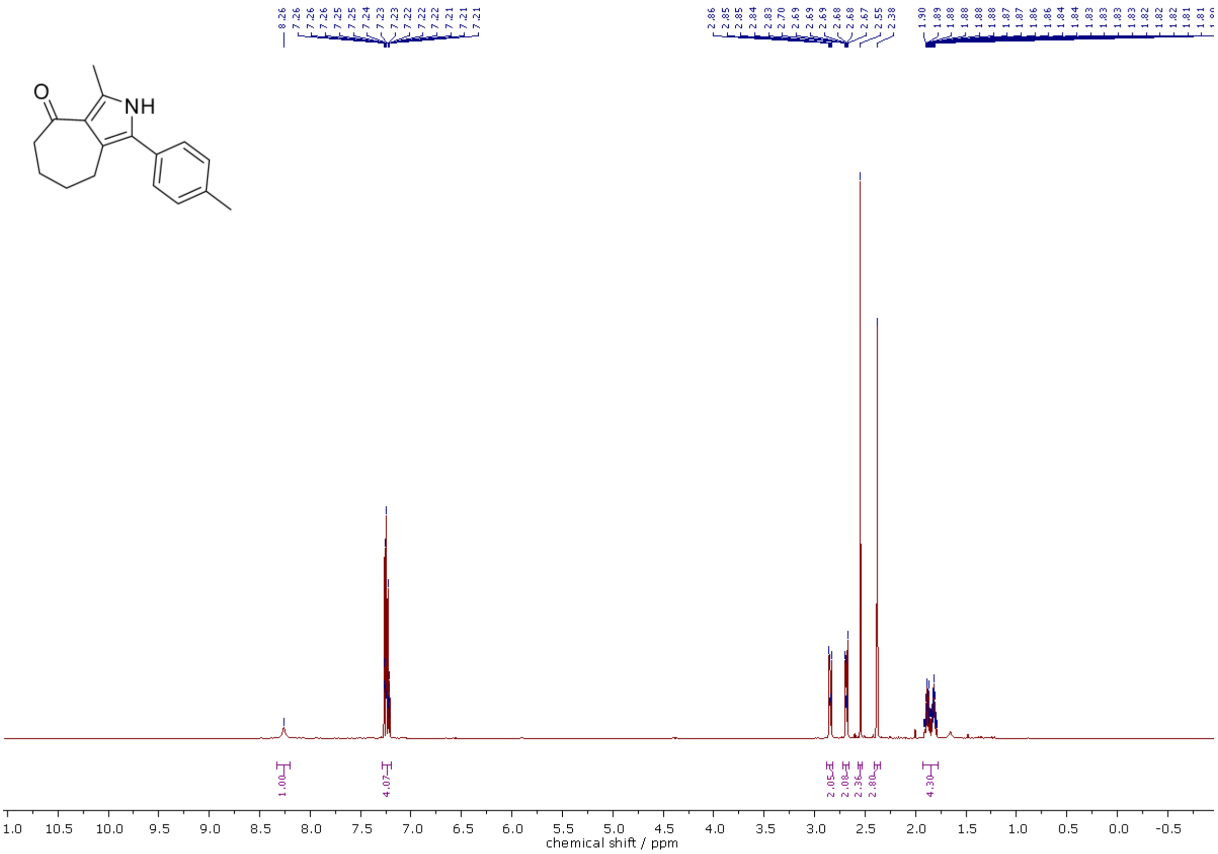

$^{1}$H NMR (500 MHz, $CDCl_3$) spectra of **315**.

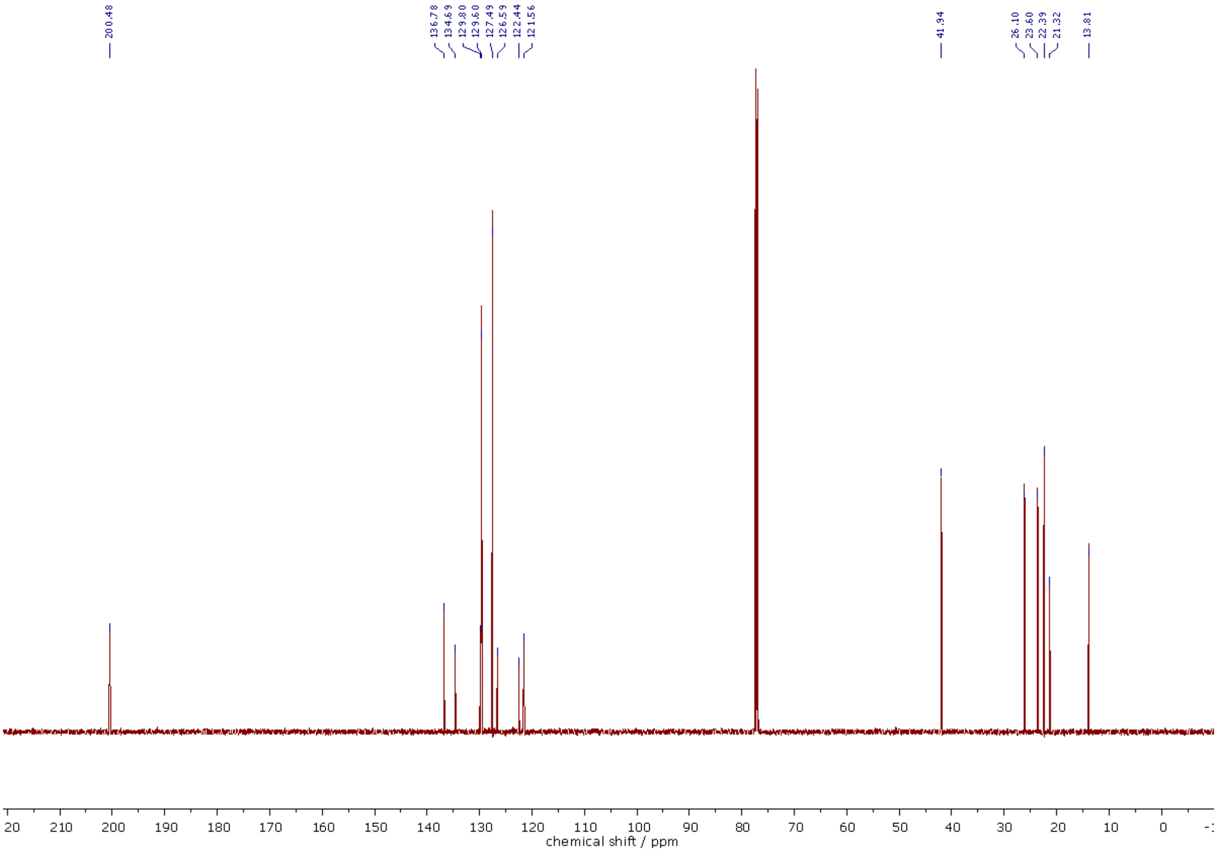

$^{13}$C NMR (126 MHz, $CDCl_3$) spectra of **315**.

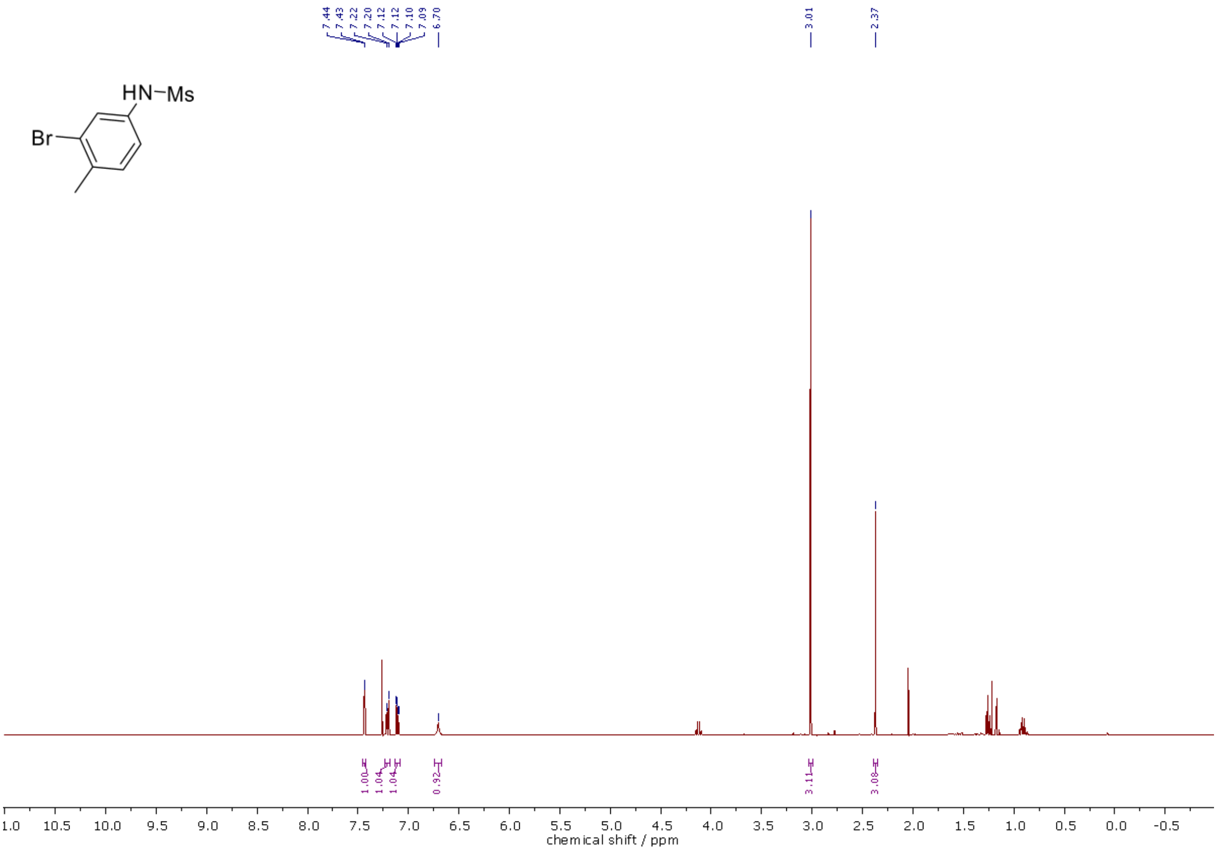

$^1$H NMR (400 MHz, $CDCl_3$) spectra of **339**.

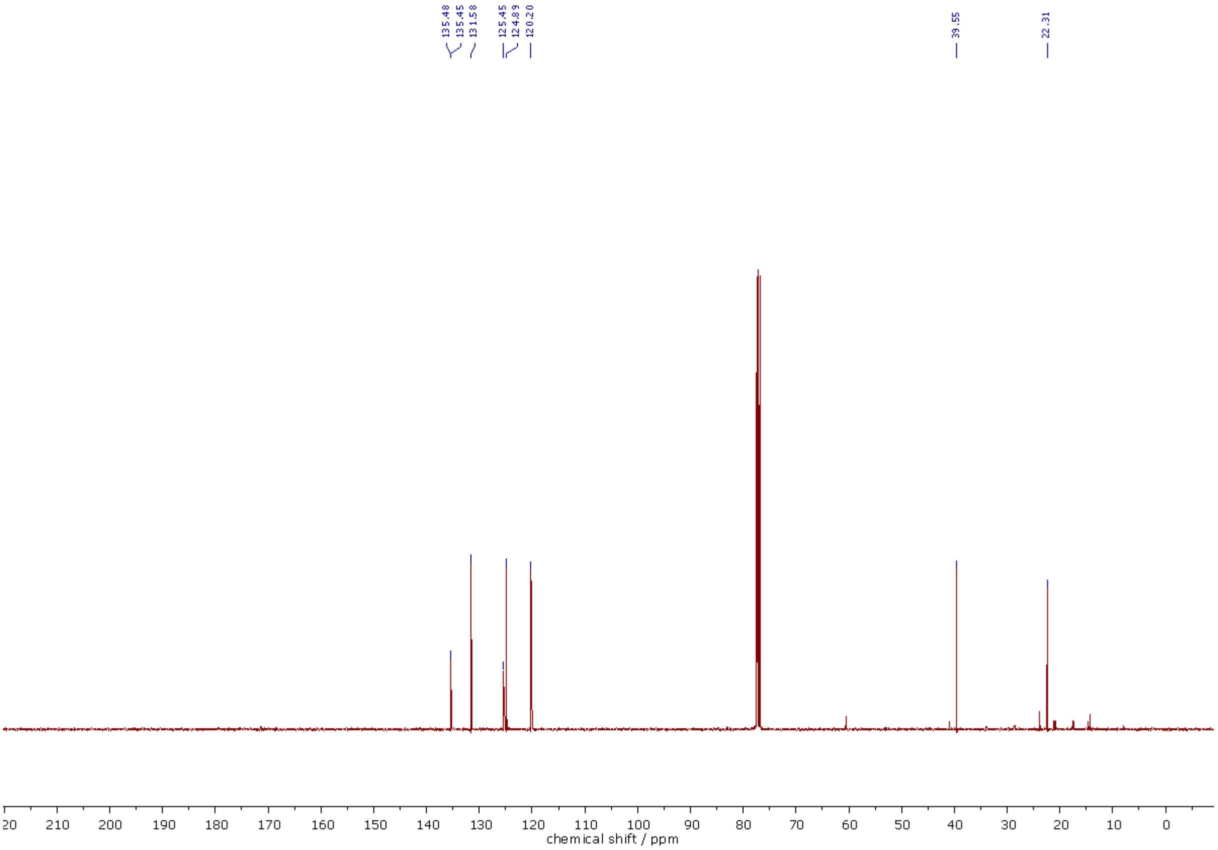

$^{13}$C NMR (126 MHz, $CDCl_3$) spectra of **339**.

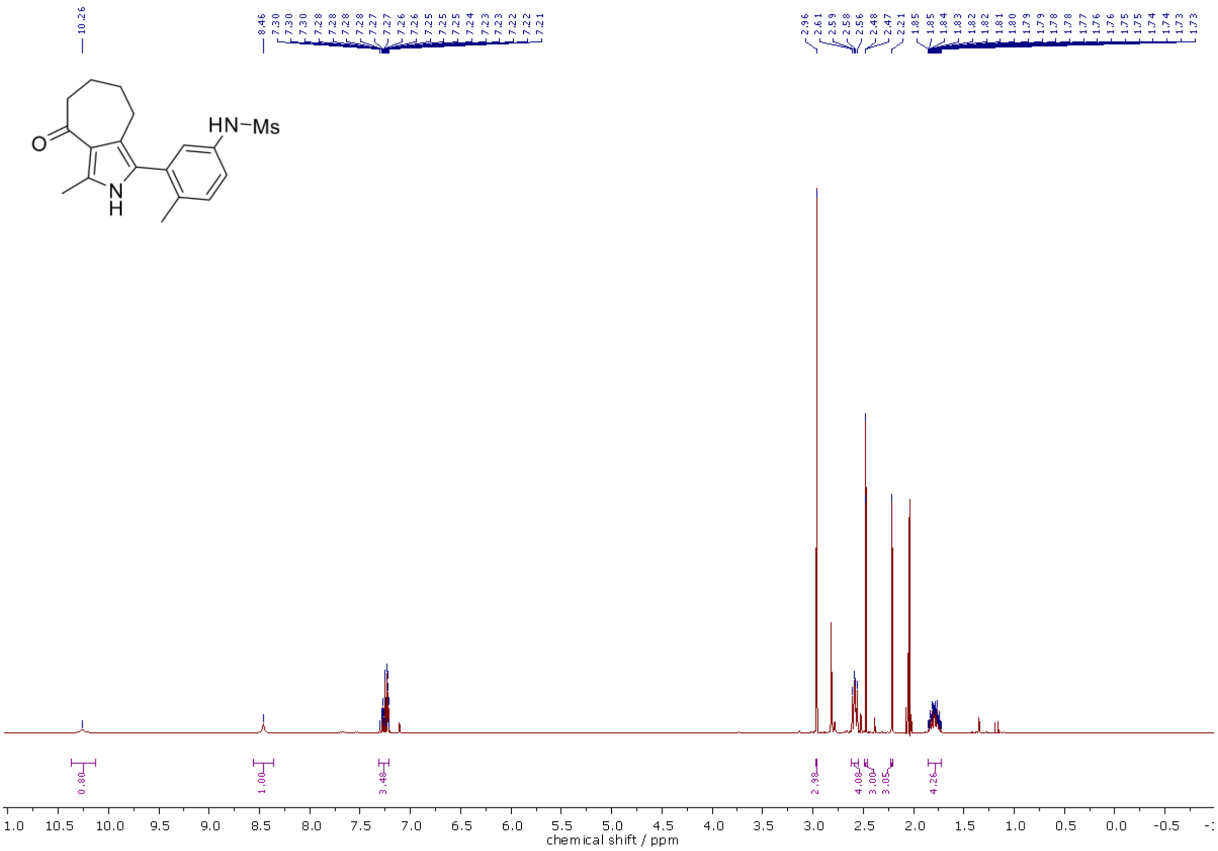

$^{1}$H NMR (400 MHz, Acetone-d$_6$) spectra of **329**.

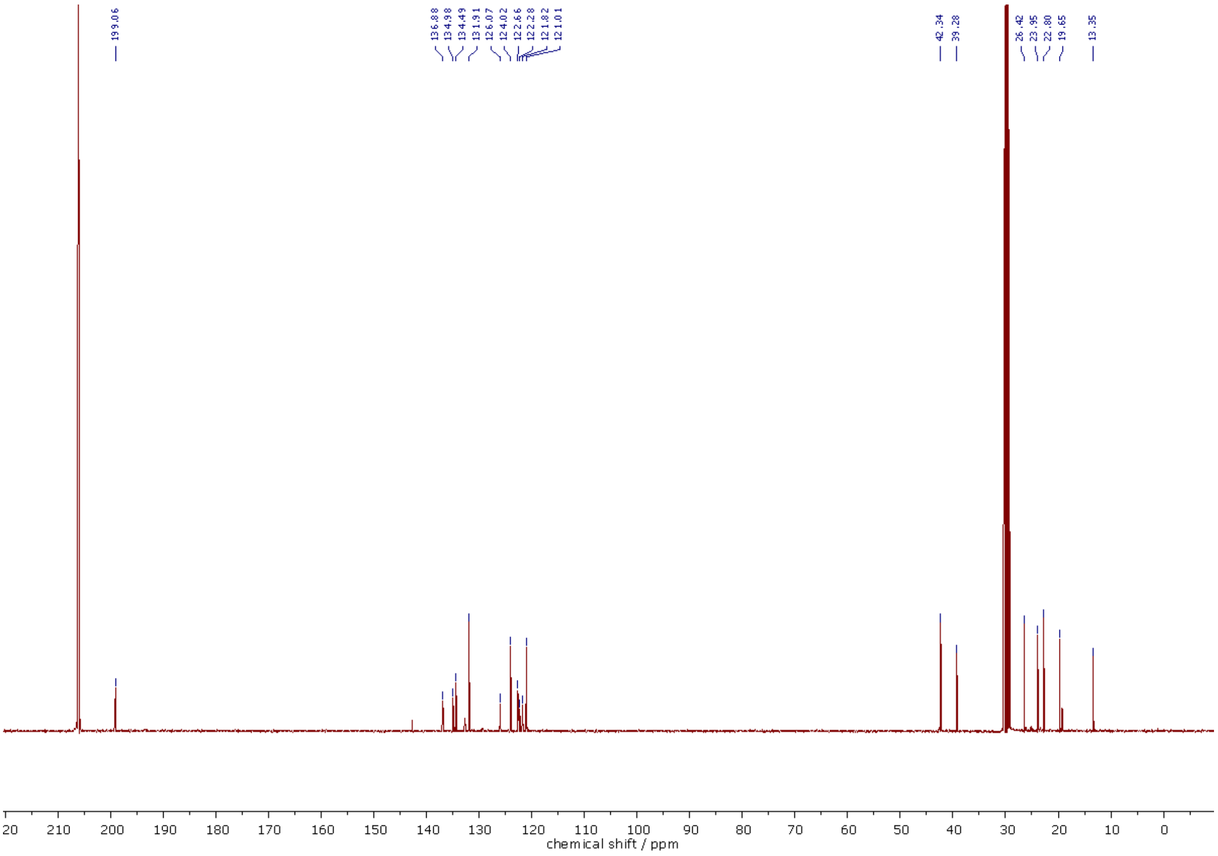

$^{13}$C NMR (101 MHz, Acetone-d$_6$) spectra of **329**.

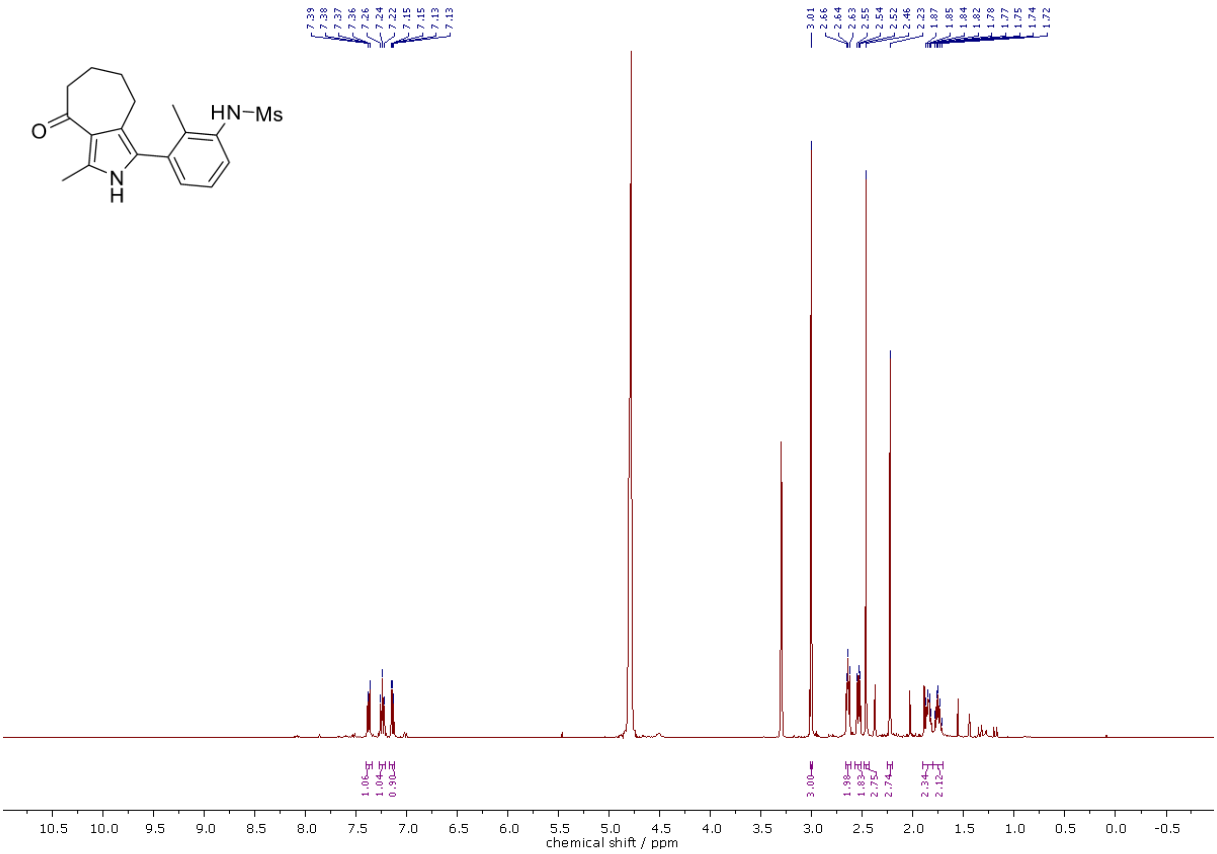

[1]H NMR (400 MHz, MeOD-d4) spectra of **332**.

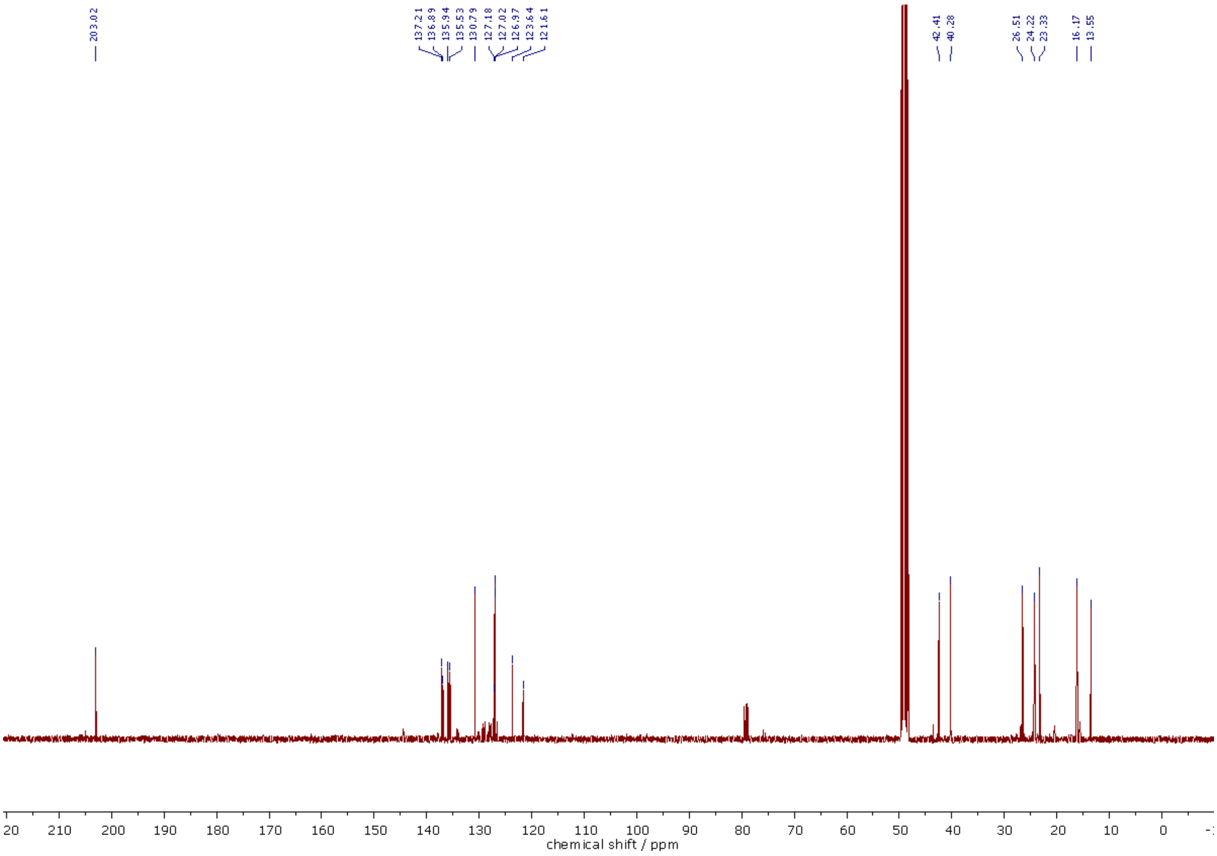

[13]C NMR (126 MHz, MeOD-d4) spectra of **332**.

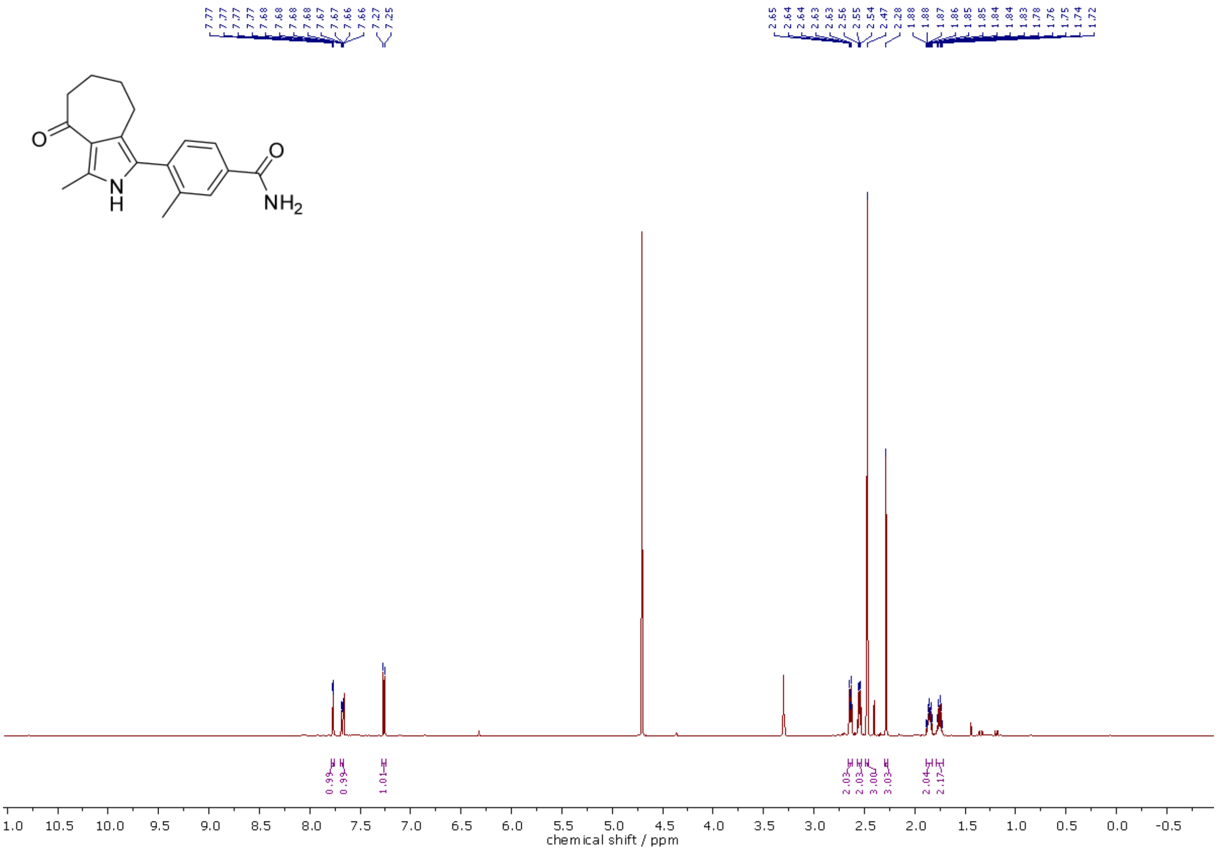

1H NMR (500 MHz, MeOD-$d_4$/$CDCl_3$, 2:1) spectra of **335**.

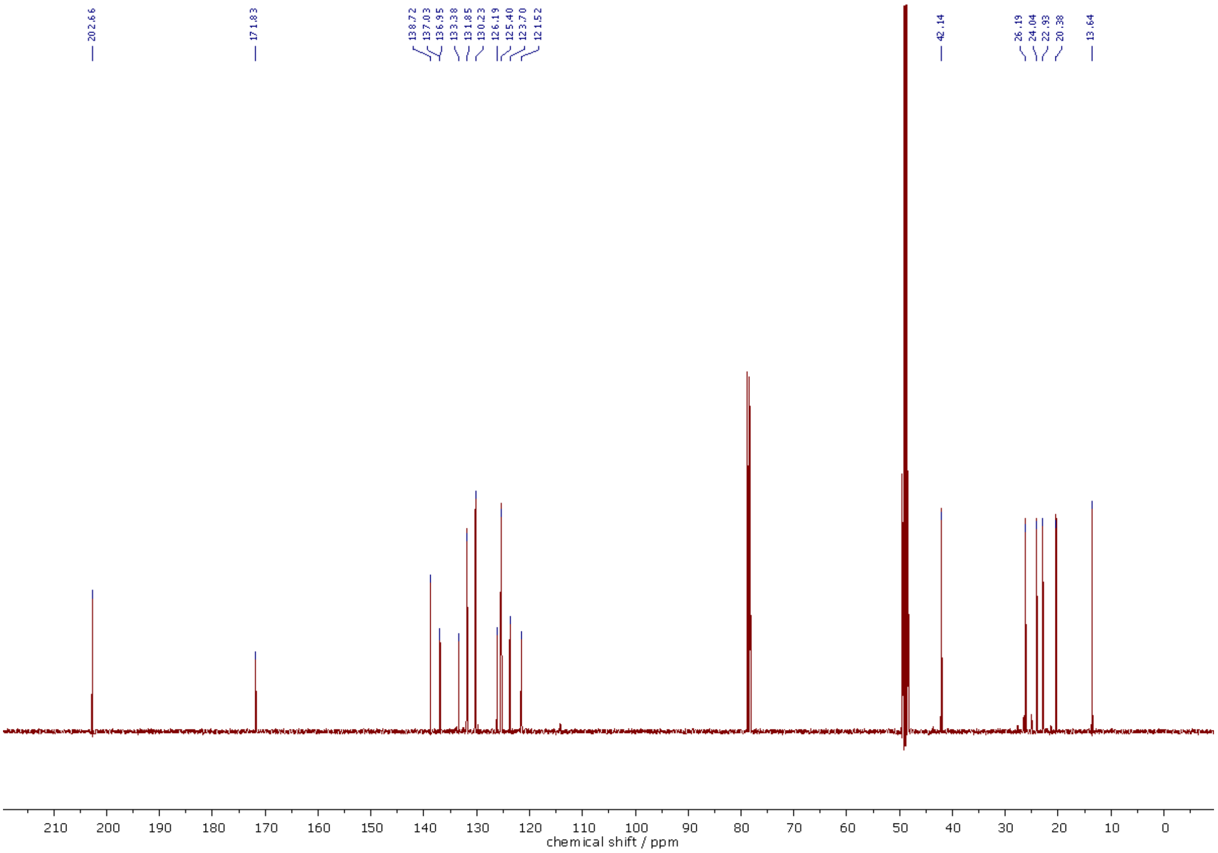

13C NMR (126 MHz, MeOD-$d_4$/$CDCl_3$, 2:1) spectra of **335**.

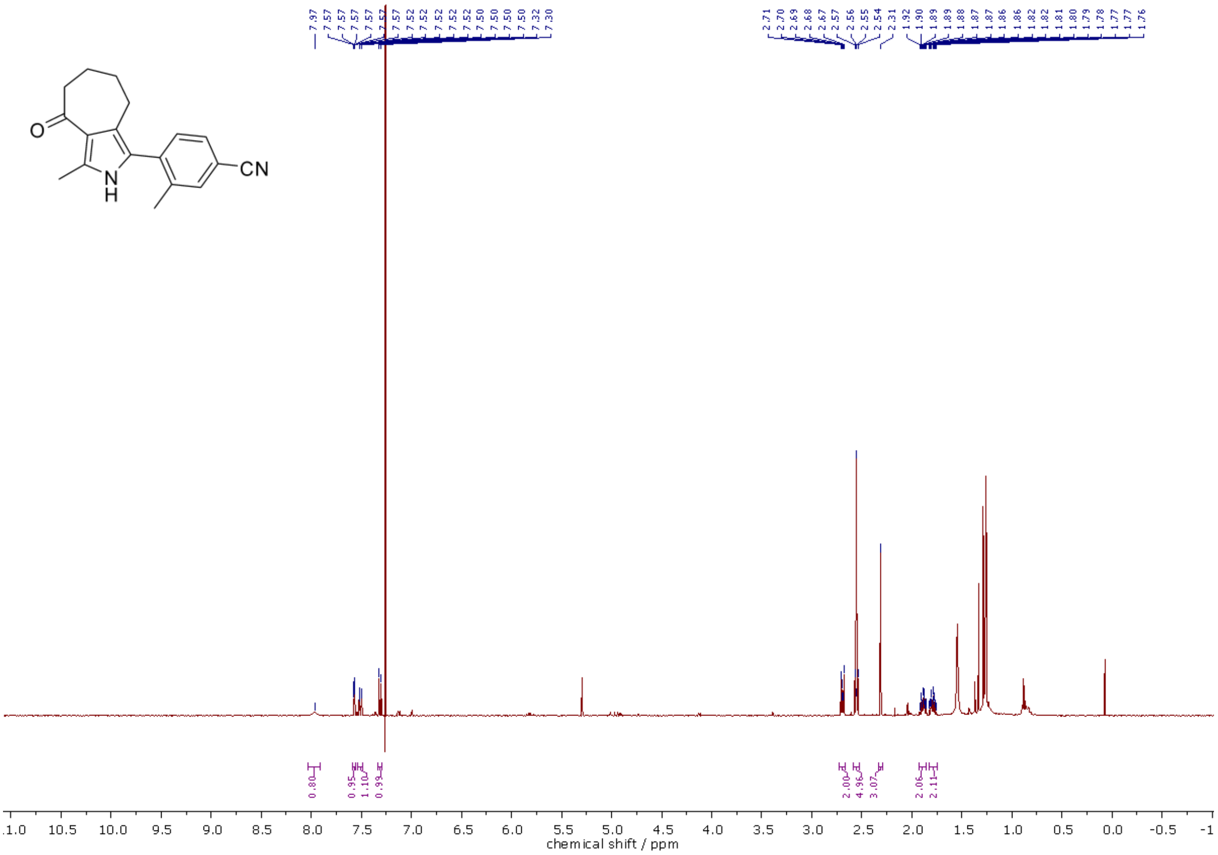

[1]H NMR (400 MHz, $CDCl_3$) spectra of **336**.

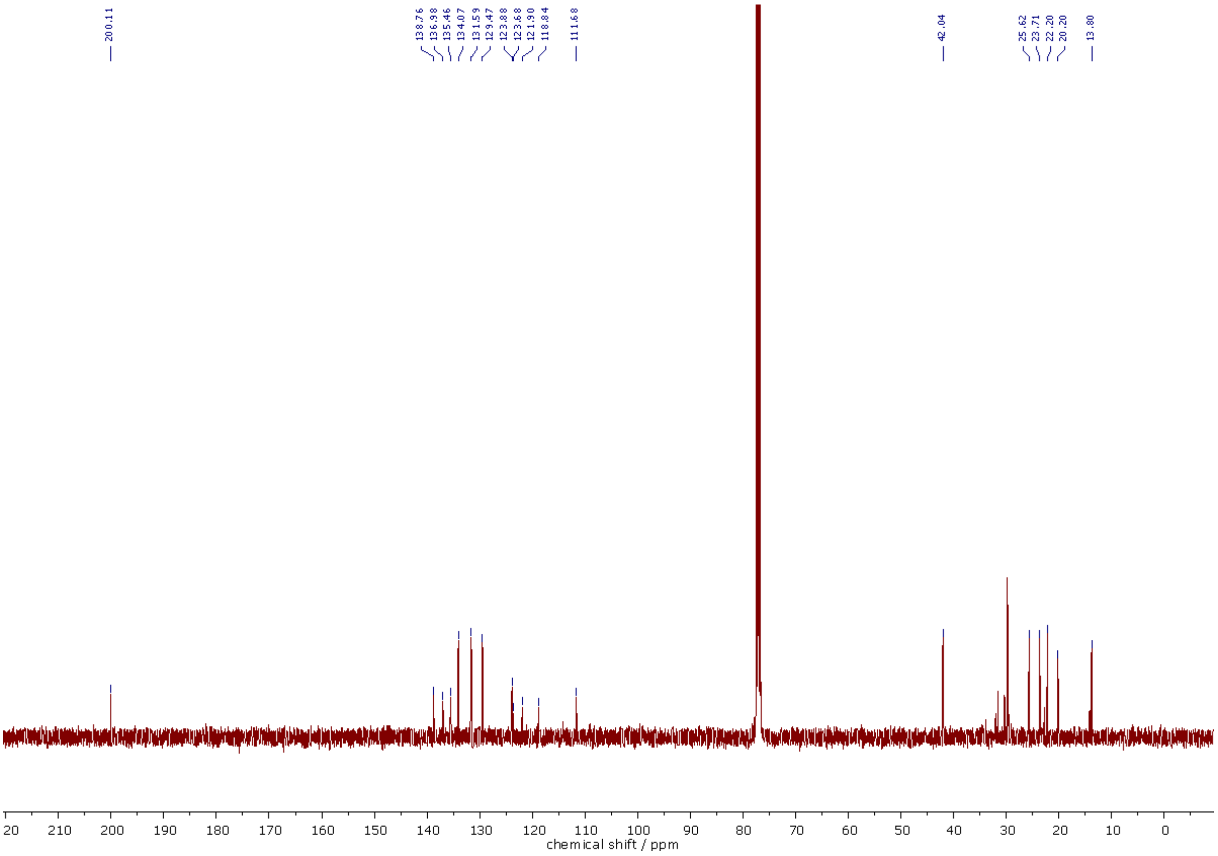

[13]C NMR (101 MHz, $CDCl_3$) spectra of **336**.

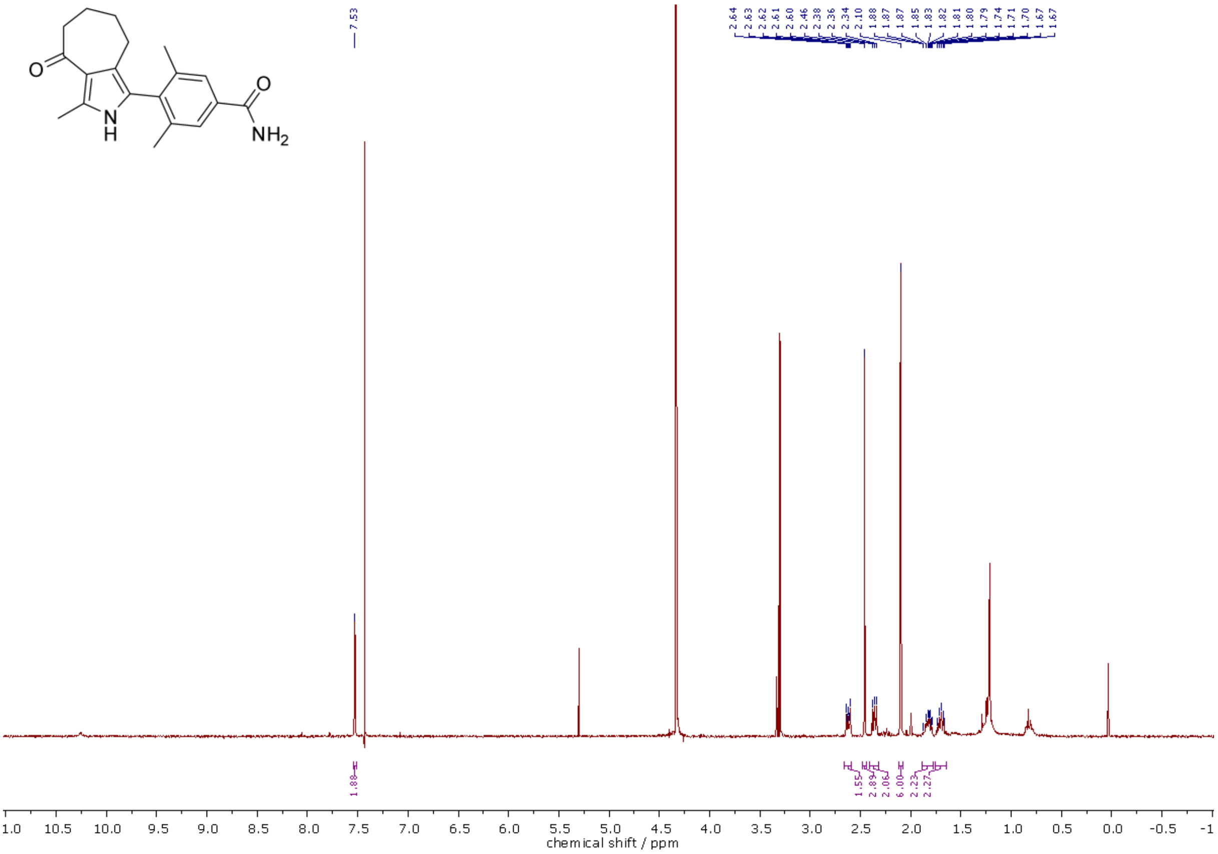

[1]H NMR (300 MHz, $CDCl_3$/MeOD-$d_4$, 2:1) spectra of **355**.

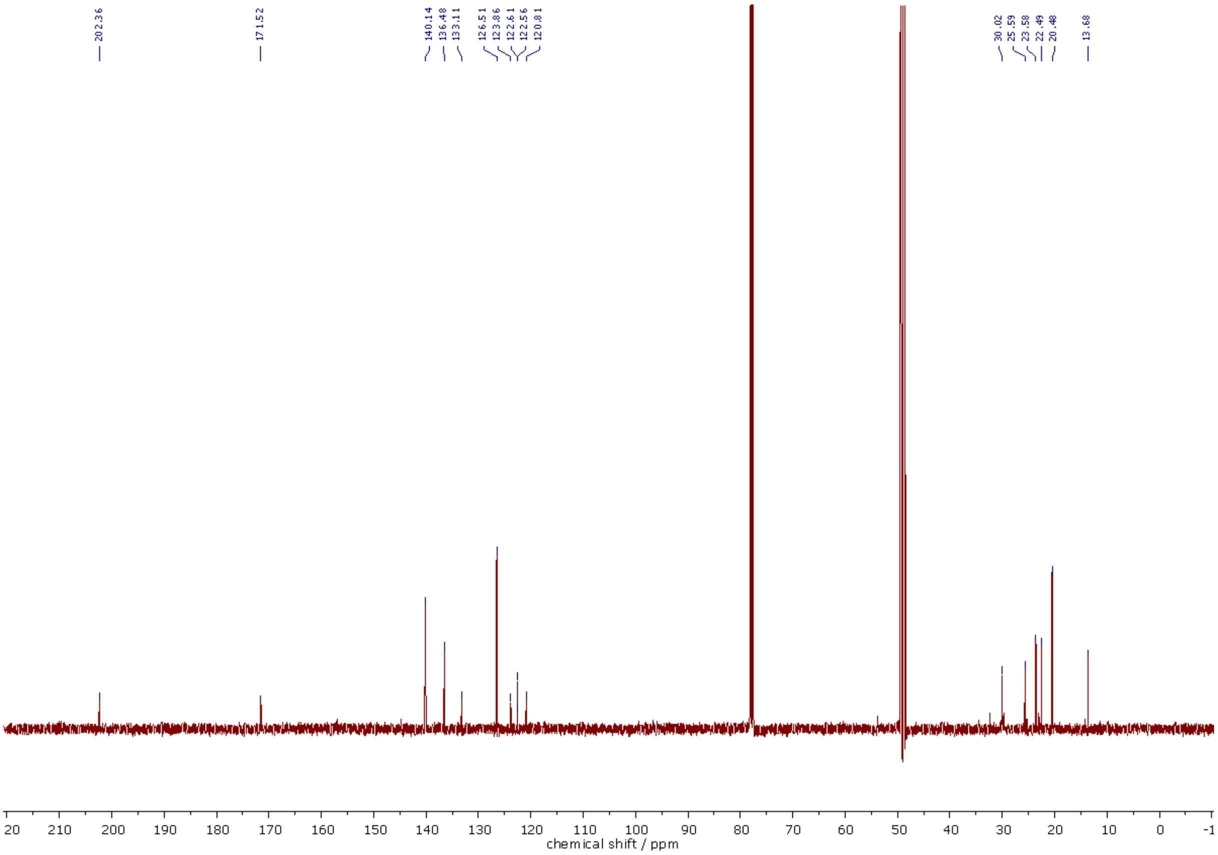

[13]C NMR (126 MHz, $CDCl_3$/MeOD-$d_4$, 2:1) spectra of **355**.

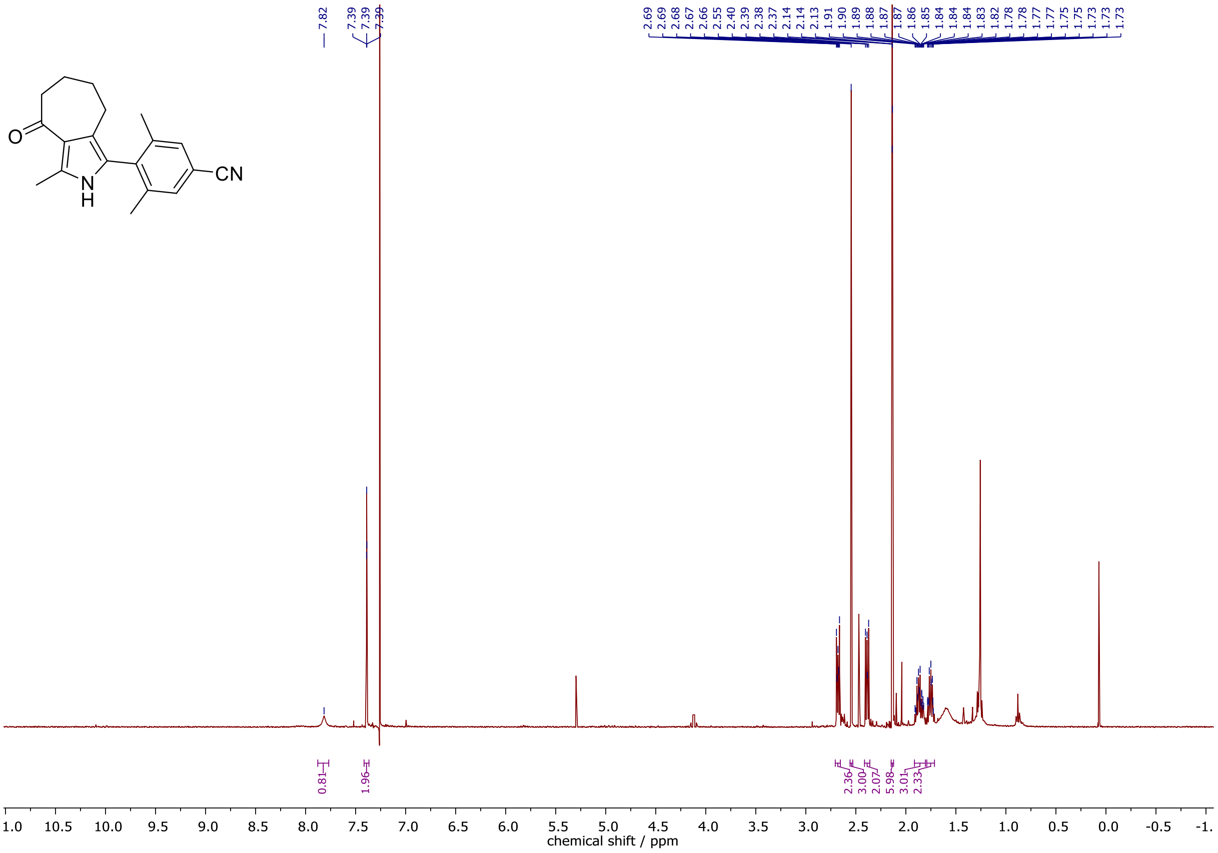

[1]H NMR (400 MHz, $CDCl_3$) spectra of **356**.

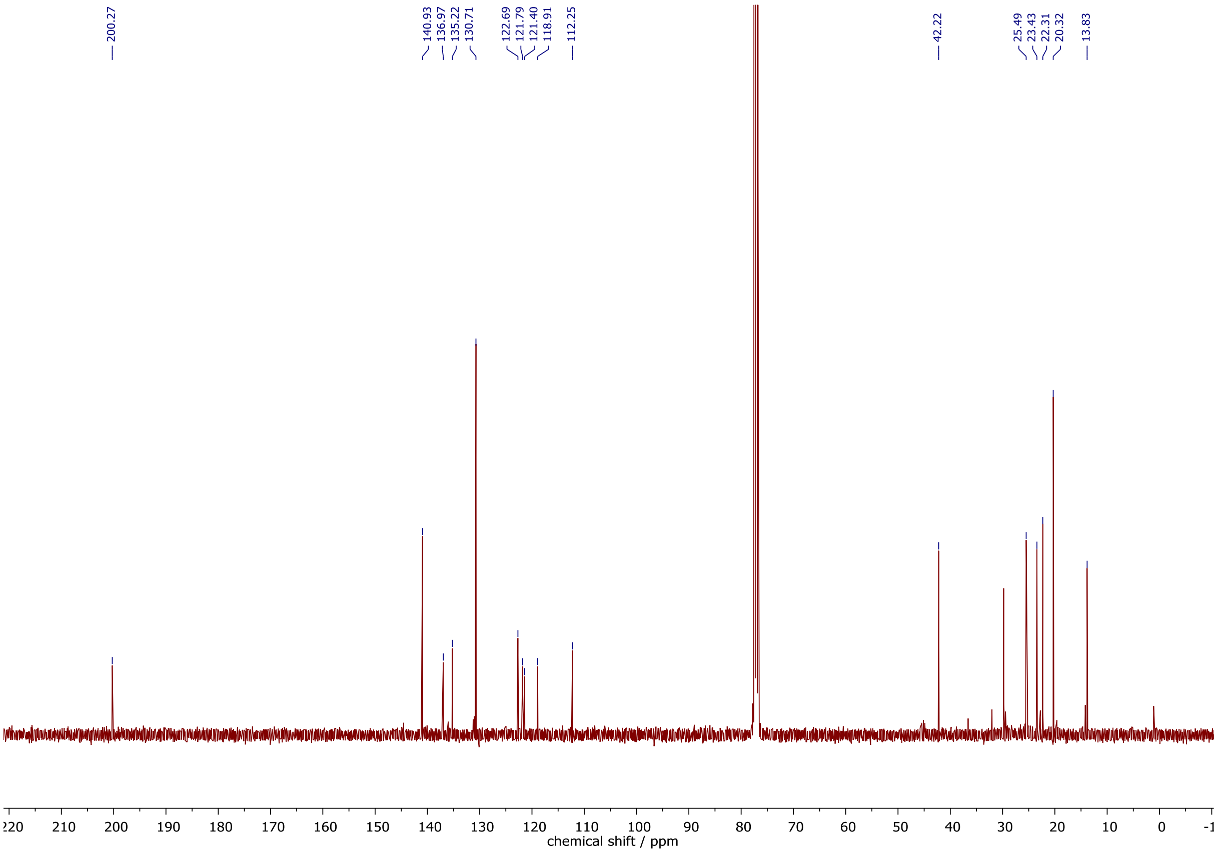

[13]C NMR (101 MHz, $CDCl_3$) spectra of **356**.

**Supporting Table S1.** Amino acid sequences of constructs used in this study

| Name | Protein Sequence |
|---|---|
| PvBD1 (crystal structures) | SFNKQWYLLANQIIQSLSKYEGGHIFEKLVDAKKQNCPDYYDVIKNPMSFSCVKTK<br>LKKGQYGLPTEFIKDVQLIFDNCSLYNTSGSLVAITGKNIEAYFNNQLIVTGYNNFVT<br>KANTINERLQKVEDENLE |
| PfBD1 (ITC and crystal structures) | GSPEFMNFQGNKQWYLLANQLILSLSKYEGGHIFEKLVDAKKQNCPDYYDVIKNP<br>MSFSCIKTKLKKGQYAYPSEFVKDVQLIFDNCSLYNTSNSVVAITGKNIETYFNNQLI<br>VMGYNNFILKEKKINDMLKL |

**Supporting Table S2.** ITC measurements of all compound variations

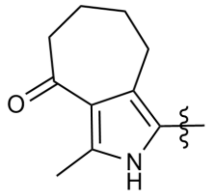

| # | R | $K_D$ [µM] | # | R | $K_D$ [µM] |
|---|---|---|---|---|---|
| MPM3 | 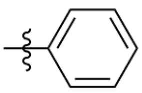 | 10.4 | RMM10 | 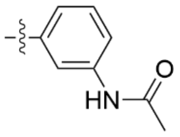 | 6.96 |
| RMM2 | 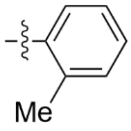 | 3.94 | RMM11 | 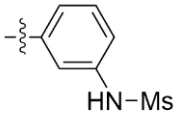 | 4.70 |
| RMM3 | 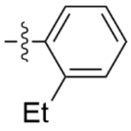 | 9.17 | MPM8 | 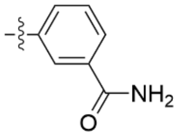 | 7.45 |
| RMM1 | 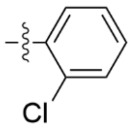 | 5.81 | RMM13 | 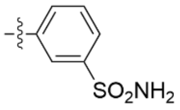 | 4.93 |
| RMM4 | 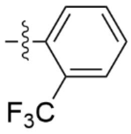 | 2.94 | RMM14 | 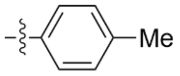 | 11.7 |
| RMM5 | 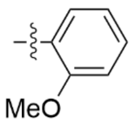 | 7.13 | RMM15 | 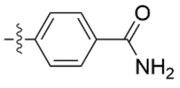 | 4.49 |
| RMM6 | 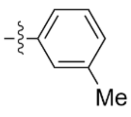 | 6.22 | RMM16 | 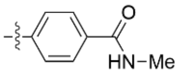 | 5.14 |
| RMM7 | 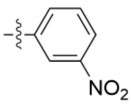 | 16.9 | RMM17 | 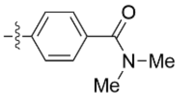 | 42.4 |

| # | R | $K_D$ [µM] | # | R | $K_D$ [µM] |
|---|---|---|---|---|---|
| MPM2 | 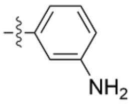 | 6.74 | RMM18 | 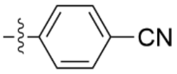 | 4.54 |
| RMM8 | 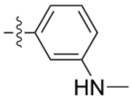 | 9.16 | RMM19 | 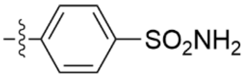 | 6.39 |
| RMM9 | 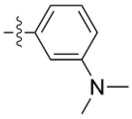 | 8.03 | RMM22 | 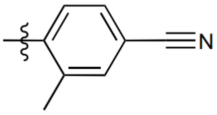 | 2.34 |
| RMM21 | 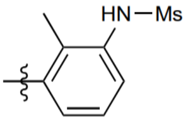 | 2.76 | RMM23 | 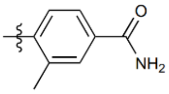 | 1.24 |
| RMM20 | 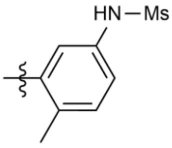 | 8.17 | RMM24 | 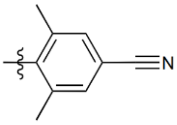 | 1.22 |
| MPM6 | 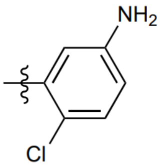 | 5.47 | RMM25 | 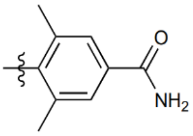 | 0.897 |

#### Supporting Table S3. X-ray data collection and refinement statistics

| | PfBDP1-MPM2 | PfBDP1-RMM2 | PfBDP1-RMM23 | PvBDP1-RMM4 | PvBDP1-RMM21 | PvBDP1-RMM23 | PvBDP1-RMM25 |
|---|---|---|---|---|---|---|---|
| **Wavelength [Ã]** | 1.00000 | 1.000033 | 0.88560 | 1.00000 | 0.999987 | 0.999987 | 0.96670 |
| **Resolution [Ã]** | 43.36 - 1.558 | 44.05 - 2.26 | 35.96 - 1.79 | 46.13 - 1.90 | 46.19 - 2.26 | 46.44 - 1.607 | 53.09 - 1.86 |
| **(high resolution shell)** | (1.68 - 1.56) | (2.59 - 2.26) | (2.05 - 1.79) | (2.05 - 1.90) | (2.59 - 2.26) | (1.68 - 1.61) | (1.98 - 1.86) |
| **Space group** | I 2 2 2 | I 2 2 2 | I 2 2 2 | P 62 2 2 | P 62 2 2 | P 62 2 2 | P 62 2 2 |
| **Unit cell (a, b, c)** | 38.36, 86.72, 107.82 | 38.447, 88.106, 108.046 | 38.12, 88.215, 108.493 | 92.26, 92.26, 72.856 | 92.382, 92.382, 69.811 | 92.883, 92.883, 70.497 | 92.798, 92.798, 70.721 |
| **(α, β, γ)** | 90, 90, 90 | 90, 90, 90 | 90, 90, 90 | 90, 90, 120 | 90, 90, 120 | 90, 90, 120 | 90, 90, 120 |
| **Total reflections** | 180212 (8919) | 117475 (10665) | 59154 (2421) | 582177 (36438) | 333586 (31733) | 47774 (5778) | 31247 (5076) |
| **Unique reflections** | 13888 (695) | 8932 (782) | 9402 (470) | 14932 (2905) | 8689 (778) | 44336 (5491) | 28680 (4779) |
| **Multiplicity** | 13.0 (12.8) | 13.2 (13.6) | 6.3 (5.2) | 38.9 (31.0) | 38.4 (40.8) | 39.1 (37.1) | 37.3 (39.6) |
| **Completeness (%)** | 53.0 (13.1) | 99.7 (97.4) | 52.8 (12.1) | 100.0 (99.3) | 100.0 (100.0) | 99.75 (99.04) | 99.63 (99.57) |
| **Mean I/Ïƒ(I)** | 13.6 (1.4) | 8.7 (1.3) | 13.4 (1.5) | 33.8 (6.1) | 20.0 (5.6) | 26.22 (1.51) | 15.89 (2.26) |
| **Wilson B-factor** | 24.39 | 31.30 | 34.02 | 31.76 | 38.58 | 27.44 | 28.29 |
| **Rmerge** | 0.221 (1.930) | 0.329 (2.676) | 0.070 (0.945) | 0.074 (0.700) | 0.137 (0.965) | 0.063 (3.092) | 0.134 (2.255) |
| **Rmeas** | 0.230 (2.009) | 0.342 (2.781) | 0.076 (1.053) | 0.076 (0.717) | 0.139 (0.977) | 0.064 (3.135) | 0.136 (2.284) |
| **Rpim** | 0.063 (0.550) | 0.094 (0.748) | 0.030 (0.451) | 0.016 (0.157) | 0.023 (0.153) | 0.010 (0.506) | 0.022 (0.361) |
| **CC1/2** | 0.996 (0.622) | 0.995 (0.685) | 0.999 (0.612) | 1.000 (0.978) | 0.999 (0.975) | 1.000 (0.854) | 1.000 (0.912) |
| **Number of non-hydrogen atoms** | 1210 | 1209 | 1175 | 1175 | 1074 | 1142 | 1083 |
| **Macromolecules** | 1042 | 1061 | 1054 | 1003 | 966 | 975 | 968 |
| **Ligands** | 44 | 19 | 44 | 33 | 53 | 33 | 29 |
| **Waters** | 124 | 129 | 77 | 139 | 55 | 134 | 86 |
| **RMS (bonds)** | 0.006 | 0.007 | 0.0021 | 0.003 | 0.002 | 0.015 | 0.013 |
| **RMS (angles)** | 0.76 | 0.80 | 0.40 | 0.48 | 0.39 | 1.15 | 1.07 |
| **Ramachandran favored (%)** | 99.21 | 98.44 | 98.45 | 99.16 | 100.00 | 100.00 | 99.16 |
| **Ramachandran allowed (%)** | 0.79 | 1.56 | 1.55 | 0.84 | 0.00 | 0.00 | 0.84 |
| **Ramachandran outliers (%)** | 0.00 | 0.00 | 0.00 | 0.00 | 0.00 | 0.00 | 0.00 |
| **Average B-factor** | 36.77 | 45.65 | 49.69 | 35.22 | 50.76 | 38.62 | 40.19 |

#### SMILES strings of all compounds presented in the manuscript

| Compound | SMILES |
|---|---|
| MPM2 | c1c(N)cccc1-c([nH]c2C)c(c23)CCCCC3=O |
| MPM3 | c1ccccc1-c([nH]c2C)c(c23)CCCCC3=O |
| MPM6 | c1c(N)ccc(Cl)c1-c([nH]c2C)c(c23)CCCCC3=O |
| RMM1 | Clc1ccccc1-c([nH]c2C)c(c23)CCCCC3=O |
| RMM2 | Cc1ccccc1-c([nH]c2C)c(c23)CCCCC3=O |
| RMM3 | O=C1CCCCc(c12)c([nH]c2C)-c3ccccc3CC |
| RMM4 | O=C1CCCCc(c12)c([nH]c2C)-c3ccccc3C(F)(F)F |
| RMM5 | c1cccc(OC)c1-c([nH]c2C)c(c23)CCCCC3=O |
| RMM6 | c1c(C)cccc1-c([nH]c2C)c(c23)CCCCC3=O |
| RMM7 | [O-][N+](=O)c(c1)cccc1-c([nH]c2C)c(c23)CCCCC3=O |
| RMM8 | CNc(c1)cccc1-c([nH]c2C)c(c23)CCCCC3=O |
| RMM9 | CN(C)c(c1)cccc1-c([nH]c2C)c(c23)CCCCC3=O |
| RMM10 | c1ccc(NC(=O)C)cc1-c([nH]c2C)c(c23)CCCCC3=O |
| RMM11 | c1ccc(NS(=O)(=O)C)cc1-c([nH]c2C)c(c23)CCCCC3=O |
| RMM12 | c1ccc(C(=O)N)cc1-c([nH]c2C)c(c23)CCCCC3=O |
| RMM13 | c1ccc(S(=O)(=O)N)cc1-c([nH]c2C)c(c23)CCCCC3=O |
| RMM14 | c1cc(C)ccc1-c([nH]c2C)c(c23)CCCCC3=O |
| RMM15 | c1cc(C(=O)N)ccc1-c([nH]c2C)c(c23)CCCCC3=O |
| RMM16 | CNC(=O)c(cc1)ccc1-c([nH]c2C)c(c23)CCCCC3=O |
| RMM17 | CN(C)C(=O)c(cc1)ccc1-c([nH]c2C)c(c23)CCCCC3=O |
| RMM18 | c1cc(C#N)ccc1-c([nH]c2C)c(c23)CCCCC3=O |
| RMM19 | c1cc(S(=O)(=O)N)ccc1-c([nH]c2C)c(c23)CCCCC3=O |
| RMM20 | O=C1CCCCc(c12)c([nH]c2C)-c3c(C)ccc(c3)NS(=O)(=O)C |
| RMM21 | c1ccc(NS(=O)(=O)C)c(C)c1-c([nH]c2C)c(c23)CCCCC3=O |
| RMM22 | c1cc(C#N)cc(C)c1-c([nH]c2C)c(c23)CCCCC3=O |
| RMM23 | c1cc(C(=O)N)cc(C)c1-c([nH]c2C)c(c23)CCCCC3=O |
| RMM24 | Cc1cc(C#N)cc(C)c1-c([nH]c2C)c(c23)CCCCC3=O |
| RMM25 | Cc1cc(C(=O)N)cc(C)c1-c([nH]c2C)c(c23)CCCCC3=O |

#### Supporting Figure S1

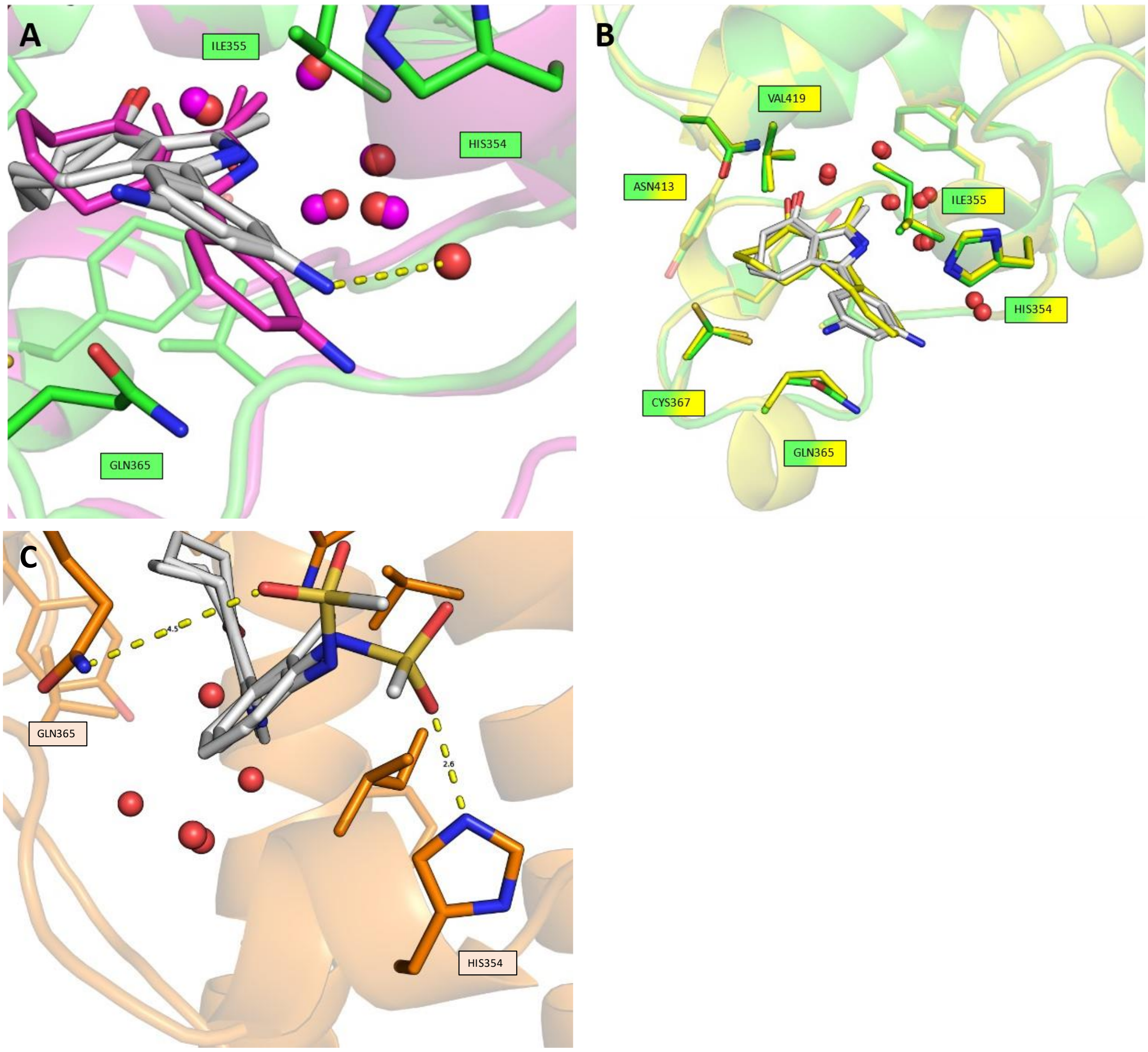

**Supporting Figure S1: A**: BRD4(1) (violet) in complex with MPM2 (violet; PDB ID:7R5B) overlayed with PfBD1 (green) in complex with MPM2 (white). MPM2 shows two conformations in PfBD1 in contrast to BRD4. The conserved water in PfBD1 forms a hydrogen bond with MPM2 while in BRD4 this water has been displaced. **B**: PfBD1 (yellow and green) each in complex with RMM2 (yellow) and MPM2 (white). **C**: PvBD1 (orange) in complex with RMM21. The methanesulfonyl group is present in two conformations, establishing hydrogen interactions with HIS354 and GLN365.

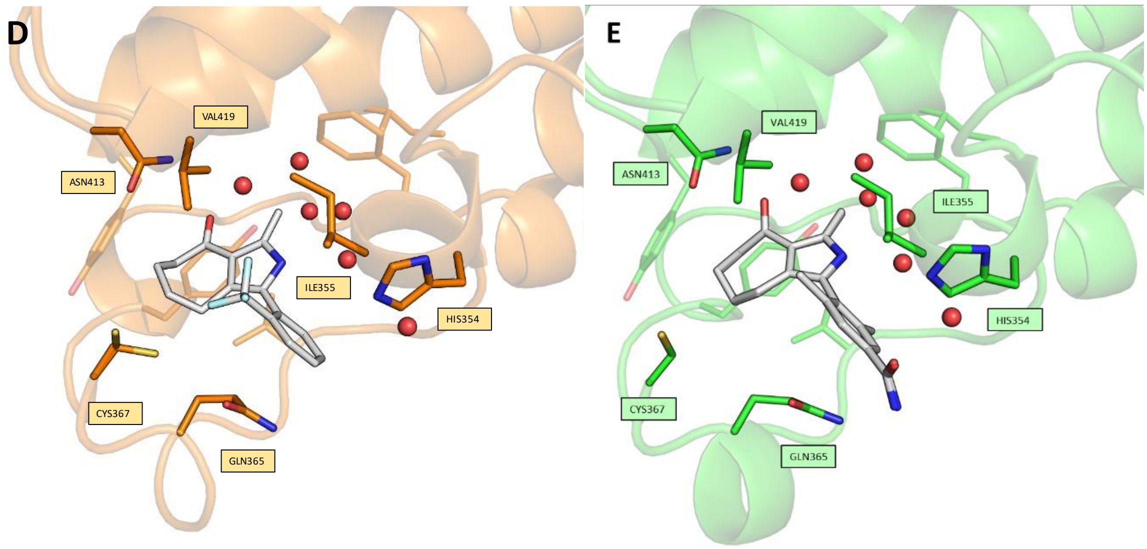

**Supporting Figure S1: D**: PvBD1 (orange) in complex with RMM4. **E**: PfBD1 (green) in complex with RMM23. The BP of the ligand is similar to RMM2 but portion of the ligand is rotated by 180 °.

#### Supporting Figure S2

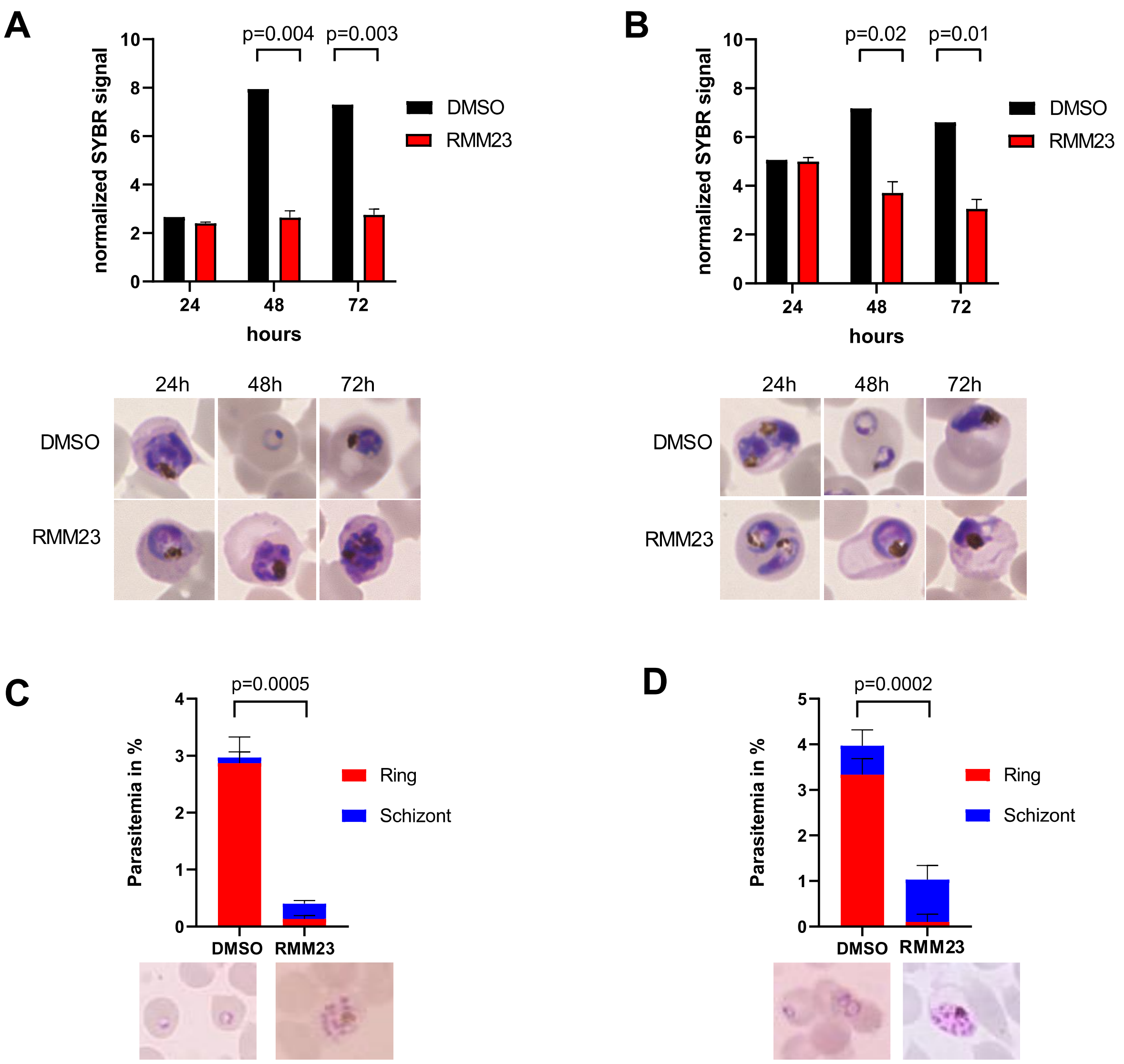

**Figure S2:** RMM23 treatment blocks parasite development at the trophozoite stage and inhibits parasite egress. **A, B.** Treatment with 100 µM RMM23 or DMSO as a control commenced in ring stage cultures of 3D7 (**A**) or NF54 (**B**) parasites. Graphs show normalized SYBR Green signal at 24 h, 48 h and 72 h demonstrating a significant reduction in growth at 48 h (N=3, unpaired t-test). Panels below show Giemsa stained smears of parasites treated for 24 h, 48 h and 72 h with 100 µM RMM23 or DMSO. **C, D.** Treatment commenced in trophozoite stage cultures of 3D7 **(C)** and NF54 **(D)** parasites. Graphs show stage distribution after 24 h treatment with 100 µM RMM23 or DMSO control determined in Giemsa smears (N=3, unpaired t-test). The morphology after 24 h indicates inhibition of parasite egress and reinvasion in RMM23 treated parasites.

#### Supporting Figure S3

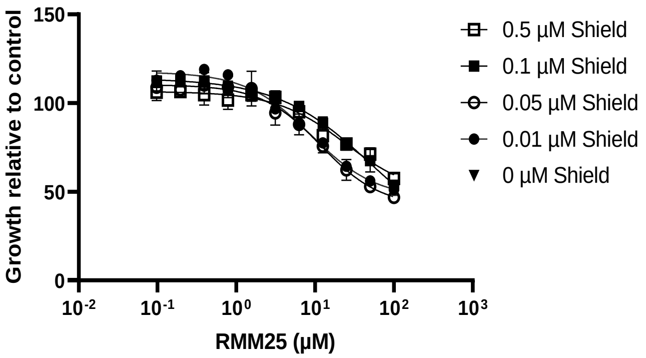

**Figure S3:** Dose-response assay of 3D7::*Pf*BDP1HADD parasites to RMM25. Susceptibility to RMM25 is only mildly affected under conditions of partial *Pf*BDP1 KD, as indicated by a moderate leftward shift in the dose-response curve. Growth was determined relative to DMSO controls for each Shield1 condition after 72 h of culture. N=4 replicates.

#### Supporting Figures ITC

ITC data of the mentioned compounds with PfBD1. Top: Singular Value Decomposition (SVD) reconstructed thermograms. Middle: Fits and derived thermodynamic binding. Bottom: Residuals plots for the fits. Below the figure are the dissociation constant as well as the thermodynamic parameters of the measurement.

MPM6

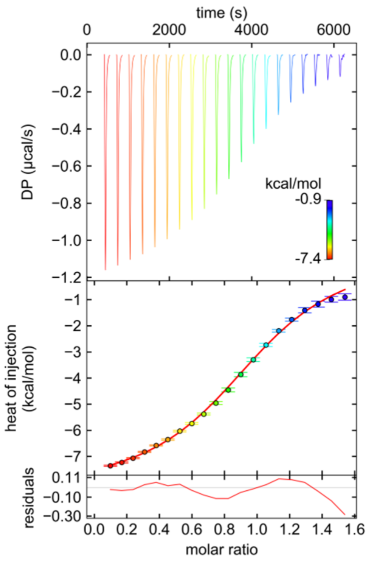

$K_D$ = 5.467 µM
ΔG = -7.179 kcal/mol
ΔH = -9.056 kcal/mol
ΔS = -6.295 cal/mol*K
P99 $_{K_D}$ = 3.895 – 7.896 µM
P99 $_{\Delta H}$ = -10.996 – -7.812 kcal/mol

MPM2

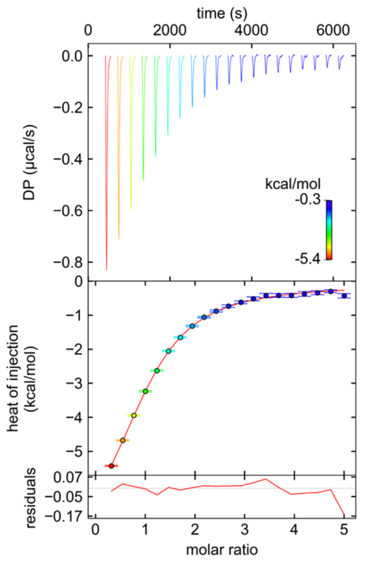

$K_D$ = 6.743 µM
ΔG = -7.055 kcal/mol
ΔH = -8.234 kcal/mol
ΔS = -3.954 cal/mol*K
P99 $_{K_D}$ = 5.524 – 8.343 µM
P99 $_{\Delta H}$ = -9.201 – -7.507 kcal/mol

MPM3

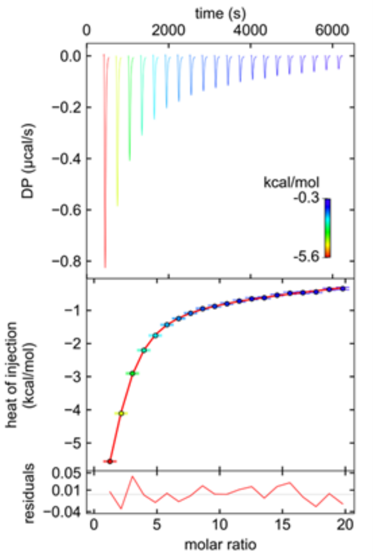

$K_D$ = 10.44 µM
ΔG = -6.796 kcal/mol
ΔH = -8.219 kcal/mol
ΔS = -4.775 cal/mol*K
P99 $_{K_D}$ = 7.333 – 15.952 µM
P99 $_{\Delta H}$ = -9.944 – -7.154 kcal/mol

RMM1

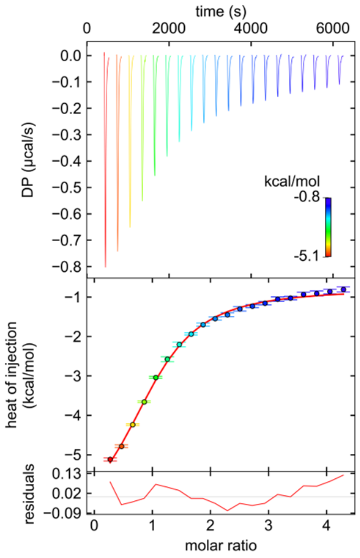

$K_D$ = 5.812 µM
ΔG = -7.143 kcal/mol
ΔH = -6,179 kcal/mol
ΔS = 3.231 cal/mol*K
P99 $_{K_D}$ = 3.743 – 9.513 µM
P99 $_{\Delta H}$ = -7.838 – -5,223 kcal/mol

RMM2

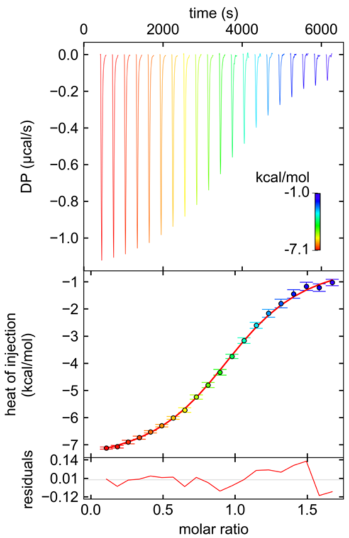

$K_D$ = 3.935 µM
ΔG = -7.374 kcal/mol
ΔH = -7.482 kcal/mol
ΔS = -0.364 cal/mol*K
P99 $_{K_D}$ = 3.079 – 4.954 µM
P99 $_{\Delta H}$ = -8.145 – -6.950 kcal/mol

RMM3

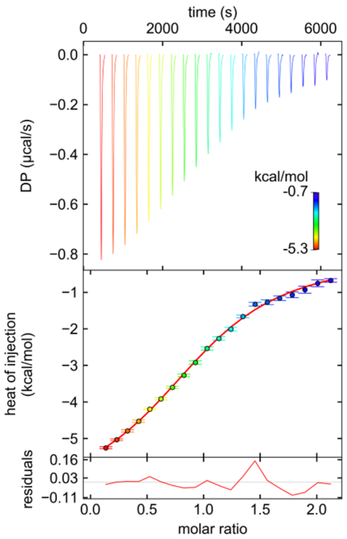

$K_D$ = 9.171 µM
ΔG = -6,873 kcal/mol
ΔH = -6,627 kcal/mol
ΔS = 0.715 cal/mol*K
P99 $_{K_D}$ = 6.371 – 13.955 µM
P99 $_{\Delta H}$ = -7,8533 – -5,8053 kcal/mol

RMM4

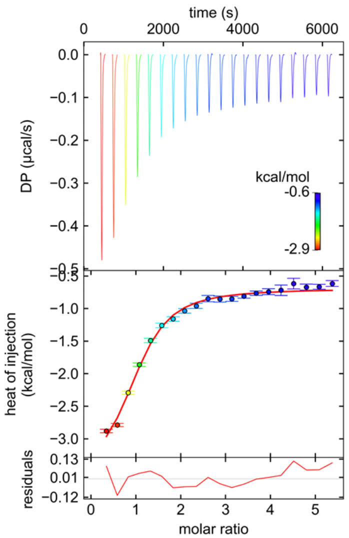

$K_D$ = 2.938 µM
ΔG = -7.547 kcal/mol
ΔH = -2.914 kcal/mol
ΔS = 15.540 cal/mol*K
P99 $_{K_D}$ = 1.126 – 8.733 µM
P99 $_{\Delta H}$ = -8.153 – -7.071 kcal/mol

RMM5

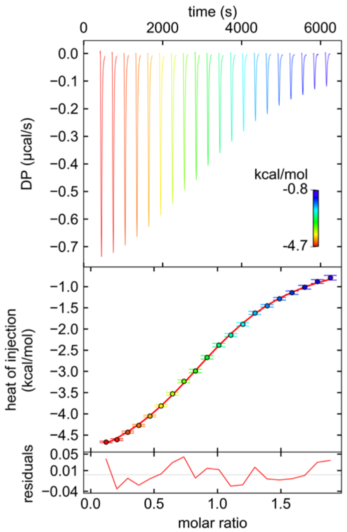

$K_D$ = 7.126 µM
ΔG = -7,022 kcal/mol
ΔH = -5,275 kcal/mol
ΔS = 5.860 cal/mol*K
P99 $_{K_D}$ = 5.466 – 9.538 µM
P99 $_{\Delta H}$ = -5,8910 – -4,8171 kcal/mol

#### RMM6

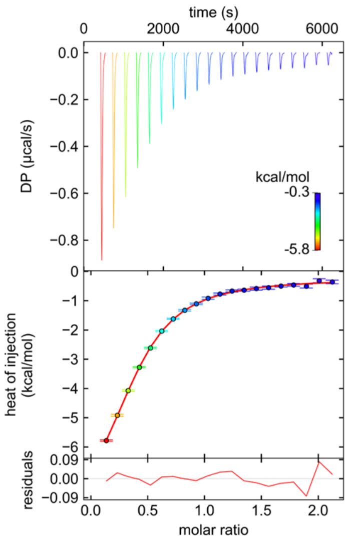

$K_D$ = 6.220 µM
ΔG = -7,103 kcal/mol
ΔH = -8,728 kcal/mol
ΔS = -5,451 cal/mol*K
P99 $_{K_D}$ = 5.228 – 7.478 µM
P99 $_{\Delta H}$ = -8,7278 – -9.6455 kcal/mol

#### RMM7

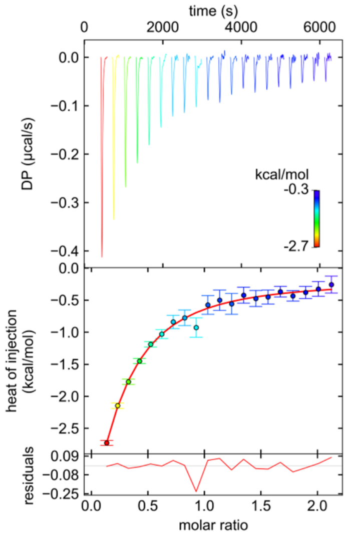

$K_D$ = 16.920 µM
ΔG = -6.510 kcal/mol
ΔH = -12.368 kcal/mol
ΔS = -19.648 cal/mol*K
P99 $_{K_D}$ = 7.104 – 30.828 µM
P99 $_{\Delta H}$ = < -100 – -4,473 kcal/mol

#### RMM8

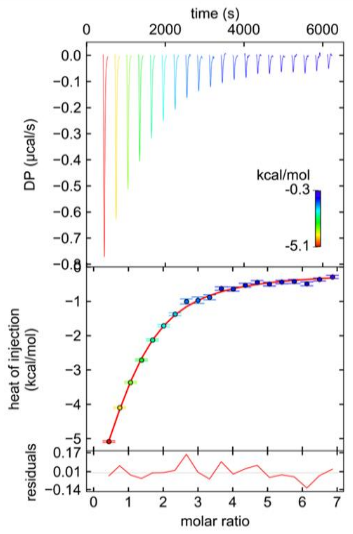

$K_D$ = 9.157 µM
ΔG = -6.873 kcal/mol
ΔH = -10.051 kcal/mol
ΔS = 5.860 cal/mol*K
P99 $_{K_D}$ = 5.857 – 15.983 µM
P99 $_{\Delta H}$ = -17,928 – -7,613 kcal/mol

#### RMM9

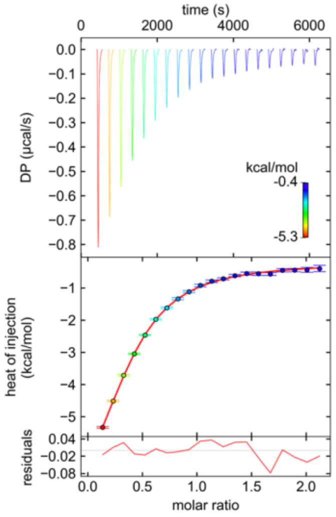

$K_D$ = 8.031 µM
ΔG = -6.951 kcal/mol
ΔH = -8.964 kcal/mol
ΔS = -6,751 cal/mol*K
P99 $_{K_D}$ = 6.440 – 10.238 µM
P99 $_{\Delta H}$ = -10,5726 – -7.8975 kcal/mol

RMM10

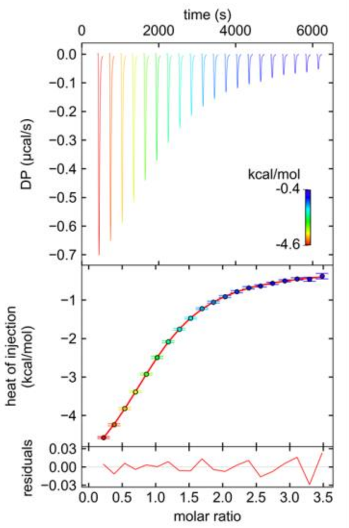

$K_D$ = 6.960 µM
ΔG = -7:036 kcal/mol
ΔH = -6:005 kcal/mol
ΔS = 3:458 cal/mol*K
P99 $_{K_D}$ = 6.341 – 7.662 µM
P99 $_{\Delta H}$ = -6.248 – -5.790 kcal/mol

RMM11

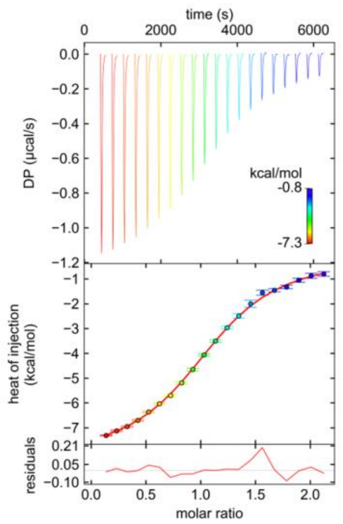

$K_D$ = 4.700 µM
ΔG = -7.269 kcal/mol
ΔH = -8.128 kcal/mol
ΔS = -2.881 cal/mol*K
P99 $_{K_D}$ = 3.848 – 5.795 µM
P99 $_{\Delta H}$ = -8.695 – -7.658 kcal/mol

RMM12

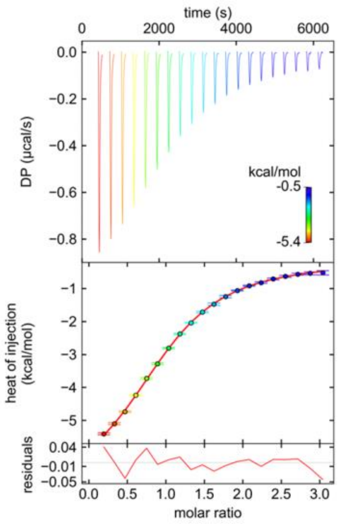

$K_D$ = 7.452 µM
ΔG = -6.996 kcal/mol
ΔH = -7.071 kcal/mol
ΔS = -0.251 cal/mol*K
P99 $_{K_D}$ = 6.253 – 8.959 µM
P99 $_{\Delta H}$ = -7,653 – -6,598 kcal/mol

RMM13

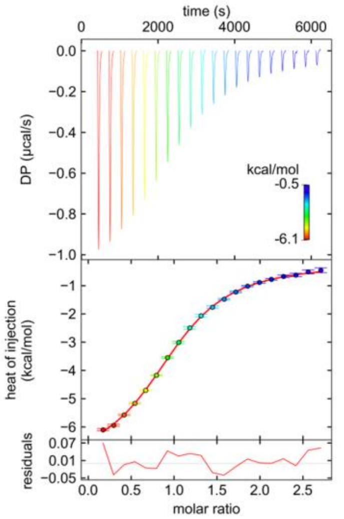

$K_D$ = 4.931 µM
ΔG = -7.240 kcal/mol
ΔH = -7.139 kcal/mol
ΔS = 0.340 cal/mol*K
P99 $_{K_D}$ = 4.020 – 6.097 µM
P99 $_{\Delta H}$ = -7.669 – -6.697 kcal/mol

RMM14

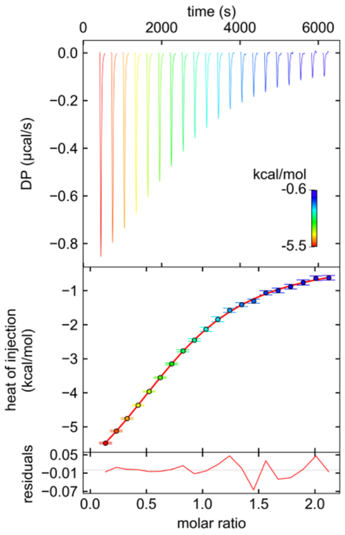

$K_D$ = 11.700 µM
ΔG = -6.728 kcal/mol
ΔH = -7.808 kcal/mol
ΔS = -3.622 cal/mol*K
$P99_{K_D}$ = 9.902 – 14.083 µM
$P99_{\Delta H}$ = -8.480 – -7.282 kcal/mol

RMM15

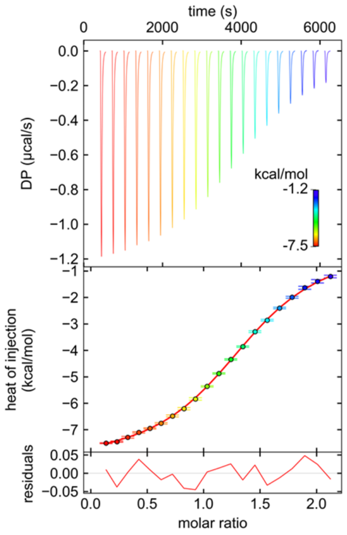

$K_D$ = 4.488 µM
ΔG = -7.296 kcal/mol
ΔH = -8.250 kcal/mol
ΔS = -3.199 cal/mol*K
$P99_{K_D}$ = 3.884 – 5.163 µM
$P99_{\Delta H}$ = -8,723 – -7,812 kcal/mol

RMM16

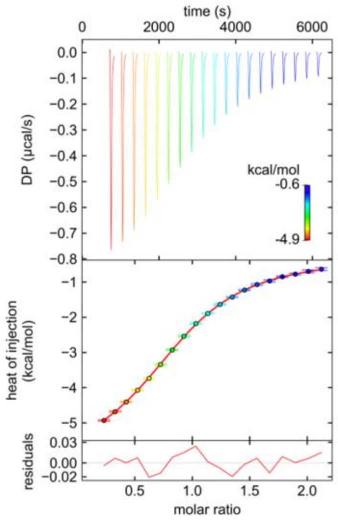

$K_D$ = 5.141 µM
ΔG = -7.216 kcal/mol
ΔH = -5.454 kcal/mol
ΔS = 5.909 cal/mol*K
$P99_{K_D}$ = 3.334 – 8.337 µM
$P99_{\Delta H}$ = -6.435 – -4.805 kcal/mol

RMM17

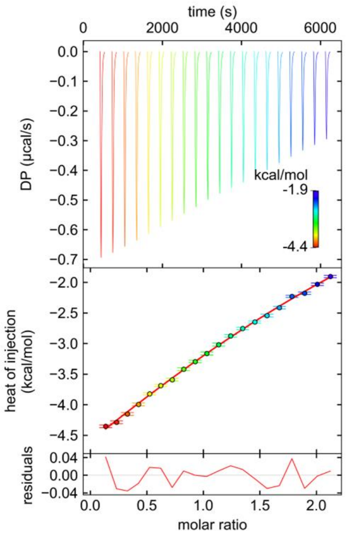

$K_D$ = 42.350 µM
ΔG = -5.966 kcal/mol
ΔH = -7.020 kcal/mol
ΔS = -3.535 cal/mol*K
$P99_{K_D}$ = 16.671 – 100.000 µM
$P99_{\Delta H}$ = -8.480 – -7.282 kcal/mol

#### RMM18

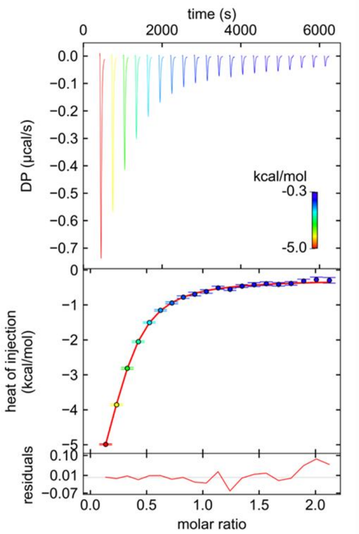

$K_D$ = 4.536 µM
ΔG = -7.290 kcal/mol
ΔH = -8.194 kcal/mol
ΔS = -3.034 cal/mol*K
P99 $_{K_D}$ = 3.786 – 5.490 µM
P99 $_{\Delta H}$ = -9.254– -7.424 kcal/mol

#### RMM19

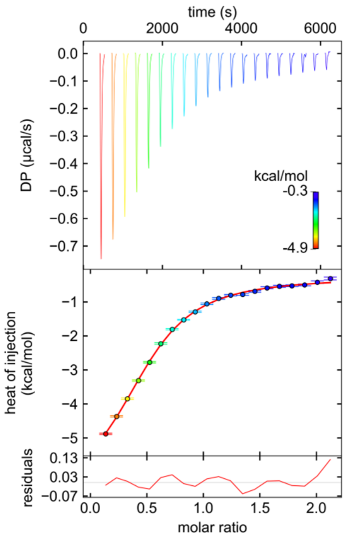

$K_D$ = 6.393 µM
ΔG = -7.086 kcal/mol
ΔH = -6.662 kcal/mol
ΔS = 1.424 cal/mol*K
P99 $_{K_D}$ = 4.297 – 10.081 µM
P99 $_{\Delta H}$ = -8.386 – -5.685 kcal/mol

#### RMM20

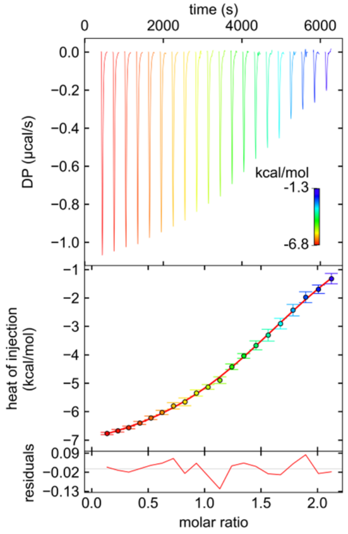

$K_D$ = 8.169 µM
ΔG = -6.941 kcal/mol
ΔH = -9.627 kcal/mol
ΔS = -9.008 cal/mol*K
P99 $_{K_D}$ = 5.981 – 11.542 µM
P99 $_{\Delta H}$ = -13.557 – -7.915 kcal/mol

#### RMM21

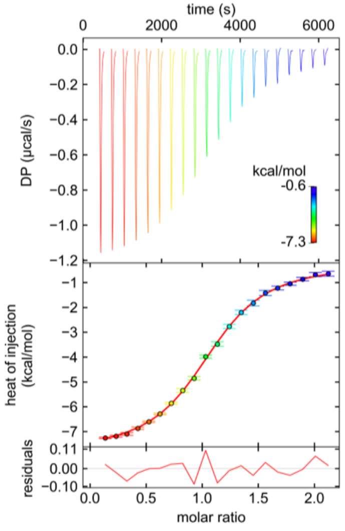

$K_D$ = 2.758 µM
ΔG = -7.584 kcal/mol
ΔH = -7.635 kcal/mol
ΔS = -0.170 cal/mol*K
P99 $_{K_D}$ = 2.284 – 3.334 µM
P99 $_{\Delta H}$ = -8.016 – -7.30 kcal/mol

#### RMM22

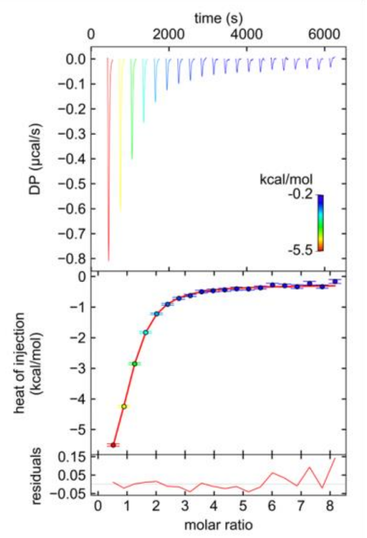

$K_D$ = 2.336 µM
ΔG = -7.683 kcal/mol
ΔH = -7.470 kcal/mol
ΔS = 0.715 cal/mol*K
$P99_{K_D}$ = 1.831 – 3.004 µM
$P99_{\Delta H}$ = -8.415 – -6.775 kcal/mol

#### RMM23

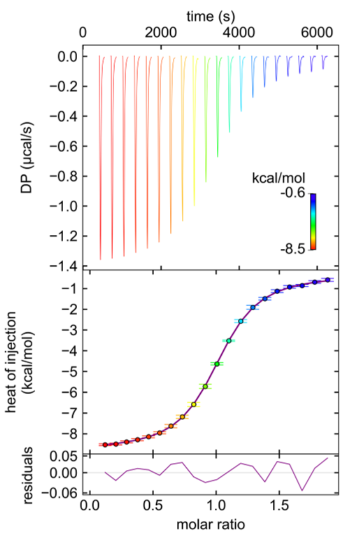

$K_D$ = 1.238 µM
ΔG = -8.059 kcal/mol
ΔH = -8.480 kcal/mol
ΔS = -1.412 cal/mol*K
$P99_{K_D}$ = 1.117 – 1.372 µM
$P99_{\Delta H}$ = -8.617 – -8.350 kcal/mol

#### RMM24

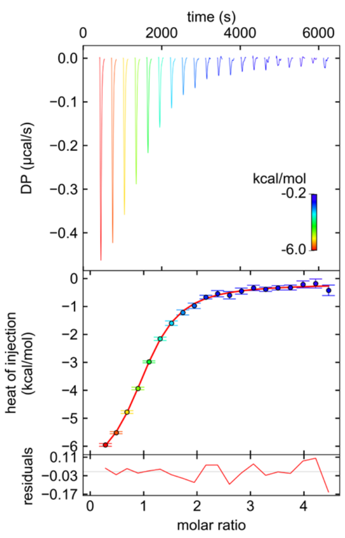

$K_D$ = 1.216 µM
ΔG = -8.070 kcal/mol
ΔH = -6.721 kcal/mol
ΔS = -4.523 cal/mol*K
$P99_{K_D}$ = 0.927 – 1.590 µM
$P99_{\Delta H}$ = -7.296 – -6.252 kcal/mol

#### RMM25

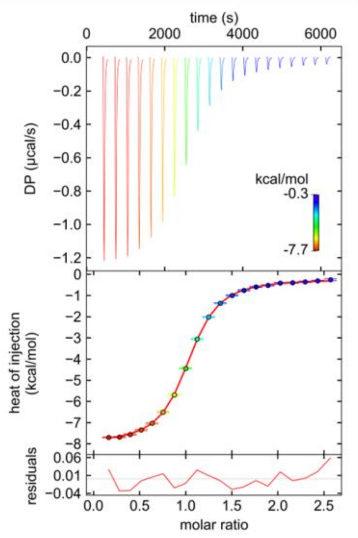

$K_D$ = 0.896 µM
ΔG = -8.250kcal/mol
ΔH = -7.765 kcal/mol
ΔS = 1.628 cal/mol*K
$P99_{K_D}$ = 0.818 – 0.981 µM
$P99_{\Delta H}$ = -7.865 – -7.667 kcal/mol
